# Multivariate methods for testing hypotheses of temporal community dynamics

**DOI:** 10.1101/362822

**Authors:** Hannah L. Buckley, Nicola J. Day, Bradley S. Case, Gavin Lear, Aaron M. Ellison

## Abstract

For ecological research to make important contributions towards understanding and managing temporally-variable global change processes, such as responses to land-use and climatic change, we must have effective and comparable ways to quantify and analyse compositional change over time in biological communities. These changes are the sum of local colonisation and extinction events, or changes in the biomass and relative abundance of taxa within and among samples. We conducted a quantitative review of currently available methods for the analysis of multivariate datasets collected at temporal intervals. This review identified the need for the application of quantitative, hypothesis-based approaches to understand temporal change in community composition, particularly for small datasets with less than 15 temporal replicates. To address this gap, we: (1) conceptually present how temporal patterns in community dynamics can be framed as specific, testable hypotheses; (2) provide three fully-worked case-studies, complete with R code, demonstrating multivariate analysis methods for temporal hypothesis testing and pattern visualisation; and (3) present a road map for testing specific, quantitative hypotheses relating to the underlying mechanisms of temporal community dynamics.

## INTRODUCTION

A key task for ecologists is to monitor, quantify, analyse, and predict temporal changes in the structure of ecological communities as a function of ongoing changes in climate, land use, and other environmental drivers. Ecological communities comprise populations of multiple taxa. The structure of communities—the number of taxa (‘richness’) and their relative abundance (‘composition’)—changes through time because of fluctuations of individual populations and local colonisations and extinctions of taxa (Micheli et al. 1999, Legendre and Gauthier 2014). These temporal dynamics are governed by both intrinsic factors, including intraspecific and interspecific interaction, colonisation, and extinction, and extrinsic factors, which are processes that relate to periodic disturbances and directionally changing environmental conditions (Connell and Slatyer 1977, Brown and Lawson 2010). Understanding the temporal dynamics of community structure can illuminate fundamental ecological processes, including effects of individual life histories on rates of ecosystem change, the relative importance of biotic and abiotic factors in determining community structure, or how taxa and the networks in which they are embedded respond to environmental change (Tilman 1999, Hobbs et al. 2007, Kampichler et al. 2012). Temporal changes in taxon richness and composition also can provide both simple summaries of ecosystem change and information that can be translated rapidly into action, either to protect and restore declining taxa, or to staunch unsustainable growth of unwanted organisms.

Representations of community structure as multivariate datasets that include information on multiple taxa can be more sensitive to changes in explanatory or predictor variables (e.g., pH, temperature, soil type), than univariate summary statistics, such as taxon richness, diversity, or biomass (Micheli et al. 1999, Chao et al. 2014), and hence more informative about causal processes. However, quantitative methods for investigating and testing hypotheses about temporal dynamics of multivariate measures of community structure (henceforth ‘community dynamics’) also are more complex than methods used for univariate data (Schaefer et al. 2005, Legendre and Legendre 2012). To date, methods for the analysis of spatial multivariate community structure data (e.g., Whittaker 1960, Legendre and Legendre 2012) have received far more attention than methods for analysis of temporal multivariate data. Well-known concepts used to describe spatial changes in community structure—grain, extent, nestedness, beta diversity, taxonomic turnover—have been transferred to the temporal domain (Azeria and Kolasa 2008, Anderson et al. 2011, Legendre and Gauthier 2014), and some well-developed multivariate methods for large spatial datasets have been used to detect temporal changes in community structure (Legendre and Gauthier 2014). However, although temporal studies of compositional change are commonly undertaken, it is not always clear which analytical methods are appropriate or available for any given research question, data type, or community structure; indeed, there may not be one ideal method for all situations and a range of methods may be required to answer a given question (Whittaker 1960). Comparisons of multivariate ecological data through time pose unique, non-trivial challenges because these analyses must also account for spatial variation in these data, (Philippi et al. 1998, Micheli et al. 1999, Collins et al. 2000, Schaefer et al. 2005).

In this paper, we: (1) review empirical studies that have undertaken analyses of multivariate temporal community datasets, to gain insight into how researchers have been applying analysis methods over the past several decades, (2) provide recommendations for the greater use of quantitative methods for testing specific temporal hypotheses, and (3) provide guidance for researchers, including those new to community dynamics research, in analysis decisions, particularly for small temporal datasets (fewer than 15 temporal replicates).

### How to use this paper

This paper does not necessarily need to be read from start to finish; we suggest readers identify the section(s) most appropriate for their interests. The remainder of this paper is structured into five main sections: (A) a review of the recent literature; (B) a conceptual description of the types of hypotheses that can be addressed in analyses of temporal community dynamics data; (C) three worked case study examples illustrating hypothesis-driven analyses of temporal community data, including code for the software programme ‘R’ (R Core Development Team 2017) to allow readers to easily implement methods described in this paper; (D) A “road map” to analysing temporal community data, including descriptions, and how to identify and present, analyses appropriate for testing particular hypotheses; (E) A brief discussion of recent advances in multivariate analysis of community dynamics.

## A. LITERATURE REVIEW

### Data collection and analysis

We reviewed and described the multivariate analyses of community dynamics in studies published up until September 2017 to provide an overview of available methods, their benefits and drawbacks, and what they can reveal about community dynamics. Specifically, we asked: Are data collected from different taxonomic groups, habitats, spatial extents, and time-scales analysed in particular (or different) ways, i.e., are there consistencies in analysis approaches due to discipline-specific preferences?

We searched the literature for studies investigating changes in ecological communities over time. We searched the ISI Web of Science Core Collection on 5 September 2017 for articles using the following search statement: [“temporal” AND “composition*” AND “communit*” AND (“community dynamic*” OR“temporal dynamic*” OR“temporal community varia*” OR“community change” OR“compositional change”)]. We refined this search to include only document types listed as ‘articles’ and published in ‘English’. This search yielded 692 articles for review. Of these, we excluded 241 papers that lacked community data (i.e. those that included fewer than three taxa), had no temporal replication or pooled all temporal data, used only artificial communities or simulated datasets, or used space-for-time substitution (e.g., using chronosequences) to infer community ages. Although we did not constrain the temporal search window, our search terms only returned papers after 1990, indicating that the specific terms we used were not part of the compositional change literature prior to the 1990s.

We extracted and collated key attributes relating to each study in the remaining 451 papers from the ISI database (e.g., publication year, number of citations). Specific study attributes were determined from the body text of each manuscript, including information on (i) habitat type (e.g., estuarine); (ii) habitat location (e.g., U.S.A.); (iii) taxonomic identity (e.g., vertebrate; fish); (iv) manipulation (e.g., natural variation, experimental resource manipulation); (v) spatial extent (e.g., local, global); (vi) temporal grain and extent (e.g., minutes, months); (vii) how composition was measured (e.g., presence-absence, relative abundances of taxa); (viii) how time was included in each analysis (i.e., explicitly as a predictor variable, or where other variables represent time, e.g., climatic factors recorded at different times); and (ix) the key perceived research question (e.g., exploration of yearly dynamics) (Data S1). We categorised organisms being studied into the following taxonomic groups: plants (Kingdom Plantae); vertebrates (Phylum Chordata); invertebrates (Phyla Annelida, Arthropoda, Cnidaria, Ctenophora, Echinodermata, Mollusca, Platyhelminthes, or Porifera); microeukaryotes (algae and Kingdoms Fungi and Protista); prokaryotes (Domains Bacteria and Archaea); and viruses (non-cellular pathogenic organisms). The data types collated and included papers are provided in the supplementary material (Data S1).

### Literature Review Results and Discussion

The 451 papers identified by our search, when compared by year to the number of papers in community ecology, showed that every year since 1990, publications on temporal community dynamics consistently have comprised approximately 24% of studies within community ecology. Thus, studies on temporal dynamics have increased from less than five publications per year in the early 1990s to more than 30 per year in the last five years (Appendix S1: Fig. S1). Few of the 451 reviewed publications (20 papers; 4% of those reviewed) contained more than one independent dataset; those instances were treated as multiple studies that were analysed as separate data points (increasing our total sample size *n* to 475). The majority of studies (70%) were conducted within North America and Europe (Fig. 1), likely reflecting the relative availability of research funding and resources.

**Fig. 1.**
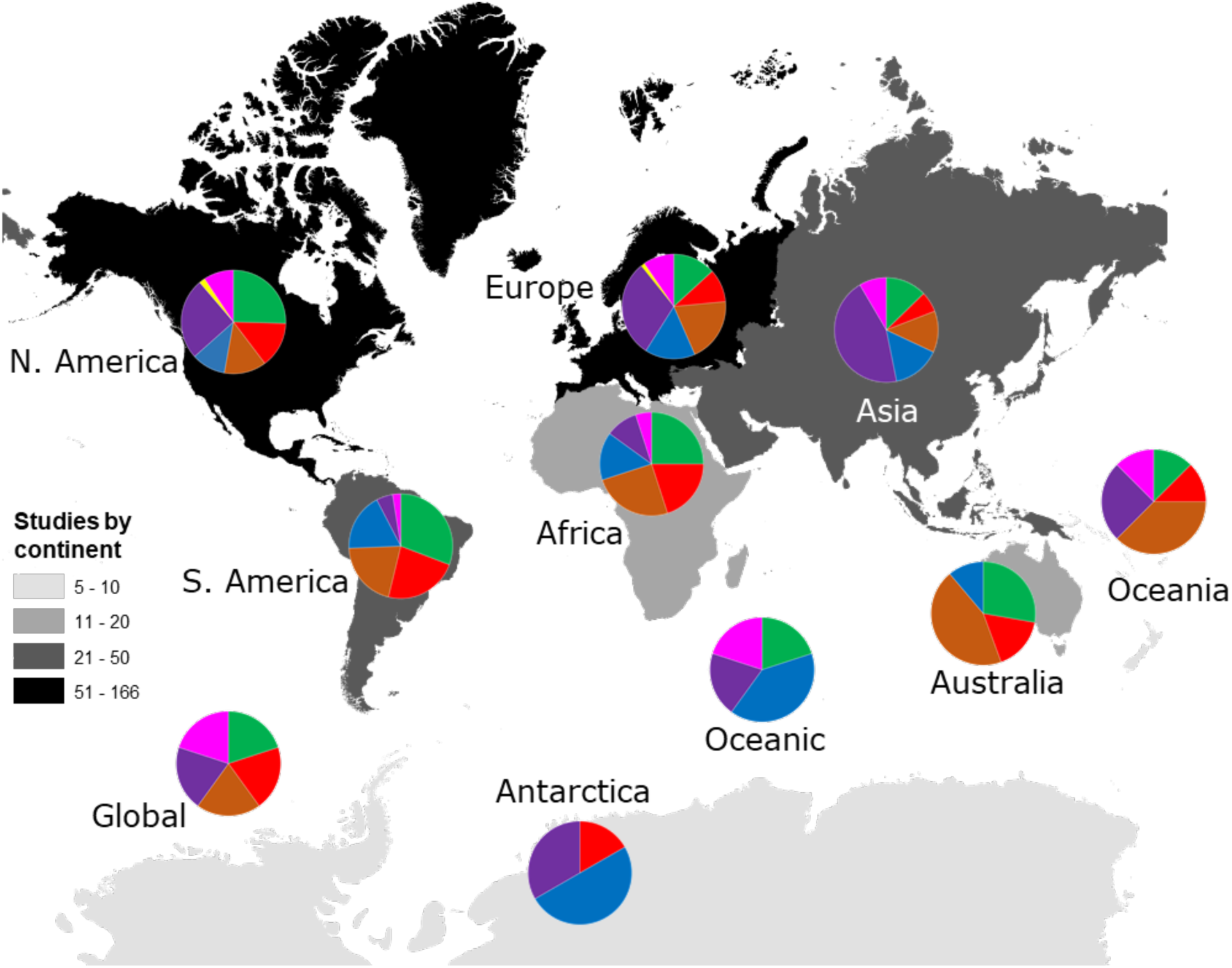
Number of studies identified by our search criteria from each continent. Numbers of studies were: Global (*n* = 5), Oceanic (*n* = 5), Africa (*n* = 20), Antarctica (*n* = 6), Asia (*n* = 47), Australia (*n* = 18), Europe (*n* = 166), North America (*n* = 161), Oceania (*n* = 8) and South America (*n* = 39). Darker colours indicate more studies have been conducted within those continents, including Oceania. Pie charts show the taxonomic focus of study data from each continent as being on (•) plants, (•) vertebrate animals, (•) invertebrate animals, (•) microeukaryotes, (•) prokaryotes, (•) viruses, or (•) mixed taxonomic groups.

Reviewed studies generally focused on organisms, scales, and habitats that were conducive to studying temporal community dynamics. For instance, prokaryotes and invertebrates were studied most frequently (Fig. 1), being relatively dynamic and easy to study over smaller temporal and spatial scales; by contrast, research on plants and vertebrate fauna together accounted for less than one-third of all studies of community dynamics. Although studies spanned a broad range of habitat types, taxa, spatial, and temporal scales, there were two-to-three times more studies at small spatial extents (micro, point sample, and local scales) across the 14 habitat categories identified from the articles (Appendix S1: Fig. S2A). Similarly, the most common temporal extent (i.e., the time between the collection of the first and last samples) of individual studies was years, and only a minority of studies on plant and vertebrate communities extended for a decade or more (Appendix S1: Fig. S2B). Smaller organisms were often measured at finer temporal resolutions (grain) than larger organisms, and many studies on short-lived prokaryote and viral communities lasted more than one year. The temporal grain of sampling generally was smaller for communities of microorganisms than for communities of vertebrate animals and plants, for which the most common grain for repeated sample collection was years rather than months or seasons (Appendix S1: Fig. S2B).

Researchers frequently used a range of different analytical methods, even within one study, to enable them to understand the complexities of observed temporal changes. Up to eight different analytical methods were used (Fig. 2A), with about two-thirds of the studies applying ≤ 3 methods, and most using two. Although the number of studies addressing community dynamics have increased dramatically since the 1990s and have begun to incorporate more computationally-complex methods over time (Appendix S1: Fig. S3A), descriptive methods (i.e., visual, non-statistical comparisons) and ordination have remained the most commonly-used analyses, comprising 33% and 22% of all applied methods, respectively (Fig. 2B). In fact, 134 out of 451 studies (30%) used only descriptive approaches to “analyse” their datasets, while another 35% of studies used descriptive methods in addition to at least one other method. We suspect that the overall ease of use of descriptive and ordination methods, their accessibility via readily-available software programs, and their focus on visually-presenting complex multivariate data, together have driven this trend. The use of the raw dissimilarity values (18%), time-lag analysis (9%), and cluster analysis (7%) comprised the next most common types of methods; the remaining 12 methods were used infrequently (Fig. 2B). Several methods captured by this review were not multivariate (see Appendix 2), including univariate linear methods, temporal stability (*a.k.a*. coefficient of variation), frequency change (testing for significant changes in the frequency of individual taxa over time), and multiplicative change (percent cover change statistic for a single taxon or taxon group used in subsequent regression analyses). Since 2005, the use of raw dissimilarity methods has increased considerably. In the same decade, a steady stream of alternative methods that incorporate time explicitly have been applied, albeit with poor uptake across the wider literature in this area. These methods often are ‘one-off’ analyses that have been applied only a handful of times; we refer to these as ‘minor’ methods (Appendix S1: Fig. S3). Preferred ordination methods changed from detrended correspondence analysis (DCA) and principal components analysis (PCA) in the early 1990s, to canonical correspondence analysis (CCA) in the late 1990s, and, since the early 2000s, to non-metric multidimensional scaling (NMDS) (Appendix S1: Fig. S3B).

**Fig. 2.**
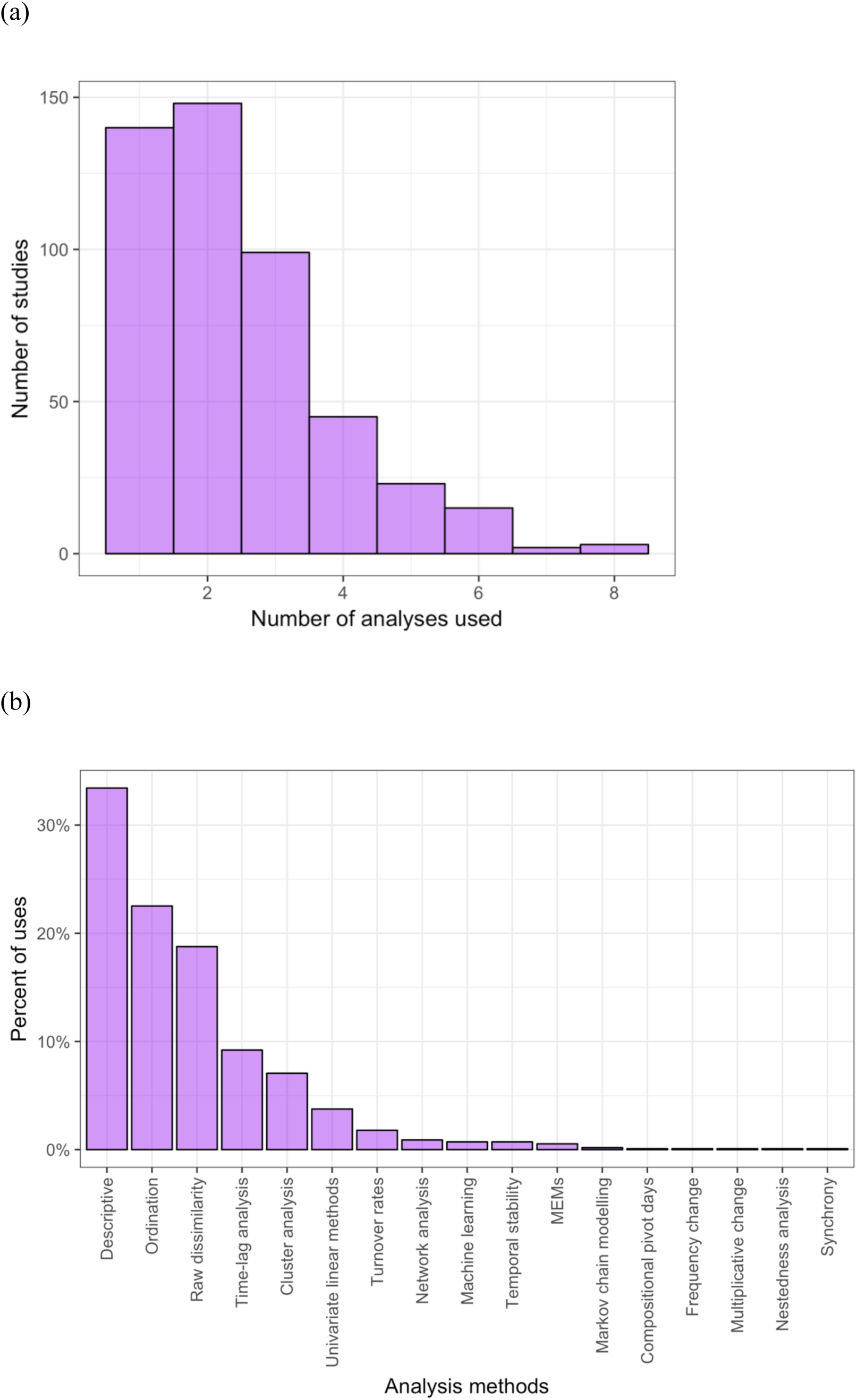
The frequency of use of different temporal community dynamics analysis methods across all reviewed studies. Specifically: (a) the number of different analyses applied in each study; (b) the percent of uses of each analysis type across all studies and times. Descriptions of analysis types are detailed in either S1 or S2 Text.

Three-quarters of all analyses compiled were from observational studies aimed at describing patterns of community change or related to monitoring community composition, rather than exploring environmental effects or studying the effects of biotic processes or interactions (Appendix S1: Fig. S4). In fewer cases, researchers were interested in the effect of environmental change, such as seasonality or climatic cycles over longer time periods.

Our review also shows how common it is for researchers to analyse datasets with low temporal replication. Across research aims and for most methods, most time series had fewer than 15 observations (mean = 14.9, median = 7, mode = 4; Fig. 3); the few studies that used network analyses and machine-learning methods used 50 or more observations taken through time. Overall, neither the number of temporal replicates, nor any other study attributes we extracted during our review (e.g., habitat, taxonomic group), appeared to explain well the differences in how researchers analysed their data or their motivations for doing so.

**Fig. 3:**
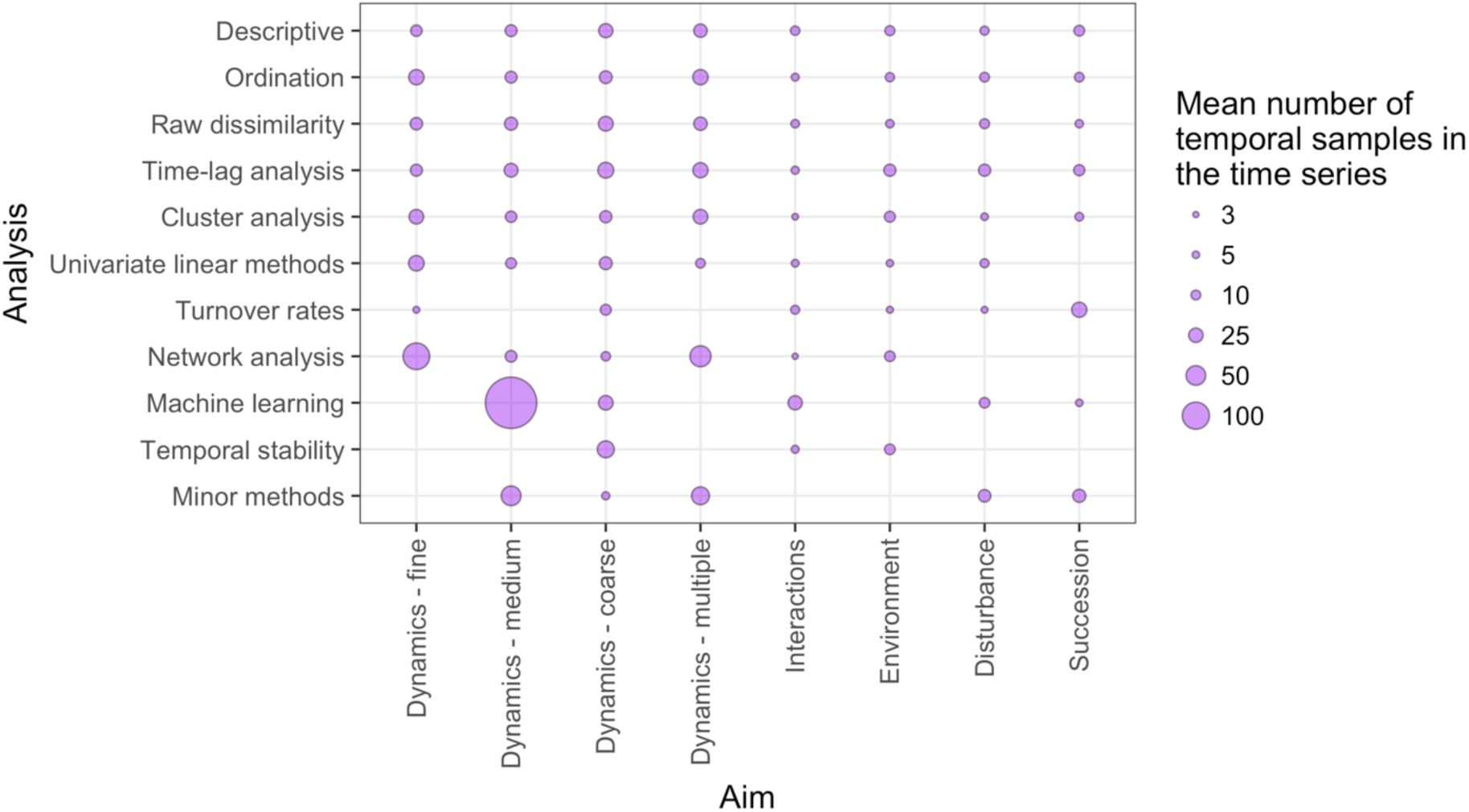
The relative distribution of the mean number of independent temporal samples across different research aims and temporal analysis methods. ‘Minor methods’ include compositional pivot days, frequency change, Markov chain modelling, multiplicative change, Moran’s eigenvector maps (MEMs), nestedness analysis, and synchrony. Descriptions of analysis types are detailed in Section D ‘Road map for analysing temporal community dynamics’ in the text. Research aims vary from understanding compositional dynamics at fine, medium and coarse temporal scales, to investigating the effect on species composition over time of species interactions, environmental conditions, and ecological disturbance, to understanding successional change in communities.

Typically, researchers will have some notion of how their communities of interest might be changing, which motivated them to undertake temporal sampling in the first place (Bestelmeyer et al. 2011). In some cases, long-term datasets exist because previous researchers began data collection with a rationale and method that is different from the current research questions, and so we can sometimes be limited by past sampling designs. Nonetheless, our review highlights that most studies describe temporal community dynamics within their studied communities without explicit, quantitative hypotheses and specific, testable predictions about the expected drivers of observed community changes (Appendix S1: Fig. S4). Clearly, there are many benefits to entering the analysis pathway with a strong *a priori* prediction of the compositional pattern(s) expected and the biological hypotheses that underpin those predictions.

## B. HYPOTHESES OF TEMPORAL COMMUNITY DYNAMICS

Quantitative and specific predictions of the nature and variability in temporal community dynamics allows deeper understanding of the patterns and processes involved. However, to achieve this, *a priori* expectations of change need to be developed and paired with the appropriate analytical method(s) to achieve rigorous tests. Here, we describe in detail the possible observed changes in community composition (taxon presence or abundance) in one sampling unit (site, host, etc.) over time (Fig. 4) or in a set of sampling units over time (Figs. 5-7) and what some of the ecological research questions are that may underpin such hypotheses. In each case, we give predictions for how pairwise dissimilarity values, the basis for most of the commonly-used quantitative measures of compositional change identified in the literature review, would change. Many of these hypotheses are not mutually exclusive although some could be used as multiple alternatives to be tested simultaneously, depending on the research question(s) of interest. Note that ‘dissimilarity measures’, also called ‘distance measures’ or ‘beta-diversity’, are the inverse of ‘similarity measures’. These metrics quantify the compositional distance between two samples (pairwise dissimilarity).

**Fig. 4.**
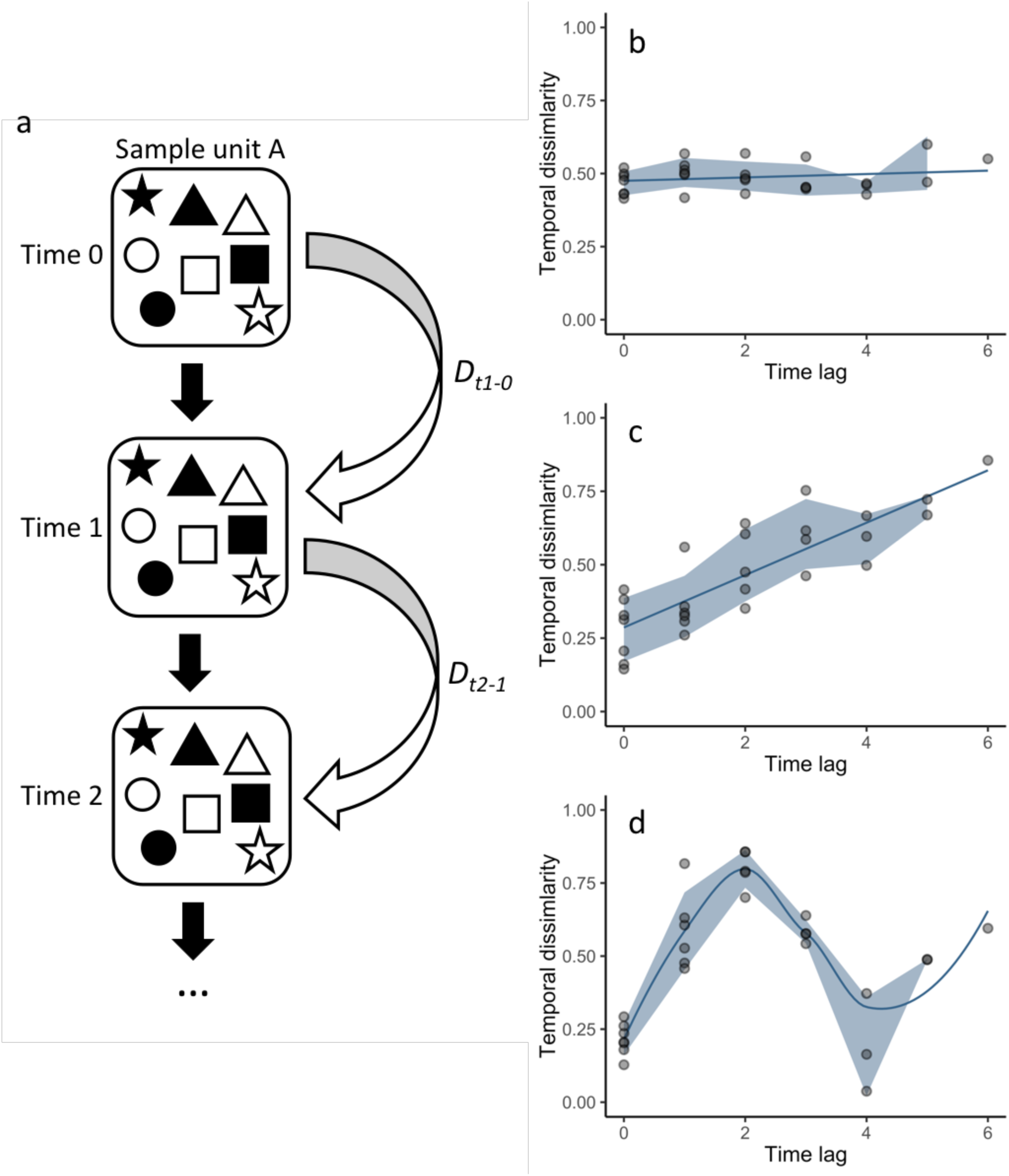
Conceptual figure illustrating the different patterns in (a) temporal community dynamics for a single sample unit expected under different hypotheses of community change, visualised as how pairwise dissimilarities (*D*), e.g., Bray-Curtis, would change over different time lags: (b) the null hypothesis of no change (‘persistence’), (c) ‘directional’ change, and (d) ‘cyclical change’/ ‘periodicity’. Points represent individual pairwise comparisons of across eight sampling occasions and the error envelope is the standard deviation (absent for the largest lag because there is only one comparison at that scale).

**Fig. 5.**
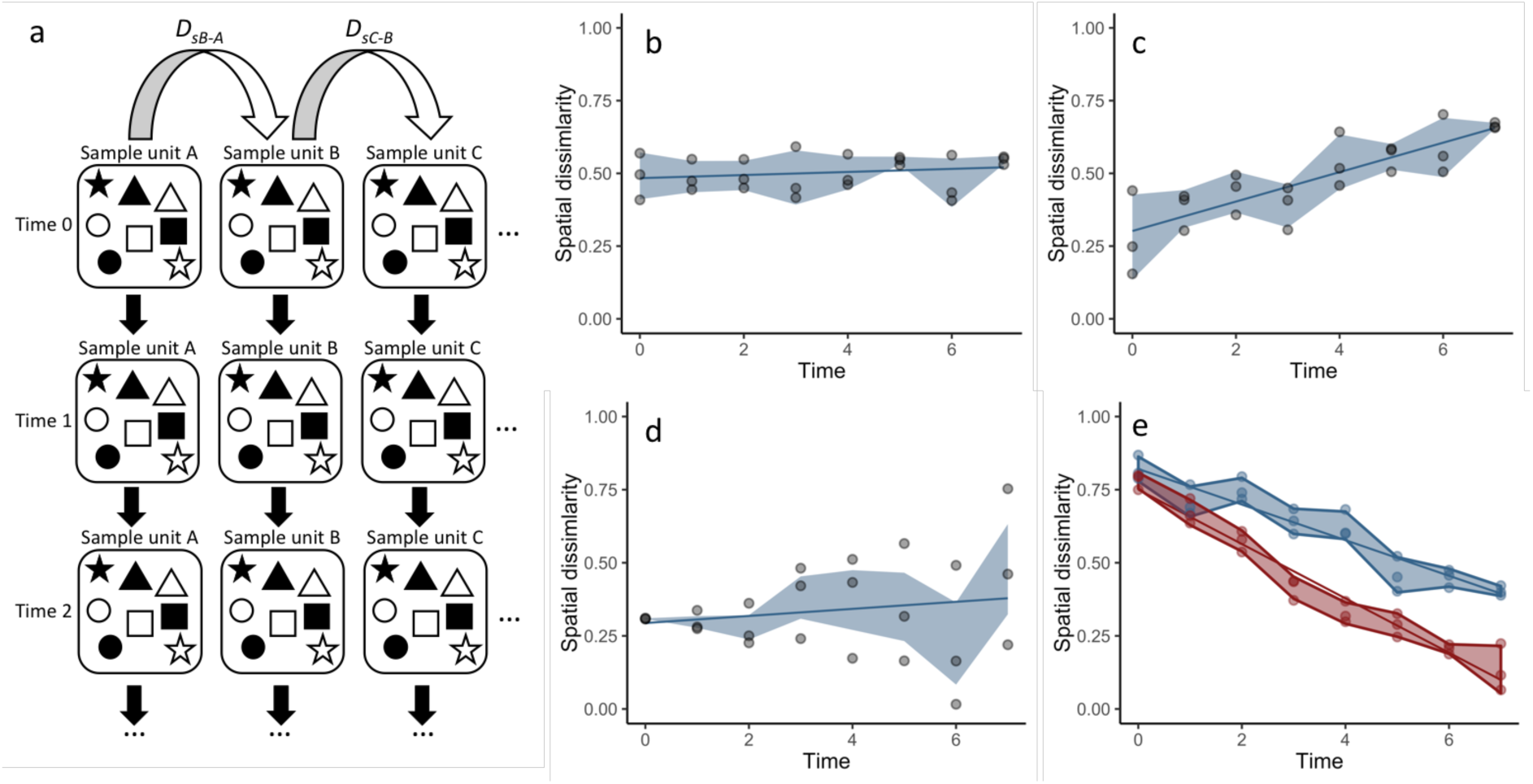
Conceptual figure illustrating the different patterns in (a) temporal community dynamics for a set of sample units (Sample units *A-C*) expected under different hypotheses of community change, visualised as how pairwise dissimilarities (*D*), e.g., Bray-Curtis, would change over eight sampling occasions (Time 0-8): (b) the null hypothesis of no change (‘persistence) or ‘directional’, whereby all sampling units are moving at the same direction and speed in compositional space, (c) ‘divergence’, (d) ‘instability’, and (e) ‘convergence’ (blue line)/ ‘homogenisation’ (red line). Points represent individual pairwise comparisons for three sample units at eight sampling occasions, and the error envelope is the standard deviation of the three values.

### Simple predictions for a single sample unit: Persistence and directional change

Aside from the ‘null’ prediction of no change in composition (‘persistence’; Fig. 4b), one of the simplest predictions of compositional dynamics is that of ‘directional’ change, whereby certain taxa either increase or decrease so that the dissimilarity of a sampling unit to itself at previous sampling occasions increases (or decreases) over time (Fig. 4c); when composition turns over as taxa are replaced, the magnitude of change in dissimilarity can be high. Such a hypothesis could arise from many ecological processes, e.g., has the community undergone ‘succession’, whereby there is a predictable replacement of individual or groups of taxa over time; has it undergone ‘degradation’, whereby rare taxa are lost from the system over time; has it had an ‘invasion’ event, where the relative dominance of one taxon or functional group increases; is it changing in composition in response to a changing biotic or abiotic variable, such as predation or temperature?

### Simple patterns for multiple sample units: Directional change, divergence and convergence

Where a set of sample units are all undergoing the same directional changes in composition over time, pairwise ‘spatial’ dissimilarity among that set of sampling units, would not change significantly over time, because all sampling units are moving in the same direction at the same rate in compositional space because the same taxa are changing in the same ways (Fig. 5b). We also might ask whether sample units are becoming more dissimilar to each other, due to interactions between the changing factors described above and characteristics of the community, such as initial composition or differences in habitat conditions. In this case ‘divergence’ represents the hypothesis where a set of sampling units are all becoming more dissimilar in their composition, so that their pairwise dissimilarities increase over time; this term therefore does not apply to change within one sampling unit (Fig. 5c).

Two very similar patterns predict decreasing pairwise dissimilarity values over time. In contrast to ‘divergence’, under ‘convergence’ taxonomic composition of two or more sampling units becomes more similar over time (Fig. 5e) in a predictable way because of increases or decreases in taxon presence or abundance; this is also called ‘homogenisation’ (Fig. 5e), where there is a concurrent decrease in diversity and dissimilarity becomes relatively low. In contrast, convergence simply implies that between sample-unit dissimilarity declines over time, but not necessarily to a very low level. In the cases where taxa are lost, the predicted overall decline in dissimilarly is large. These hypotheses of change would be relevant if we are interested in the effects of an experimental treatment designed to increase the growth of a particular set of taxa, or under invasion, such as the biotic homogenisation of cities due to urbanisation, or loss of native taxa due to competition with the invading taxa. Note that the predicted effect of invasion on dissimilarity values depends on whether we are considering the trajectory of one sample unit (divergence) or many sample units (convergence) and the among-sample unit variation in other interacting factors.

### More complex temporal patterns: Perturbations, resilience, and cyclical change

More complex patterns in temporal dynamics involve compositional shifts that change in their magnitude or direction over time. Natural or manipulative experiments asking questions about the effects of disturbance on community composition can lead to complex predictions of change. ‘Perturbation’ is a pattern for which the change in composition is initially high but lessens over time and can be detected by comparison to undisturbed treatments or areas (Fig. 6a). The perturbation can occur as either a ‘press perturbation’ (Fig. 6b), such as the removal of a predator or a continuous nutrient addition treatment, or a ‘pulse perturbation’ (Fig. 6c), such as a disturbance event or a population density reduction; these two perturbations result in different rates of change in pairwise temporal dissimilarities. Post-perturbation, a community either could remain in a state that is dissimilar to un-perturbed sample units (‘tipping point’), or could return to its previous state over time (‘resilience’). The prediction of a ‘pivot point’ or ‘threshold’ can be used to test for a significant compositional shift for one or more sampling units over a relatively short period of time, meaning that temporal pairwise dissimilarity among times for each sampling unit is low before, and also after, the pivot point, but is high for temporal pairs across the pivot point (Fig. 6c).

**Fig. 6.**
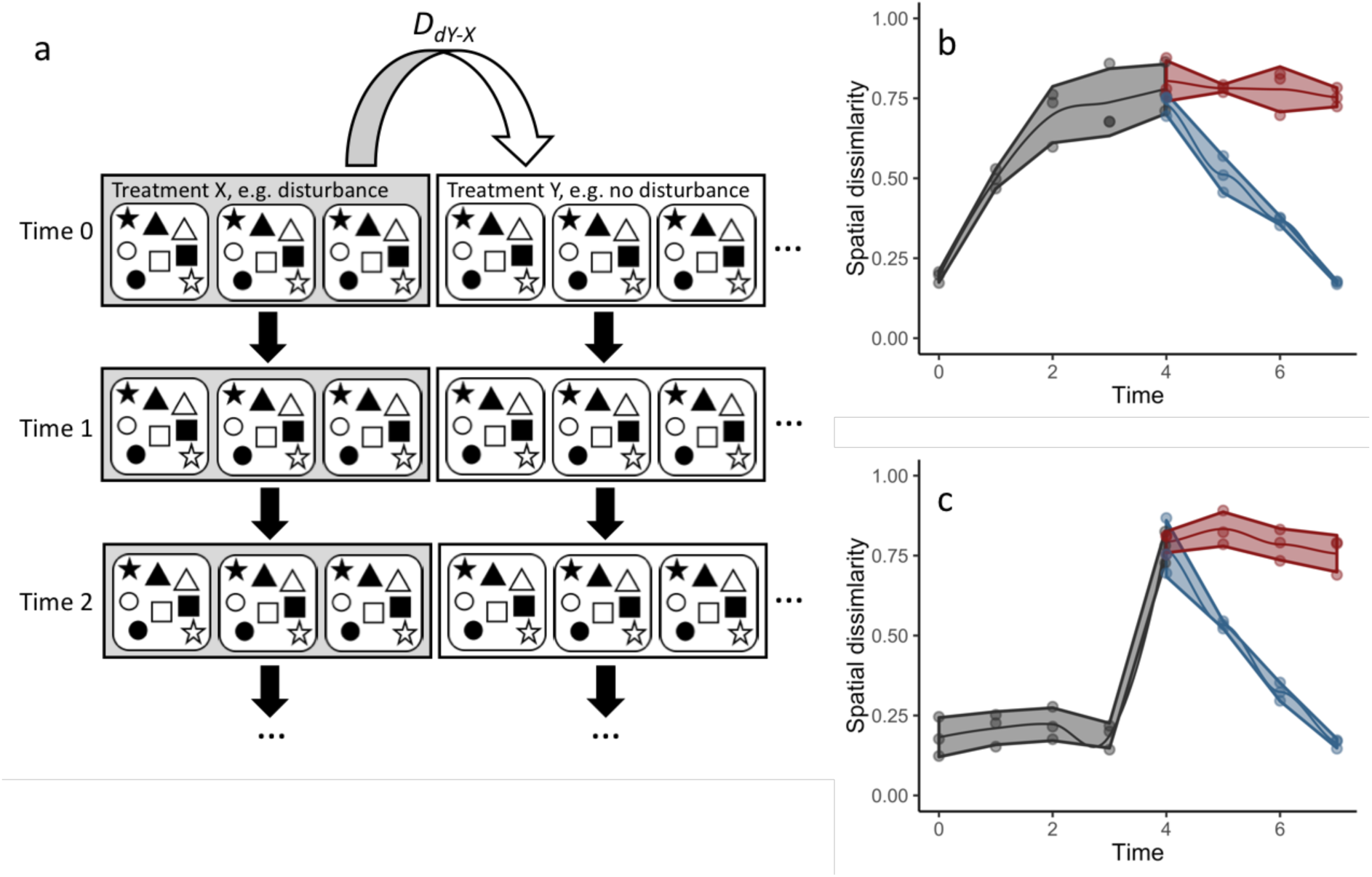
Conceptual figure illustrating the different patterns in (a) temporal community dynamics for a set of experimental sample units in two treatments (*X* and *Y*) expected under different hypotheses of community change, visualised as how pairwise dissimilarities calculated between treatment pairs (*D*), e.g., Bray-Curtis, would change over eight sampling occasions (Time 0-8): (b) a ‘press perturbation’ that occurs at Time 0 or a (c) a ‘pulse perturbation’ that creates a ‘pivot point’ or ‘threshold’ that occurs at Time 3. Both pulse and press perturbations can generate either a ‘tipping point’ (red line) or ‘resilience’ (blue line) type response. Points represent individual pairwise comparisons for three sample units at eight sampling occasions, and the error envelope is the standard deviation of the three values.

Questions about the scale of temporal dynamics in a community, such as the presence of seasonal cycles, lead to predictions of ‘cyclical’ change or ‘periodicity’, which represent shifts in composition that repeatedly diverge and re-converge back to a previous state; dissimilarity increases at short temporal scales but decreases at larger temporal scales, and will alternate between high and low at increasing temporal scales that correspond to some cyclical process, e.g., warming events (Fig. 4d).

### Predictability of change: Instability and stabilisation

Questions about the predictability of compositional changes over time resulting from interactions between causal-change processes, such as temperature shifts throughout a year, and biotic or abiotic differences among sampling units, could also lead to hypotheses regarding the variability in relative compositional change. For instance, ‘instability’ predicts significantly increasing temporal variation in dissimilarity for a set of sampling units because relative increases and decreases among taxa are not correlated or consistent; change is not directional although mean dissimilarity would also likely increase with the variance (Fig. 5d). In contrast, ‘stabilisation’ predicts the opposite: significantly decreasing temporal variation in dissimilarity as initially significant compositional change lessens over time; pairwise dissimilarity is initially high at short temporal scales (the opposite of Fig. 5d).

### Taxon dynamics: Synchronicity

In many cases, we are interested in asking if different components of the community are responding in similar or different ways over time. Where individual taxa, or particular groups of taxa (e.g., functional groups), change in similar ways at the same time (‘synchronicity’), we can detect this because among-taxon-group dissimilarity increases, but within-taxon-group dissimilarity does not change (this involves comparing the pairwise sampling unit dissimilarities separately for different taxonomic subsets) (Fig. 7a). Synchronicity may result in directional or other types of compositional dynamics patterns within one sampling unit or a set of sampling units (Fig. 7b).

**Fig. 7.**
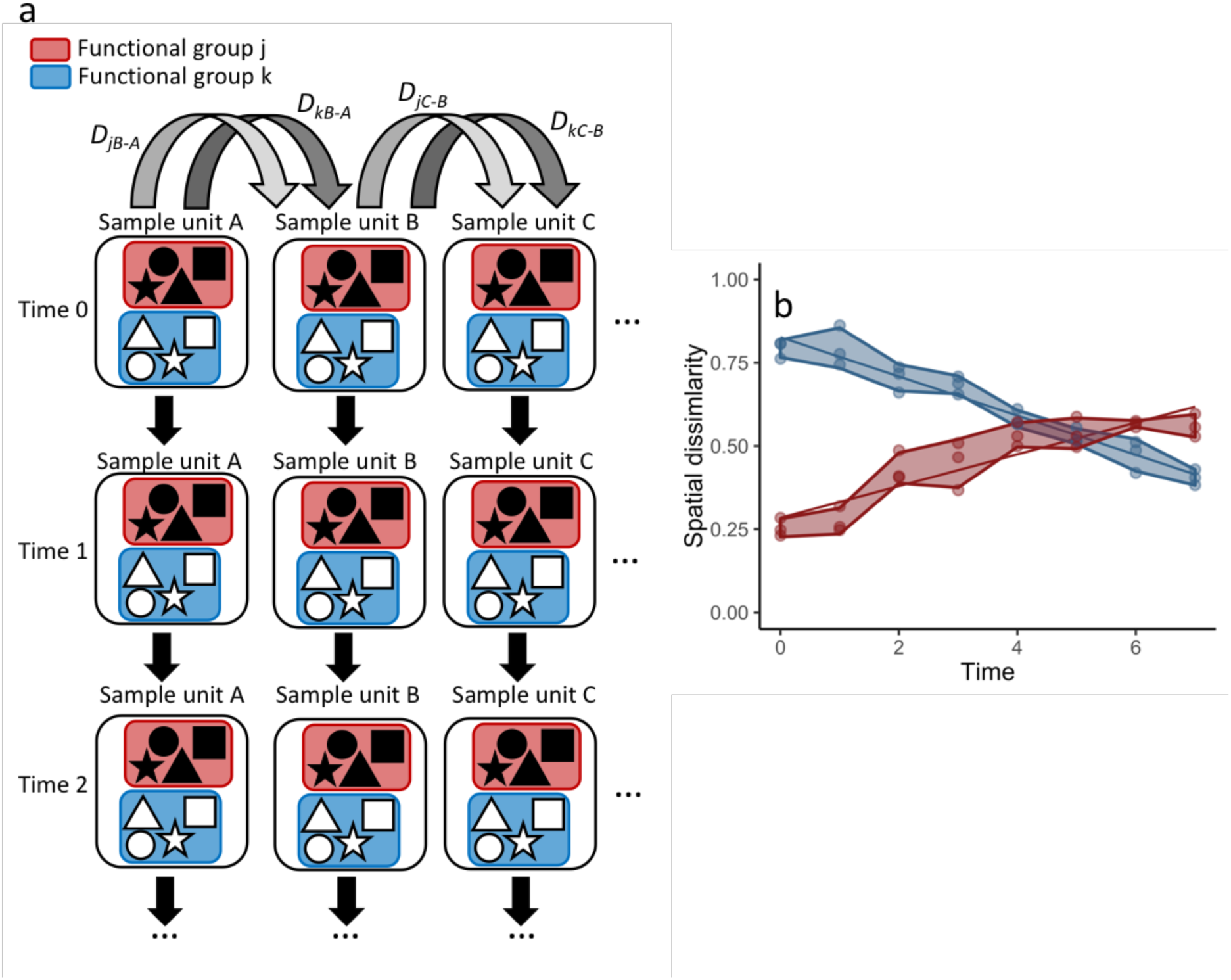
Conceptual figure illustrating the idea of ‘synchronicity’, whereby different functional or taxonomic subsets of a community are changing differently from other subsets (but the taxa within those respective groups are changing synchronously). In this case, (a) pairwise dissimilarities are calculated separately for the different subsets and (b)any possible differences in changes over time between the two subsets can be observed, e.g., one functional group might be becoming more homogeneous (blue), whereas another might be changing directionally (red). Points represent individual pairwise comparisons for two different functional groups (red and blue) from three sample units at eight sampling occasions, and the error envelope is the standard deviation of the three values.

Hypotheses of biological processes causing dynamics may be very specific, such as a decrease in certain prey taxa in response to a predator, or more general, such as a shift towards a grassland community in response to fire disturbance of a woodland or an increase in annual plant taxa in spring compared to winter. More specific and directional (increases or decreases) hypotheses lead to more powerful tests of compositional change; we strongly encourage researchers to generate such hypotheses *a priori* and to pair these with predictions that are as quantitative as possible, e.g., define the predicted timescales of variation.

## C. WORKED CASE STUDY EXAMPLES

Here, we provide three worked examples to demonstrate the hypothesis-testing approach we advocate for different types of community dynamics. We used open-access datasets to test for: (1) significant divergence in temporal community dynamics among disturbance treatments imposed upon plant communities in experimental grasslands, (2) neutral community dynamics for a fully-censused forest plot, and (3) seasonal (cyclical) temporal dynamics in marine aquatic bacterial communities. We present a brief description of each dataset and describe the results and interpretation for the relevant exploratory data analysis (EDA; see section ‘D. Road map for analysing temporal community dynamics’) and hypothesis-testing analyses. More detailed descriptions, explanations and R code for each case study are provided as supporting information (Data S2-4).

### Testing for divergent community dynamics within a controlled experiment

Porensky et al. (2016a, 2016b) measured plant species composition annually from 2007 to 2014 in quadrats on 24 transects to which six grazing treatments were applied since 1982 (four transects in each treatment): none (exclosure), light, moderate, heavy, heavy-to-light (lightly grazed since 2007; ‘new light’) and heavy-to-none (new exclosure since 2007). They calculated the mean cover in 25 quadrats for each species on transects. We tested the hypothesis that increasing grazing intensity causes increasing divergence (directional change; as detailed in section B) in species composition relative to the control (no-grazing) treatment.

EDA (as described in section D)) showed that the 24 transects were dominated across all measurements by ten common species out of a total of 92 species (Data S2: row and column summaries, relative abundance plots and rank abundance curves). Species turnover (section D 4.0) through time was high; mostly 30% or greater for time period-treatment combinations (Fig. 8). Principal coordinates analysis (PCoA; section D 1.4 Ordination) suggested that the main compositional differences corresponded to the grazing treatments, and that transect composition changed over time, but not markedly (Data S2: PCoA analysis and figure). PERMANOVA (section D 1.4a) showed that transects were significantly compositionally distinct and changed significantly over time in different ways (Data S2: PERMANOVA outputs – all three coefficients for a two-way PERMANOVA with transect, year and their interaction were significant *P* < 0.05). Partitioning spatial and temporal beta diversity (section D 1.2) showed that two of the control (exclosure) treatments differed significantly (Fig. 9). Principal Response curves (section D 1.4b) showed that the grazed treatments differed from the control and that the relative abundances of four species in particular were influential in driving these differences among treatments, but did not show increasing divergence over time (Fig. 10). This result is consistent with the analysis of temporal coherence (icc = −0.04; see section D 1.4a and Data S2 for detailed description of this statistic), which showed that transects were not synchronous in their temporal changes (Data S2: temporal coherence analysis). Thus, we conclude that treatment differences in turnover and shifts in composition were not consistent or directional and so we reject our hypothesis that increasingly grazed transects diverged more over time relative to the control.

**Fig. 8:**
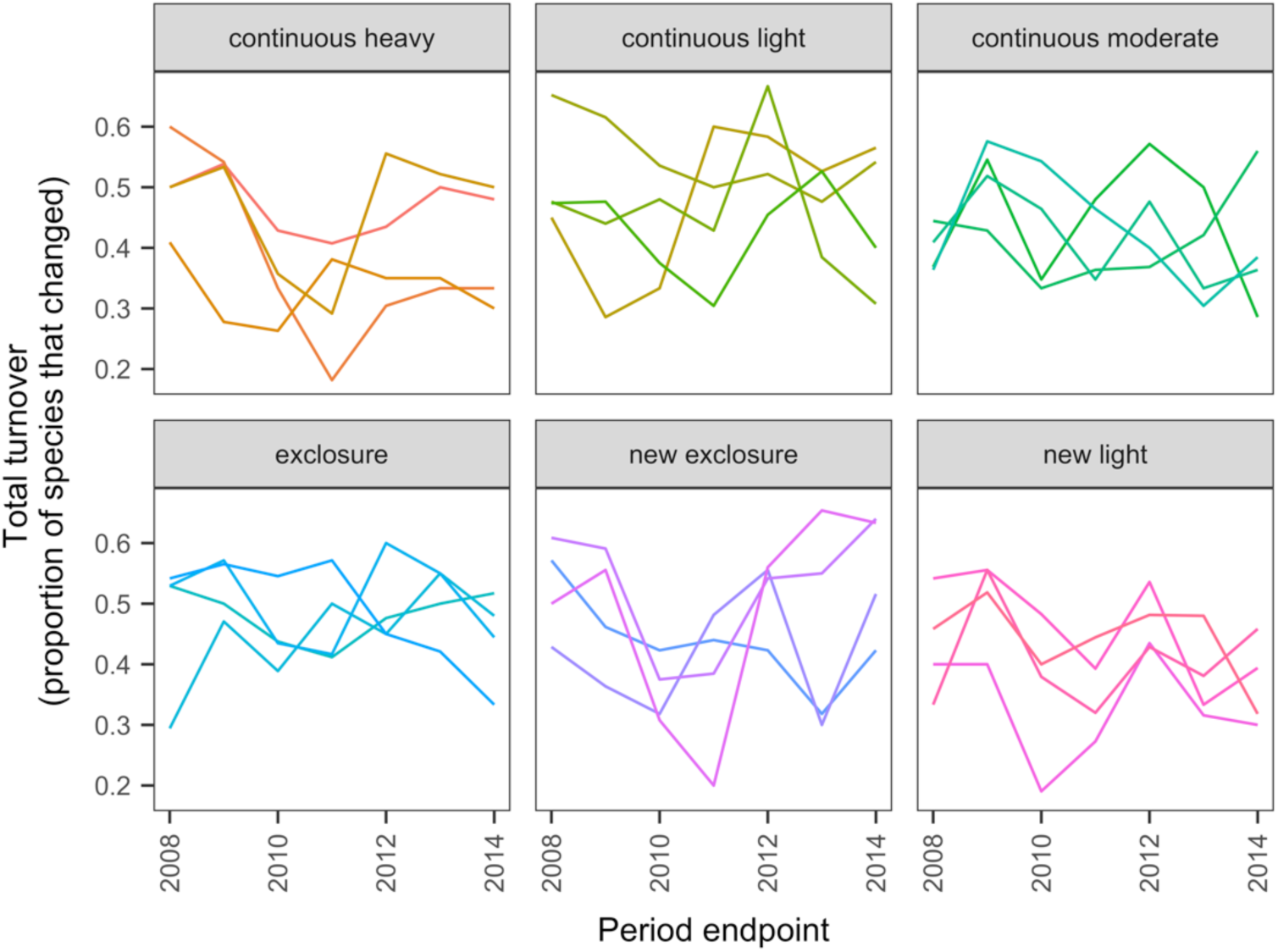
Changes in compositional turnover over time for data from the Porensky et al. (2016a, b) case study. Four transects were subjected to one of six grazing treatments and were sampled annually from 2007 to 2014. Note that “exclosure” treatment is the control (no grazing). The different lines represent different transects.

**Fig. 9:**
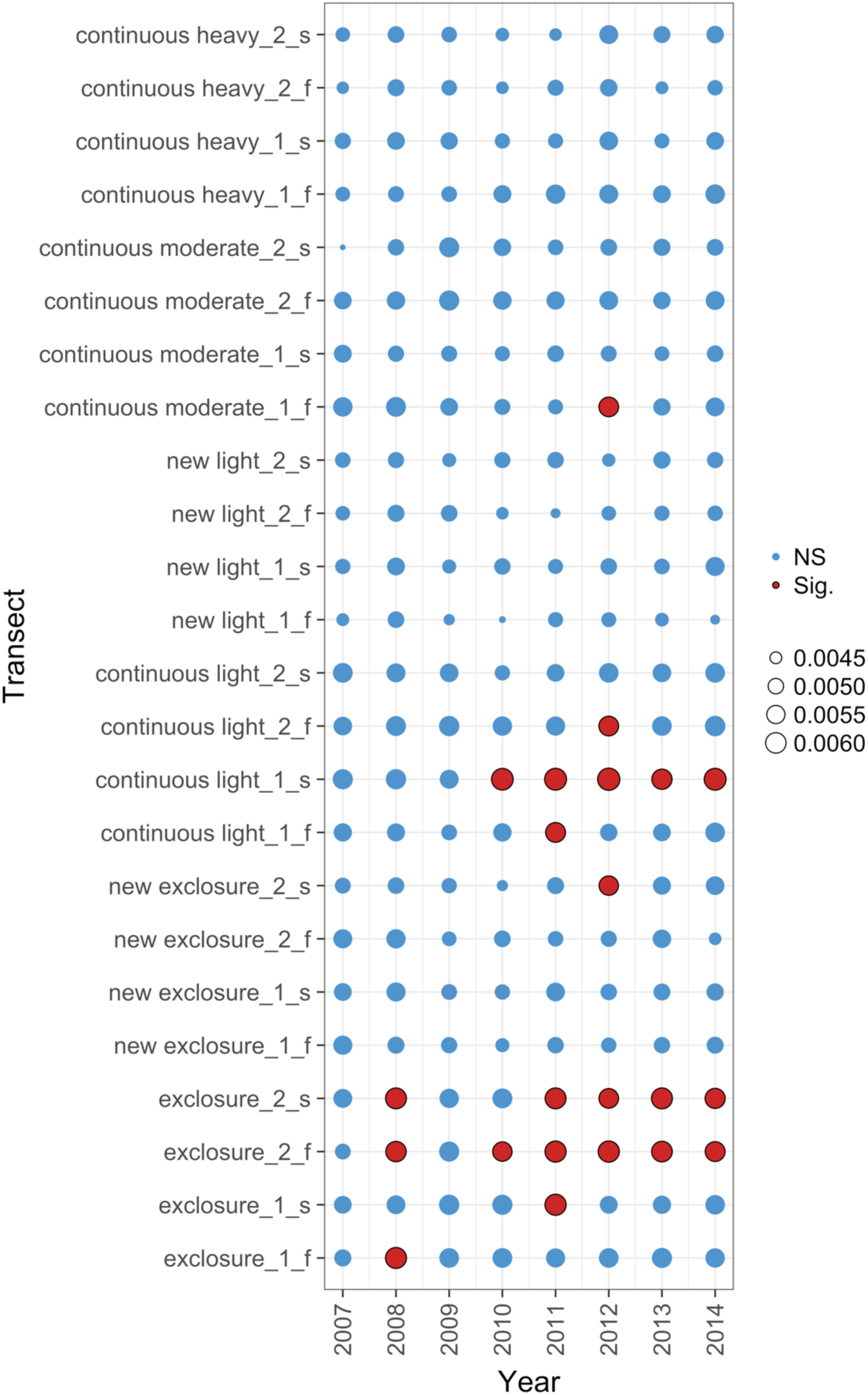
Local contributions to beta diversity (LCBD) for each transect at each sampling time for data from the Porensky et al. (2016a, b) case study. Plot shows the relative contributions of the different transects and times to the overall beta diversity. Symbol size is proportional to the LCBD value and significance is indicated by the red, outlined circles; blue circles are not significant. Transects had a grazing treatment and were established in one of two replicate pastures (1 or 2) and were either north-facing (f) or south-facing (s). Grazing treatments were: none (exclosure), light, moderate, heavy, heavy-to-light (lightly grazed since 2007; ‘new light’) and heavy-to-none (new exclosure since 2007).

**Fig. 10:**
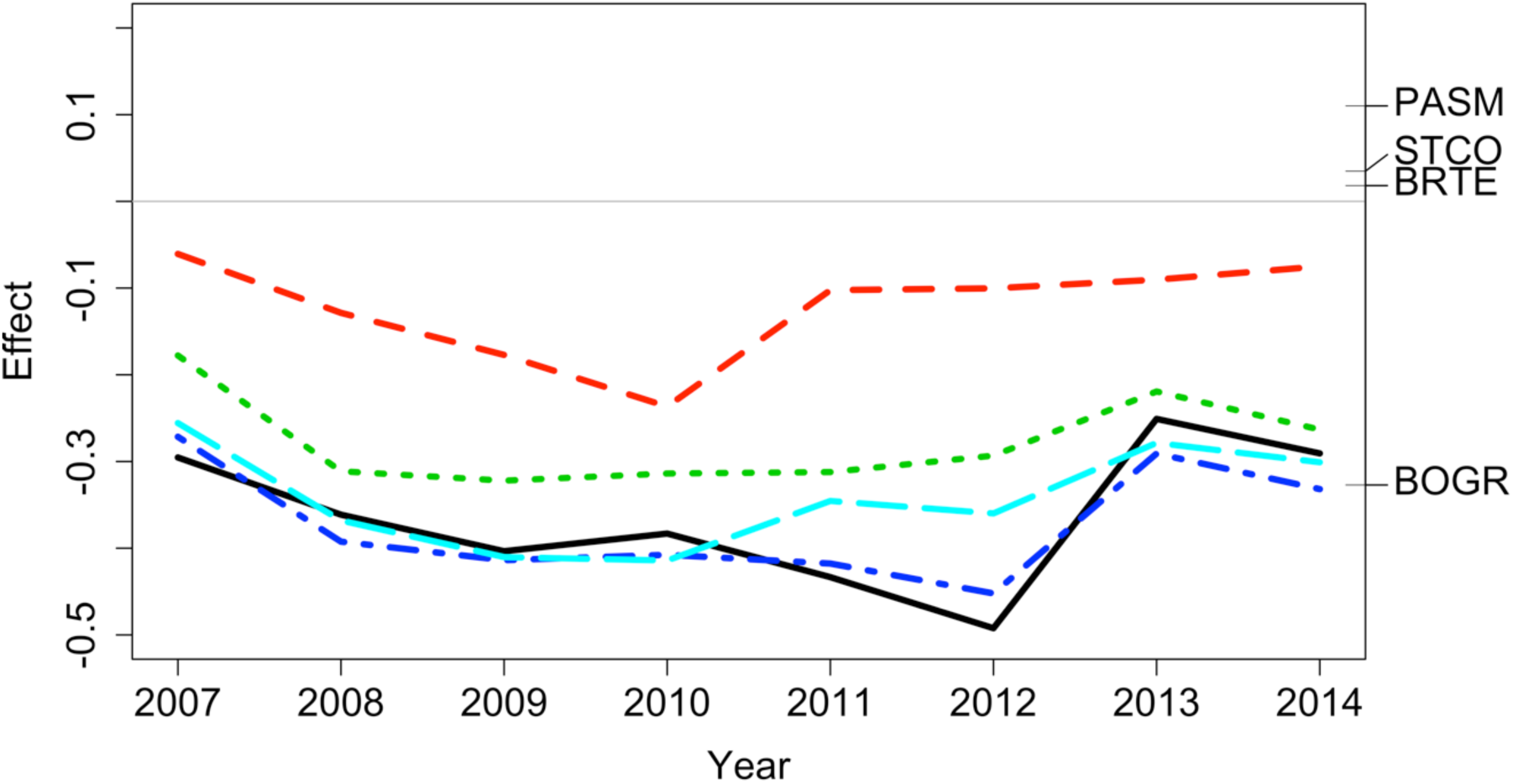
Principal response curve analysis of data from the Porensky et al. (2016a, b) case study. Results show that treatments differed in plant species composition from the control (exclosure; grey line) but the differences between treatments were inconsistent over time. Treatments were: new exclosure (since 2007; dark blue), continuous light (red), new light (since 2007; light blue), continuous moderate (green), and continuous heavy (black). Four particularly influential species where either lower (BOGR = *Bouteloua gracilis*) or higher (PASM = *Pascopyrum smithii*, STCO = *Hesperostipa comata*, and BRTE = *Bromus tectorum*) in abundance than in the control (black horizontal lines represent the abundance of each species relative to the control).

### Testing for neutral community dynamics across spatial scales

The 50-ha forest census plot on Barro Colorado Island, Panama, has been censused eight times between 1982 and 2015 (Condit 1998, Hubbell et al. 1999, http://ctfs.si.edu/webatlas/datasets/bci/, 2005). This plot was established in part to test for neutral community dynamics (Hubbell 1979) by fully censusing all woody stems greater than 1 cm diameter at breast height every five years. Under this hypothesis, we expected to see no directional or other patterns in temporal community dynamics (‘persistence’; as described in section B.). EDA showed that the dataset consisted of 324 species generating a highly right-skewed abundance distribution (most species were rare) (Data S3: EDA analyses and section D). A line graph plot of species’ abundance values over time showed very little change in total abundance over time (Data S3: line graph of abundance; Appendix S2: Univariate linear methods). Two analyses that allowed us to test for stability in community composition in time, i.e., change in species composition is lower than expected if individual trees were randomly distributed in time (keeping their spatial positions fixed), showed that change in species composition was lower than expected: (1) Zeta diversity analysis (Data S3: zeta diversity and section D 2.0) showed that the presence of species across census times was more consistent than expected (Fig. 11) and (2) total turnover in species abundances (Data S3: total turnover rates analyses and section D 4.0) at each of the seven time periods was lower than expected (Fig. 12), particularly when the plot data were at larger spatial scales. Thus, we conclude that the observed lack of pattern is consistent with our hypothesis of neutral community dynamics.

**Fig. 11.**
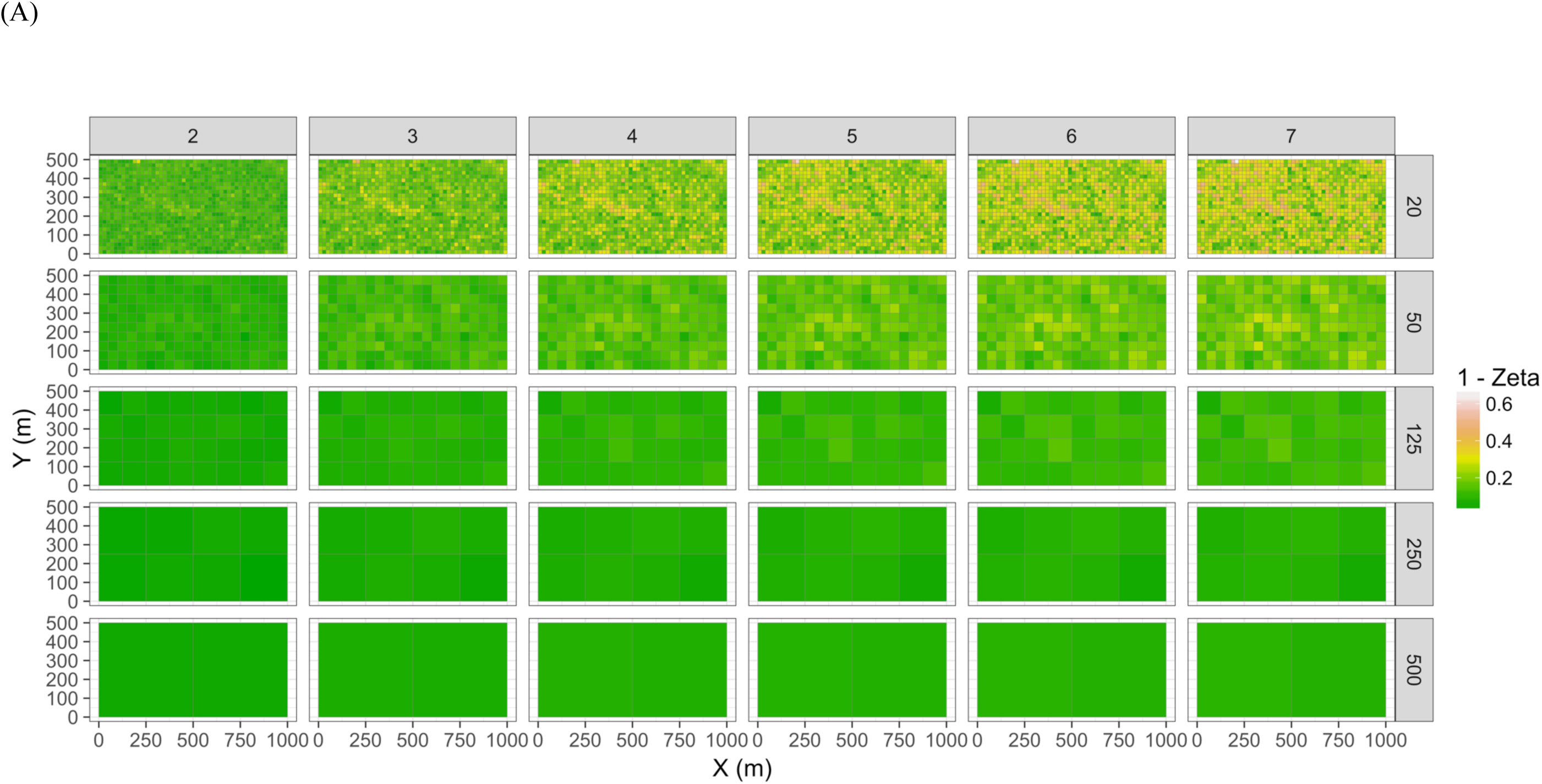
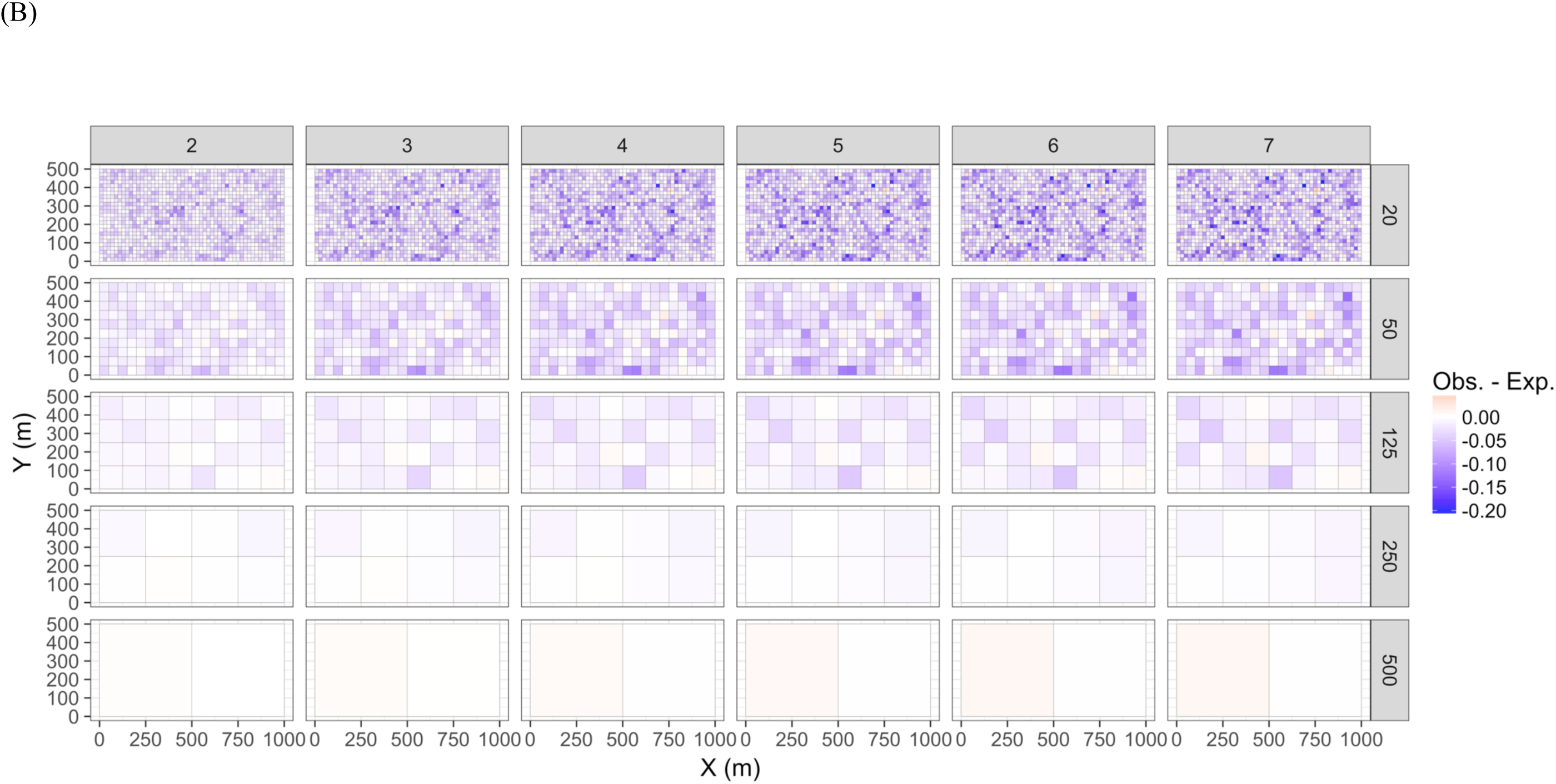
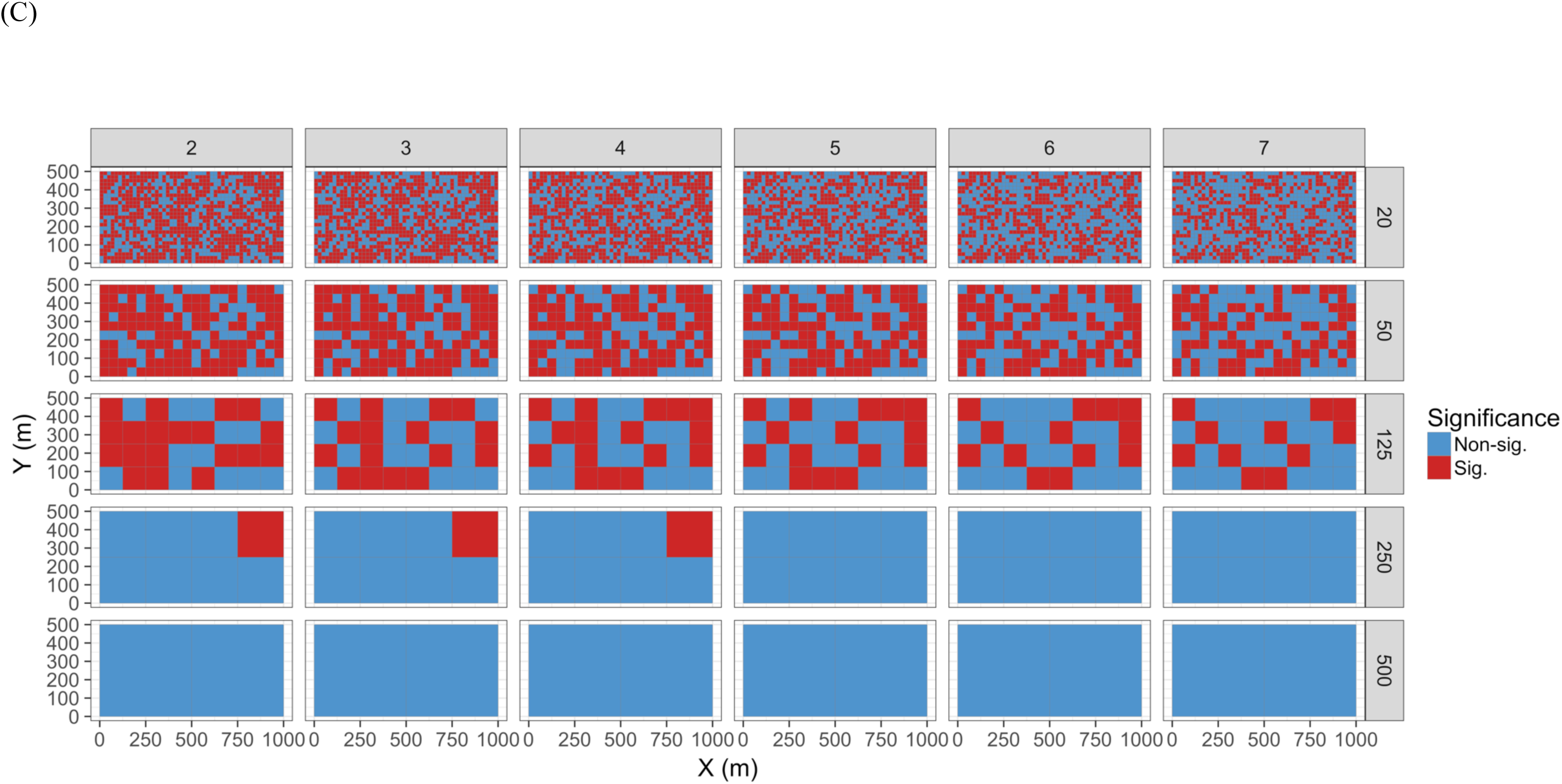
Zeta diversity analysis, based on the number of species shared by a set of temporal samples from the Barro Colorado Island case study. Results show that the woody forest community in the 50-ha forest census plot on BCI has been significantly more stable over time than random expectation, particularly at large spatial scales where richness is higher. Maps show for 1 - zeta diversity calculated for quadrats of increasing sizes (20 × 20 m, 50 × 50 m, 125 × 125 m, 250 × 250 m and 500 × 500 m) and for between 2 and 7 orders of zeta (the number of temporal samples compared): (A) the observed values, (B) observed - expected values under a null model of random change in species composition in quadrats over time, and (C) significance of the observed values. Results are given as 1 - zeta so that they are comparable to the turnover results in that higher values represent higher dissimilarity in species composition.

**Fig. 12.**
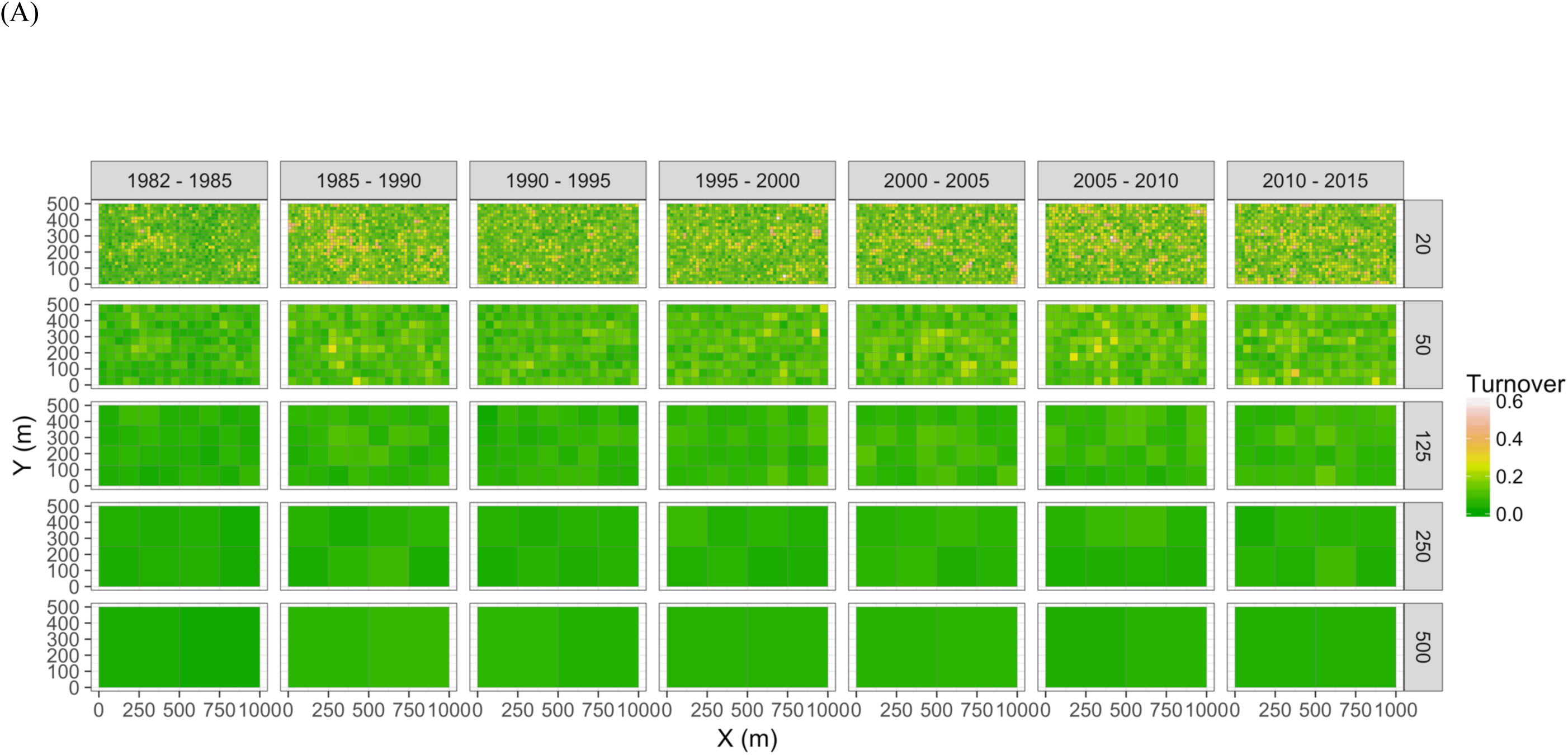
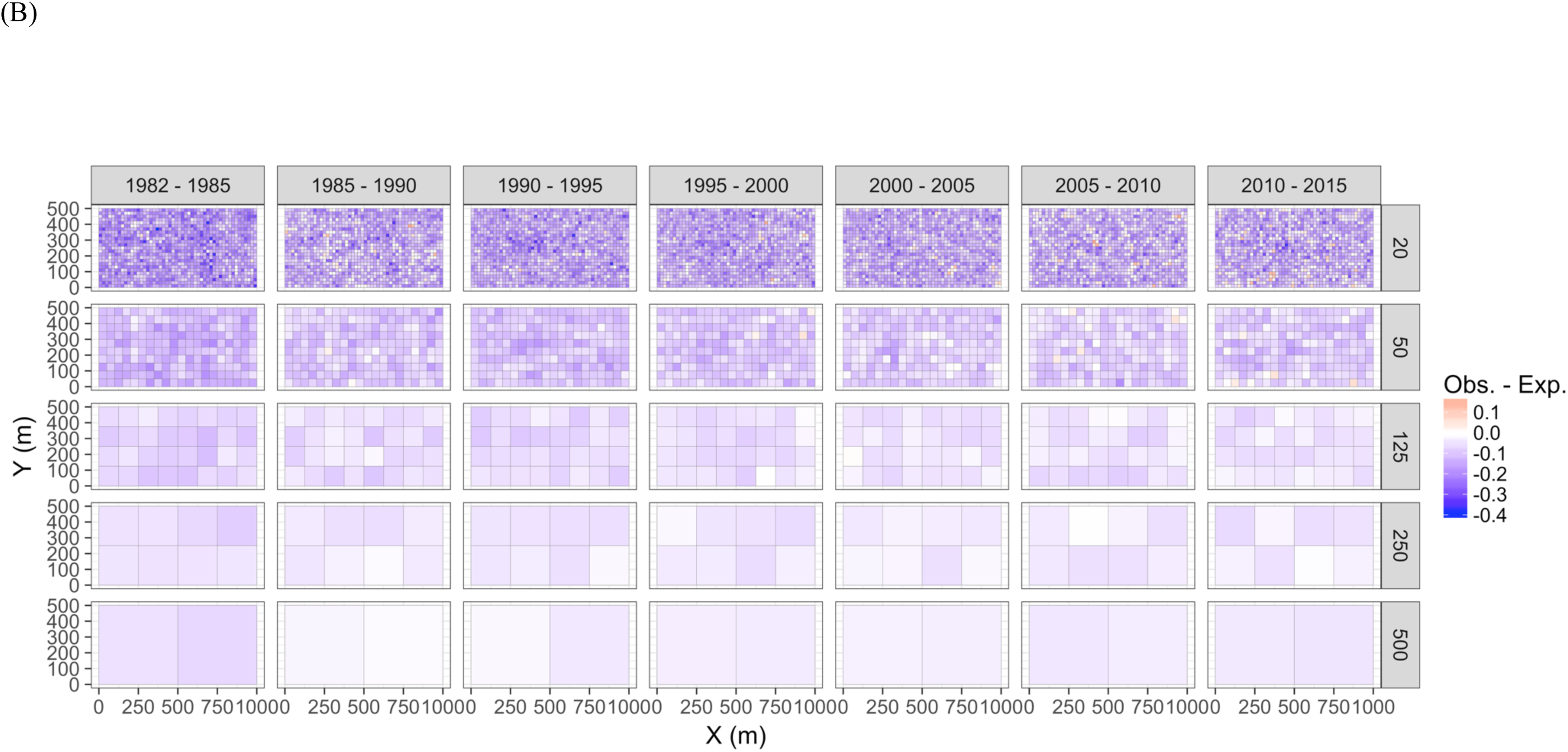
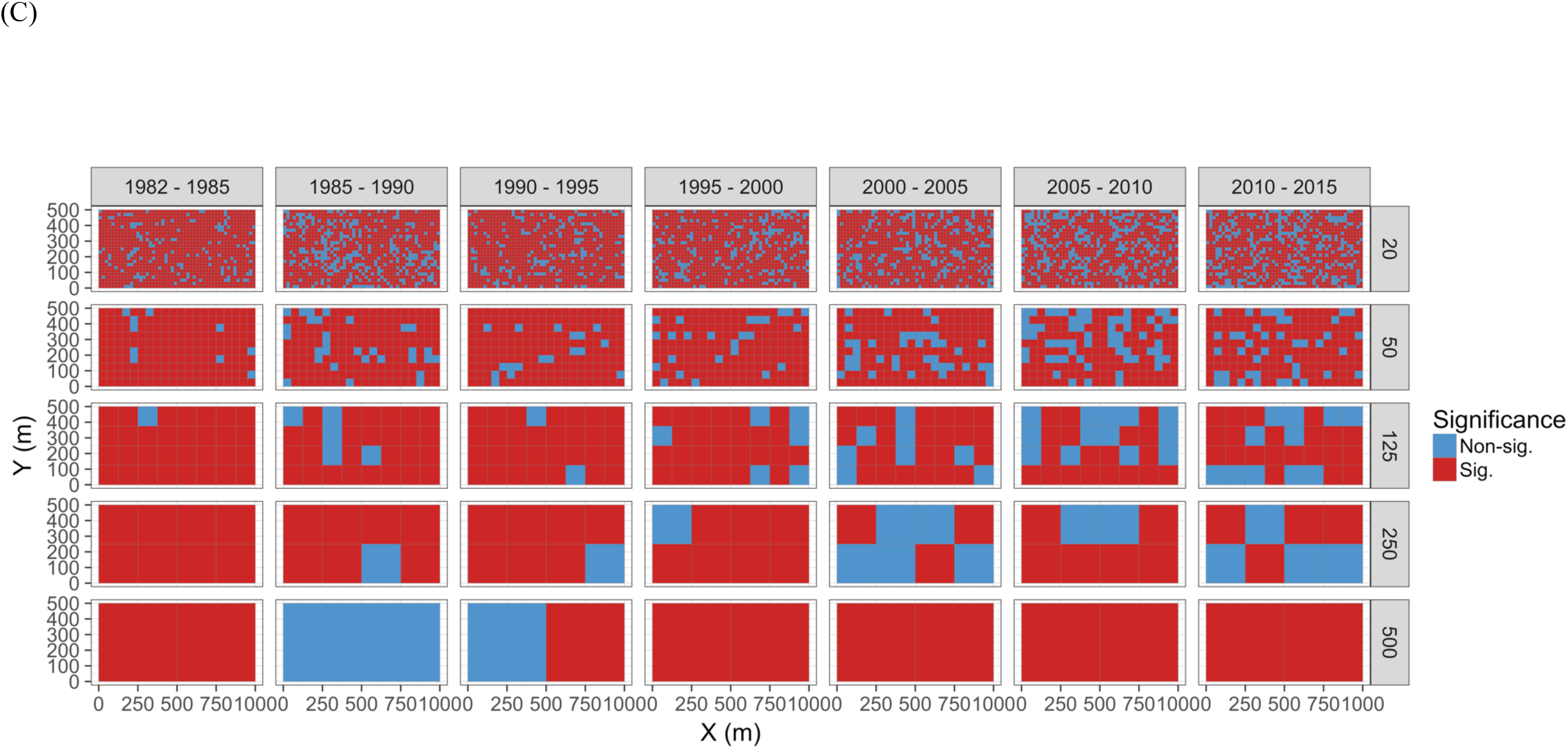
Turnover analysis results for the seven time intervals for the Barro Colorado Island forest community. Results show that the proportion of species turning over each measurement interval was significantly lower than under random expectation, particularly at large spatial scales where richness is higher and composition is more stable: Maps show proportion of species turnover calculated for quadrats of increasing sizes (20 × 20 m, 50 × 50 m, 125 × 125 m, 250 × 250 m and 500 × 500 m) across the seven time intervals: (A) the observed values, (B) observed - expected values under a null model of random change in species composition in quadrats over time, and (C) significance of the observed values.

### Testing for seasonal community dynamics

Gilbert et al. (2012) measured bacterial taxonomic composition monthly over six years using pyrosequencing of the 16S rRNA gene for water samples from a single site off the coast of Plymouth, United Kingdom (source data can be obtained from the journal’s online supplementary file). We tested the hypothesis that subsets of the community would show cyclical patterns of temporal change due to seasonality. EDA (section D) showed that the dataset consisted of 74 operational taxonomic units (OTUs) across 62 temporal samples. Most taxa were common through time. Proportional turnover between temporal samples varied from 4% to 48% (Data S4: turnover calculation and section D 4.0), which is relatively low, and was consistent with the detrended correspondence analysis (DCA) result showing a short gradient length of 1.8 for axis 1 (Data S4: DCA and section D. EDA). Similarly, PCoA graphs (section D. EDA) showed little consistent change within years (Fig. 13). However, 13 of the OTUs were deemed to be highly variable in that they changed by more than 500 reads across the sampling period (mean number of reads = 296 ± 829 S.D..; Data S4: row and column summaries, relative abundance distributions and abundance plots and section D. EDA).

**Fig. 13.**
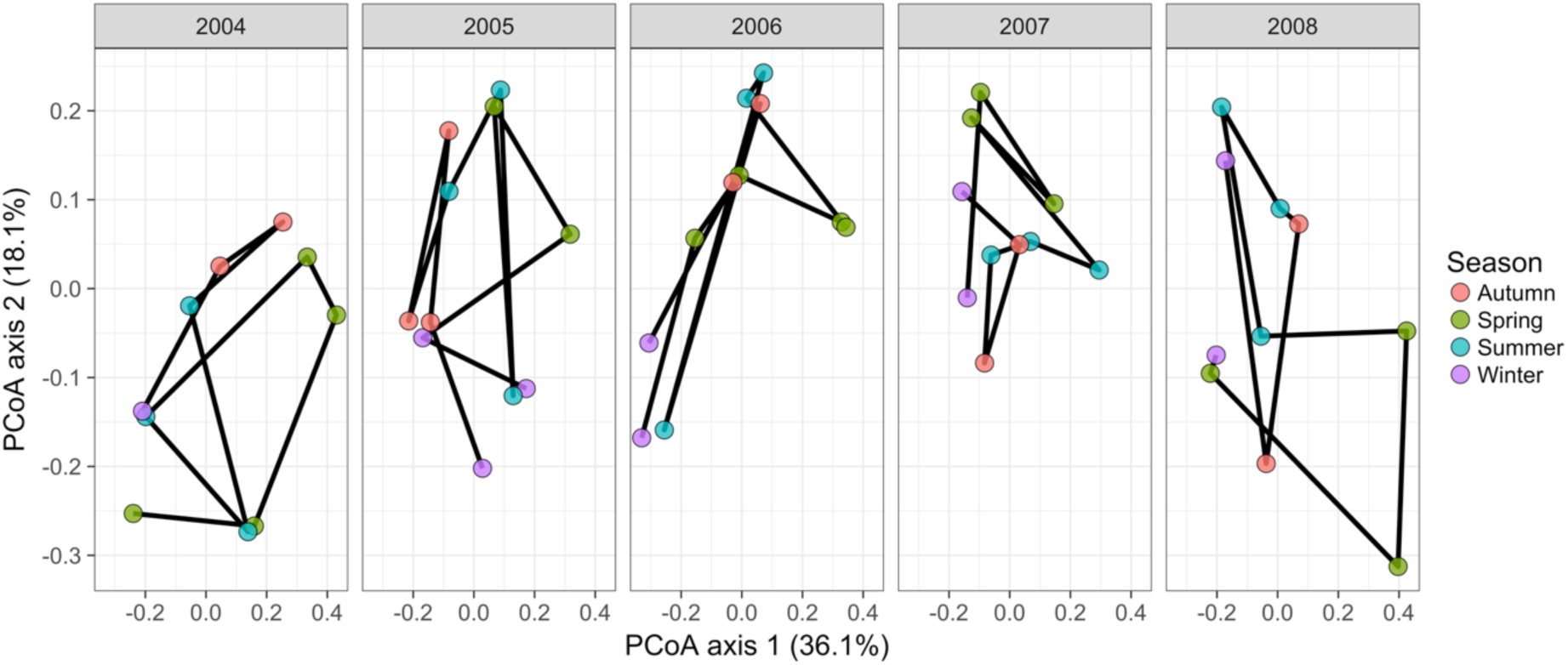
Principal coordinates analysis of 74 marine bacterial OTUs across 62 temporal samples from the Gilbert et al. (2012) case study. Results show no visible consistency in compositional change by season (coloured points) within or between years for the single site analysed. Black lines link samples taken consecutively within years.

Time-lag analysis (section D 1.3) showed that although there was a significant increase in dissimilarity within increasing time lag, variability was high, showing that temporal changes did not occur in a simple divergence over time pattern (Fig. 14). Temporal synchrony analysis (sensu Loreau and de Mazancourt 2008) showed that some subsets of OTUs in the community were significantly changing in similar ways (Synchrony = 0.27; *P* < 0.05; Data S4: temporal synchrony analysis and section D 3.0). When patterns in these variable taxa were examined through time using zeta diversity analysis (Hui and McGeoch 2014) (Data S4: zeta diversity analysis; section D 2.0), which is based on the presence of species in samples, winter samples shared significantly more species across sets of temporal samples than autumn, spring and summer (Fig. 15; ANOVA, *P* < 0.01; Appendix S2: Univariate linear methods). Further analysis of the most variable taxa using the raw dissimilarity values based on abundance data calculated for temporal samples within seasons showed clear cyclical patterns (Fig. 16; Data S4; section D 1.1). Thus, we conclude that the patterns are consistent with our hypothesis of seasonal dynamics in community composition, largely driven by a small subset of OTUs that varied in their abundance, rather than presence, over time.

**Fig. 14.**
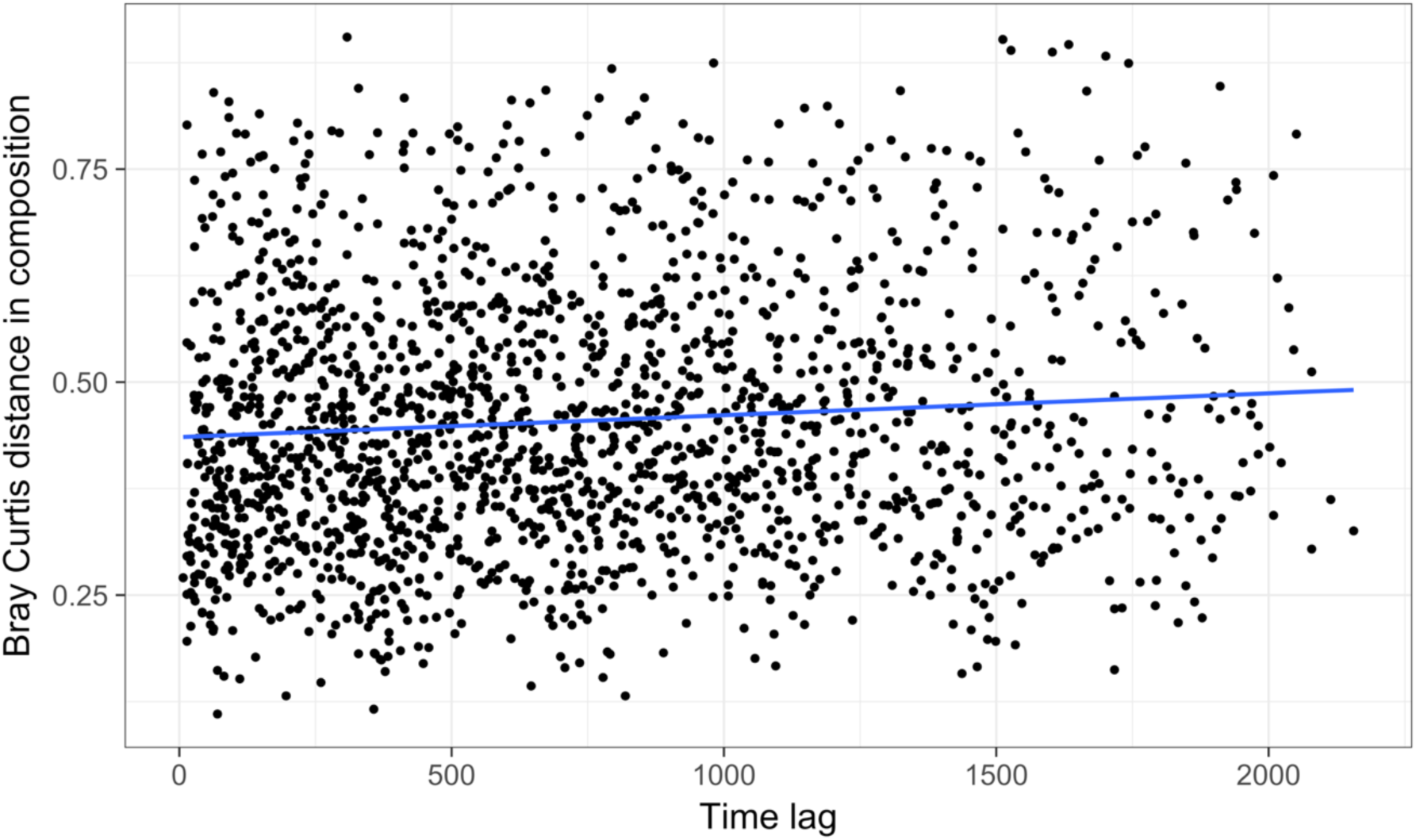
Time lag analysis of 74 marine bacterial OTUs across 62 temporal samples at one site off the coast of Plymouth, United Kingdom from the Gilbert et al. (2012) case study. Results show high variability in pairwise Bray-Curtis dissimilarity values with slight divergence occurring over time. The blue line is a linear regression line. The high variability suggests that there is not consistent, directional temporal change at this site.

**Fig. 15.**
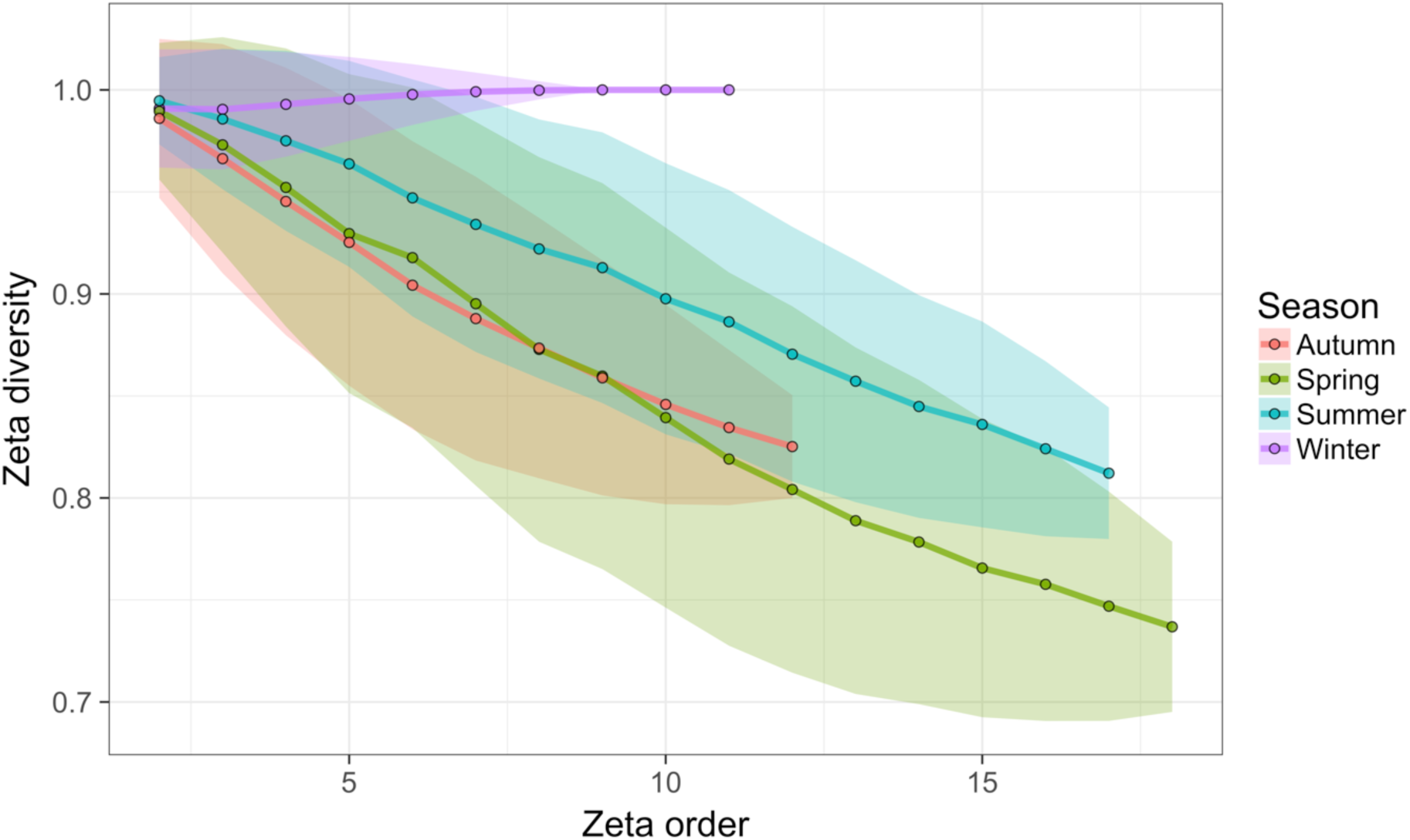
Zeta diversity analysis of the 13 marine bacterial OTUs that were most variable in abundance across 62 temporal samples from the Gilbert et al. (2012) case study. Results show that spring samples were more diverse than winter or summer samples. Zeta order represents the number of temporal samples compared and so each point on the graph shows the mean number of OTUs shared among these different numbers of temporal samples.

**Fig. 16.**
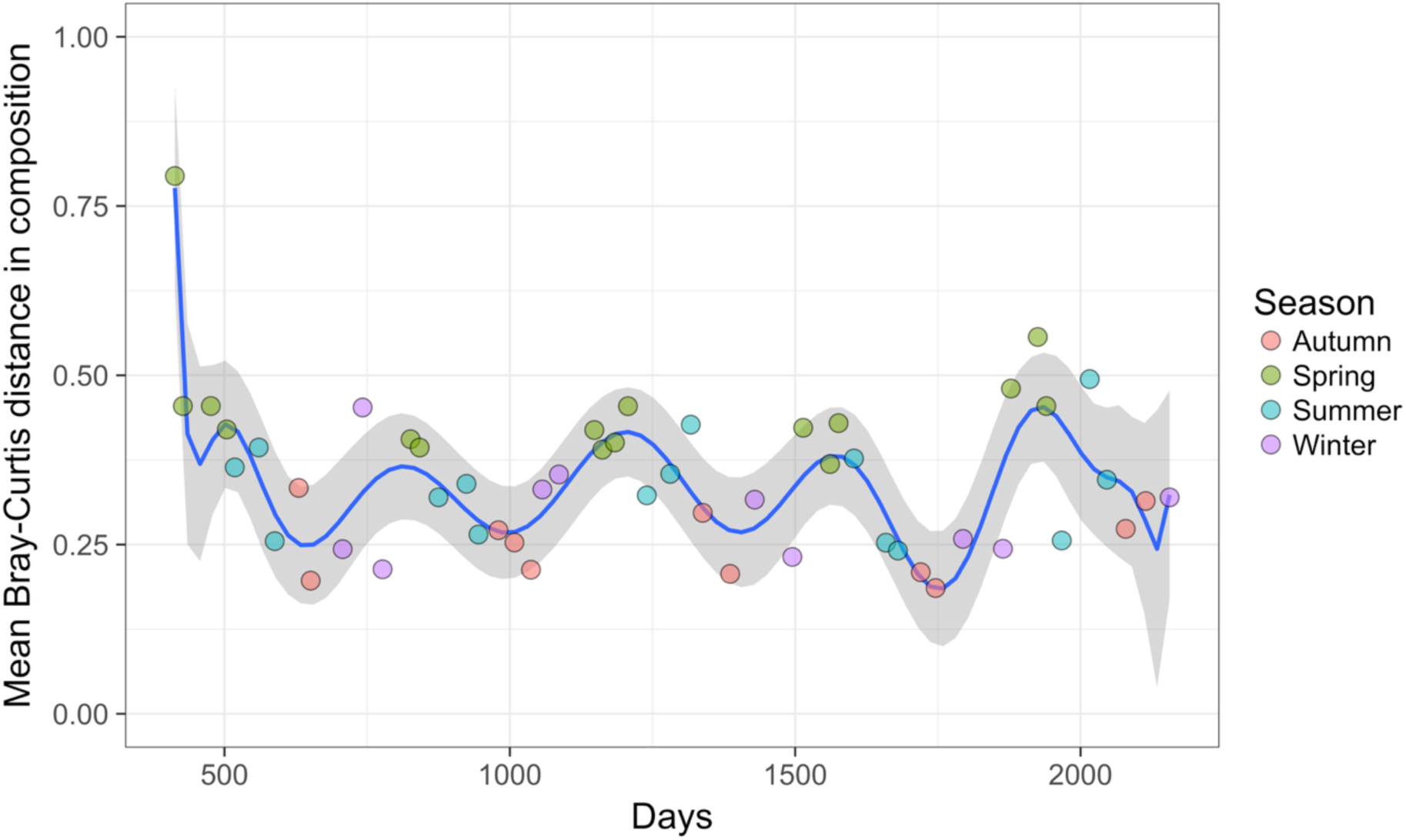
Change in the mean Bray-Curtis dissimilarity values calculated for temporal samples (n = 62) grouped by season from the Gilbert et al. (2012) case study. Results are for the 13 most variable marine bacterial OTUs at the one site analysed, showing cyclical community dynamics. The blue line is a polynomial regression line with 16 terms. Multiple comparisons from an analysis of variance showed that samples taken in spring were significantly less similar to other spring samples than were samples taken in summer, autumn, or winter (*P* < 0.001). Samples taken in summer were significantly less similar to other summer samples than were samples taken in autumn (*P* < 0.05).

## D. ROAD MAP FOR ANALYSING TEMPORAL COMMUNITY DYNAMICS

The two most common sets of methods seen in our review of the temporal community dynamics literature were descriptive (mostly tables and figures that show changes in taxon presence or abundance) and ordination-based methods. Although these techniques are useful for illustrating temporal trends and qualitatively exploring patterns in the data, they are used less often for testing specific quantitative hypotheses of patterns and processes in community dynamics. We recommend that researchers augment such descriptions with methods that quantify temporal dynamics in taxonomic composition in more powerful ways, thus allowing more quantitative and potentially comparable descriptions of spatiotemporal patterns and explicit hypothesis-testing of mechanisms underlying observed patterns in community dynamics.

Once the hypotheses to be tested have been generated, we suggest that researchers undertake five steps: (Step 1) perform exploratory data analysis (EDA), (Step 2) choose multivariate methods that can test the *a priori* temporal hypotheses, (Step 3) test for statistical significance, (Step 4) report on the analysis workflow, and (Step 5) consider how to present the results of the selected methods in the text and in figures.

### Step 1: Exploratory data analysis (EDA)

As with any data analysis, the first step in the analysis of temporal community dynamics is an exploratory data analysis (EDA) phase (Zuur et al. 2010). This generates basic information that can be used to both error-check the data and to gain an understanding of key aspects of the dataset (Table 1). R code for the relevant EDA steps is provided in three worked examples, described previously (Data S2-4).

**Table 1:** Recommended steps for exploratory data analysis (EDA) of community data prior to selecting analysis methods for temporal data. R code for the methods is provided in the supplementary materials Data S2-4.

| Step | Description | Rationale | Methods |
| --- | --- | --- | --- |
| 1 | Count the number of taxa,<br>the number of samples and<br>the number of empty<br>samples | Some ordination methods are influenced<br>by rare taxa; identify empty samples;<br>determine replication for significance<br>testing against explanatory variables | Row and column totals of the sample × taxon matrix.<br>Plot the relative abundance distributions. |
| 2 | Examine taxon frequency<br>distributions and response<br>curves | Some ordination methods assume either<br>unimodal or linear pattern of frequency<br>distribution | Plot the abundance of species across time or other<br>explanatory variables. |
| 3 | Determine the ratio of temporal to spatial variation in composition | If temporal change is to be detected within different spatial sampling locations, the analysis must adequately tease apart the variation in time and space. Where little spatial variation exists, spatial samples could be averaged at each time. | Mean spatial and temporal turnover should be calculated following Collins et al. (2000) to allow a ‘global’ comparison of the amount of compositional variability among samples from each time and among times (see Hallett et al. 2016 for details); it can be calculated for different subsets of samples or times for which different predictions about temporal dynamics have been made, e.g., different experimental treatments.<br>Ordination approaches, such as DCA (ter Braak and Šmilauer 2015), can be used to quantify spatiotemporal variation and to identify ‘outliers’, i.e., samples that are very different to other spatial or temporal replicates. |
| 4 | Determine the optimal dissimilarity measure and whether data should be standardised | The dissimilarity measure selected depends on the response variable type (presence data, relative abundance data, etc.)<br>This choice of appropriate distance measure and/ or standardisation determines the degree of emphasis of sample richness, composition or relative abundances in the resulting distance matrix, and thus may also relate to the research question (Anderson et al. 2011). | Possible dissimilarity measures to compare samples should be selected using criteria listed in sources such as Anderson et al. (2011), Legendre & Legendre (2012) and Legendre & Cáceres (2013).<br>Principal coordinates analysis or non-metric multidimensional scaling (NMDS) of the different dissimilarity matrices can be compared. The distance measure that gives the greatest spread of points in a few dimensions and, therefore better discrimination among the samples, should be selected. If NMDS is used, the stress values associated with the different distance measures can be compared, because lower stress values show that the ecological distances between samples are being better represented in ordination spaces. |

#### a. Count the number of taxa, the number of samples and the number of empty samples and missing data points

The ratio of the number of taxa and samples is an important consideration because some multivariate methods are more susceptible than others to the influence of rare taxa (Lepš and Šmilauer 2003) and these may be of more or less interest, depending on the research hypotheses. Whilst long-term data and monitoring are highly valued (e.g., Hobbs et al. 2007), it is unfortunately common for long-term datasets to contain missing values; these values should be denoted as missing (i.e., ‘NA’ in R) and not given zero or other numerical values. The importance of missing values and how to deal with them during data analysis depend on the number of missing data points and how they are distributed across the dataset. Multivariate analyses based on compositional dissimilarities, e.g., some ordination methods, cannot be applied to datasets containing samples where no taxa are present (McCune and Grace 2002). However, empty samples may be important for analyses where we are interested in understanding responses to disturbance and local extinctions. The ratio of the number of explanatory variables to the number of independent replicates is also important for temporal analyses because it affects the degrees of freedom available for statistical testing (Johnstone and Titterington 2009).

#### b. Examine taxon frequency distributions and response curves

Some multivariate analyses assume specific frequency distributions of taxa (a histogram of the number of individuals or their relative abundance across samples for each taxon), e.g., CCA assumes that taxa have optima along environmental gradients (‘unimodal’ responses). Examining the shapes of taxon frequency distributions is necessary for determining whether taxa are responding strongly to environmental gradients in the dataset and whether their distributions conform to analysis assumptions. Where many taxa are involved, the common ones should be examined.

#### c. Determine the ratio of temporal to spatial variation in composition

The degree of spatial and temporal replication is important in determining the method of analysis because few sites with many measurements over time would be analysed differently from many sites measured only a few times. If the data have a nested structure, e.g., samples are nested within sites, this needs to be accounted for in the analysis, particularly if one wants to report significance values. For example, if plant species composition was recorded in quadrats that were nested within transects that were nested within sites, then quadrats are not fully independent replicates. Similarly, information from technical replicates (such as PCR products) should be kept separate at this stage because they provide useful information about variability at different scales in the dataset. It is inadvisable to average or pool data until the significant sources of variation are understood. For instance, if we want to test hypotheses of temporal dynamics, but there is a high degree of spatial variation among samples in the dataset, this can alter analysis choices or how results are presented and interpreted (Thrush et al. 2008). The relative amount of change in taxonomic composition can be estimated from the differences in sample scores on DCA ordination axes, e.g., the ‘gradient length’ statistic quantifies the turnover in taxonomic composition (Lepš and Šmilauer 2003); this relatively descriptive analysis would ideally be conducted as part of the EDA process prior to selecting an analysis pathway appropriate for testing a quantitative hypothesis of temporal dynamics, such as divergence or convergence.

#### d. Determine the optimal dissimilarity measure

A community dataset may be an abundance matrix (number of individuals), relative abundance matrix (e.g., percent cover, sequence abundances), rank-abundance matrix (ordinal abundance ranks), or a taxon-occurrence (presence-absence) matrix. These different response variables can influence the choice of analysis methods, including which distance measure to use to calculate pairwise community dissimilarity values for samples (see Magurran 2004, or Legendre and Legendre 2012 for a comprehensive description). To ensure that abundance data from different taxa are comparable using dissimilarity, data should be standardised prior to analysis if they are measured on different scales (Legendre and Legendre 2012). For example, taxon abundance values within each sample can be divided by the total abundance within the sample, or can be standardised across all samples so that each taxon has a mean of zero and standard deviation of one (Borcard et al. 2011). Alternatively, a distance measure that incorporates a standardisation can be used, e.g., Hellinger distance (Legendre and Gallagher 2001, Anderson et al. 2011, Legendre and Legendre 2012). The commonly-used Euclidean distance measure is not usually appropriate for taxon abundance data, which are often characterised by non-linear relationships among taxa, abundance values measured on different scales, or by the ‘double-zero problem’, which is the idea that samples that are missing the same taxa and therefore, have lower dissimilarity when using this distance measure (Borcard et al. 2011). Comparing results based on presence-absence data with those generated using relative abundance data allows us to evaluate the role of taxon abundances in generating temporal patterns (e.g., Angeler and Johnson 2012). Estimating abundances of each population within the community has its own important considerations, i.e., variation in taxon detection probabilities, which have been addressed elsewhere (e.g., animals: Gaston and McArdle 1994, metagenomics data: Amend et al. 2010). Many distance measures cannot be computed if there are empty samples or missing values. Deleting entire rows or estimating missing data so that the dataset can be analysed may be an option in some cases (McCune and Grace 2002). However, other measures, such as the Gower coefficient, can be calculated while ignoring comparisons among samples with missing data (Legendre and Legendre 2012; coefficient S15: 278).

### Step 2: Choose analyses for testing hypotheses of temporal dynamics

In this section, we describe a range of methods for testing hypotheses of community dynamics, including their outputs, highlighting in each case, the relevant ecological questions and the kinds of hypotheses that those analyses can be used to address: pairwise dissimilarity-based methods (section 1, below), zeta diversity (section 2), synchrony (section 3), turnover rates (section 4) and multivariate regression modelling (section 5). Further, each of the described methods has been cross-referenced with (i) which hypotheses it is best suited to test; (ii) whether or not it allows the identification of individual taxon effects on, or responses to, temporal change; (iii) whether or not explanatory and spatial variables can be explicitly incorporated; (iv) the type of statistical testing normally used; and (v) any requirements for the length of the time series (number of temporal samples) (Fig. 17). The selection of an appropriate analytical method also can be influenced by the kind of output desired (Table 2). For instance, some analyses result in outputs that require more effort to interpret than others, whilst others show differences in taxon responses or generate outputs that can be added to a geographic map. Further, as our literature review highlighted, in many cases several different analytical methods often are appropriate for asking questions about temporal dynamics and presenting the results at the desired level of complexity; it is also the case that many analyses can be used to test a wide range of hypotheses, so our focus is on describing the strengths of each analysis method and the minimum requirements for use, rather than on making explicit recommendations. Thus, we recommend that, at this point, readers use Tables 2 and 3 to help identify the methods of interest and then refer to the relevant sub-sections of section C that describe those particular methods.

**Fig. 17.**
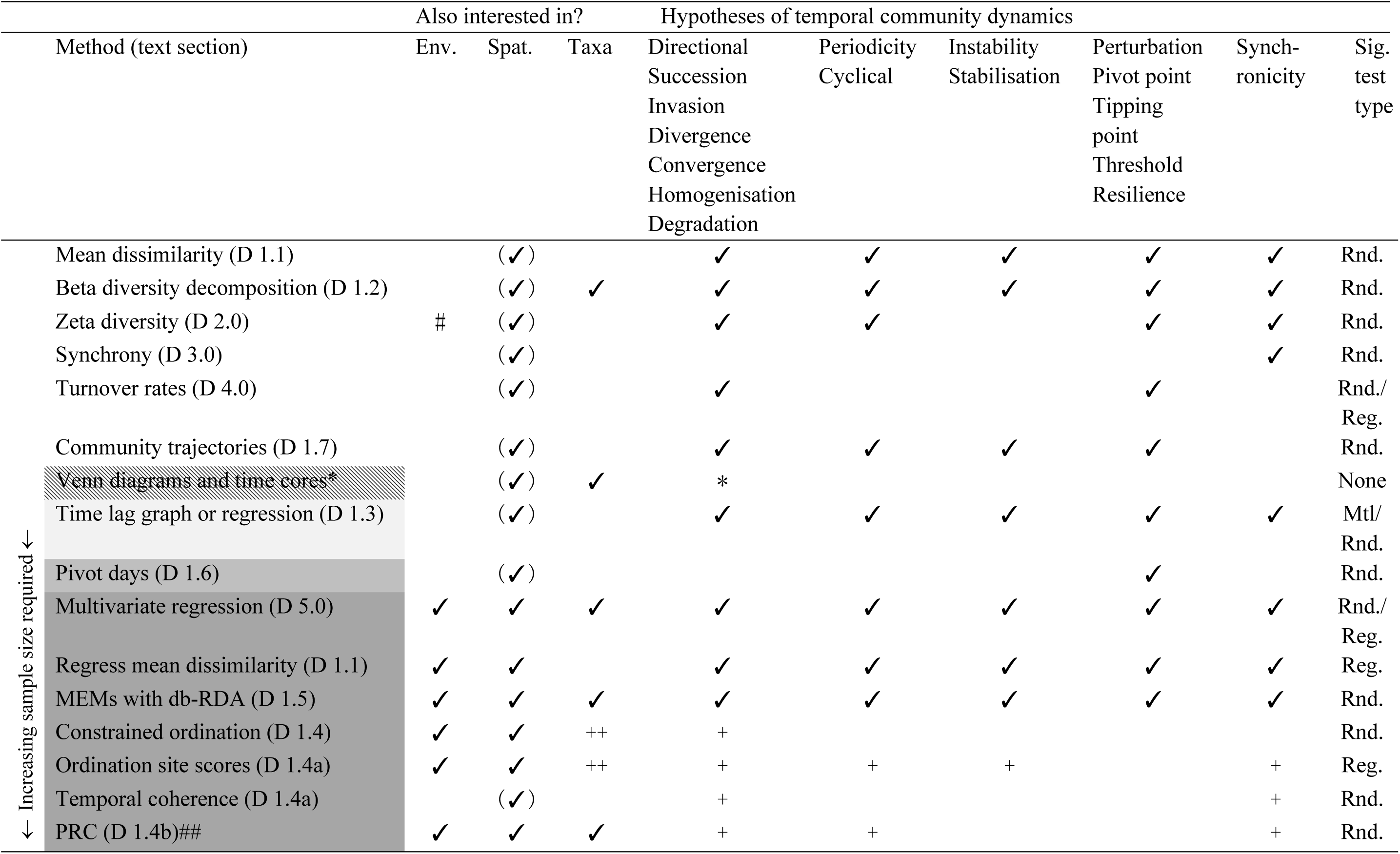
Potential methods that are appropriate for testing different predictions of temporal patterns in community dynamics indicating the required temporal sample size and whether: environmental variables can be included in the analysis (Env.); the analysis can be made explicitly spatial to account for temporal samples being clustered in different sites by including spatial variables (Spatial); or the role of individual taxa are identified in the analysis (Taxa), and the type(s) of significance testing (Sig.) that are used for the analysis (Rnd. = randomisations, Reg. = regression statistics, None = significance testing is not normally applied, Mtl. = Mantel test). Note that the null hypothesis for each of these predictions is that there is no change over time in either taxon presence or abundance (‘persistence’). Cell shading indicates the number of temporal samples in the time series for which each analysis method is appropriate (note that some hypotheses require more temporal samples than others): No shading—method applies well to time series of any length; cross-hatched pattern—method only applies well for few measures in a time series (< 5 temporal samples); Light grey—only applies well for ≥ 5 temporal samples; Mid-grey—only applies well for ≥ 10 samples; Darkest grey—only applies for ≥ 20 samples, but many more than 20 spatial and temporal replicates may be required, depending on the number of explanatory variables in the regression model; note that a time series of at least 20 temporal samples are required for calculating Moran’s eigenvector maps (MEMs). (✓) indicates that although these statistics are not explicitly spatiotemporal analyses, they can be calculated for spatiotemporal subsets of sample units and therefore, such groupings could be used to compare variation in both space and time. + Requires a strong gradient in composition that is well represented by the ordination axis scores for samples. ++ Weak inference from species’ positions on biplot. # Zeta diversity modelling can be combined with generalised dissimilarity modelling (Ferrier et al. 2007) to assess the effect of environmental variables. ## PRC requires a ‘baseline’ or a ‘control’ treatment level against which other samples are compared through time. Beta diversity decomposition refers to nestedness, turnover, and ‘Local Contributions to Beta Diversity’ (LCBD). * Venn diagrams and time cores are specifically useful for detecting taxon replacement over time, and are described in the section “Visualising temporal patterns”. Numbers in parentheses next to each method refer to the text section that gives a detailed description of each method.

**Table 2:**
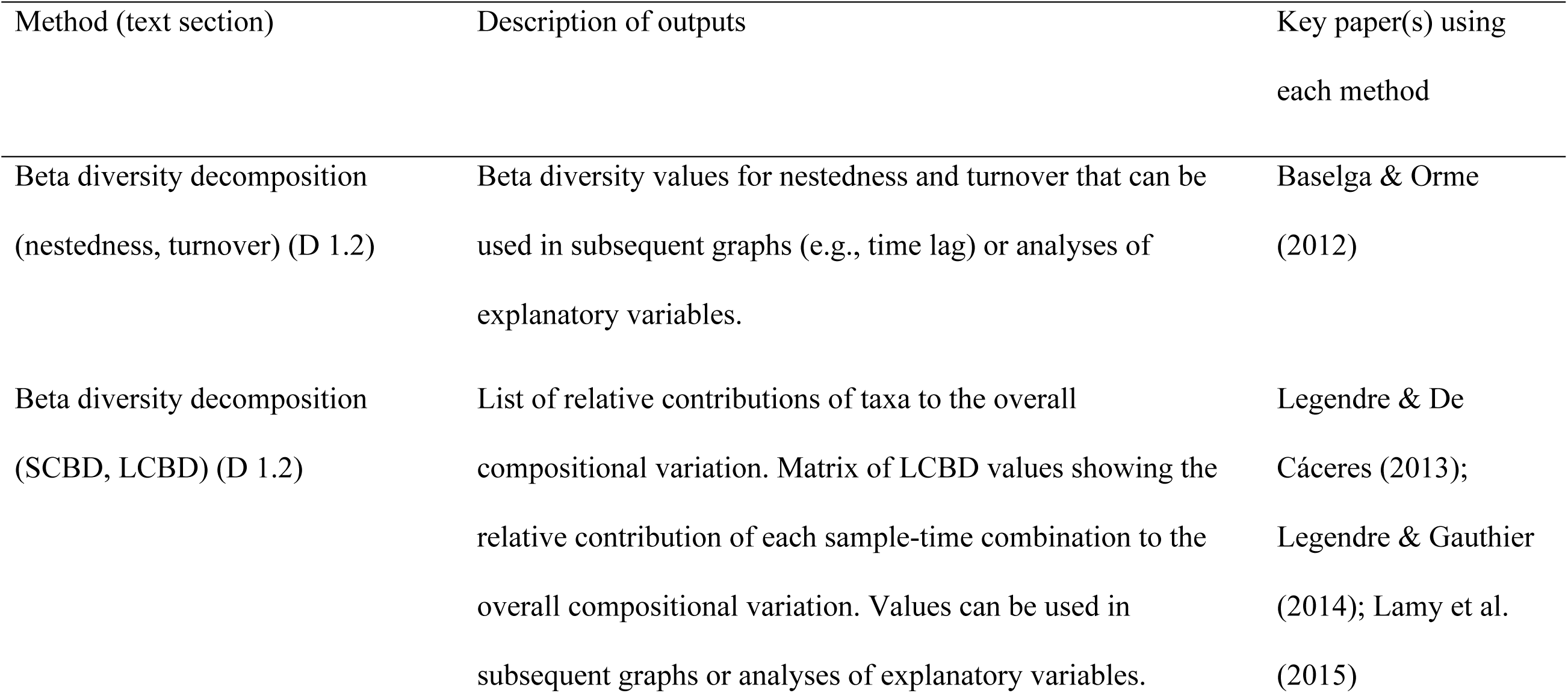
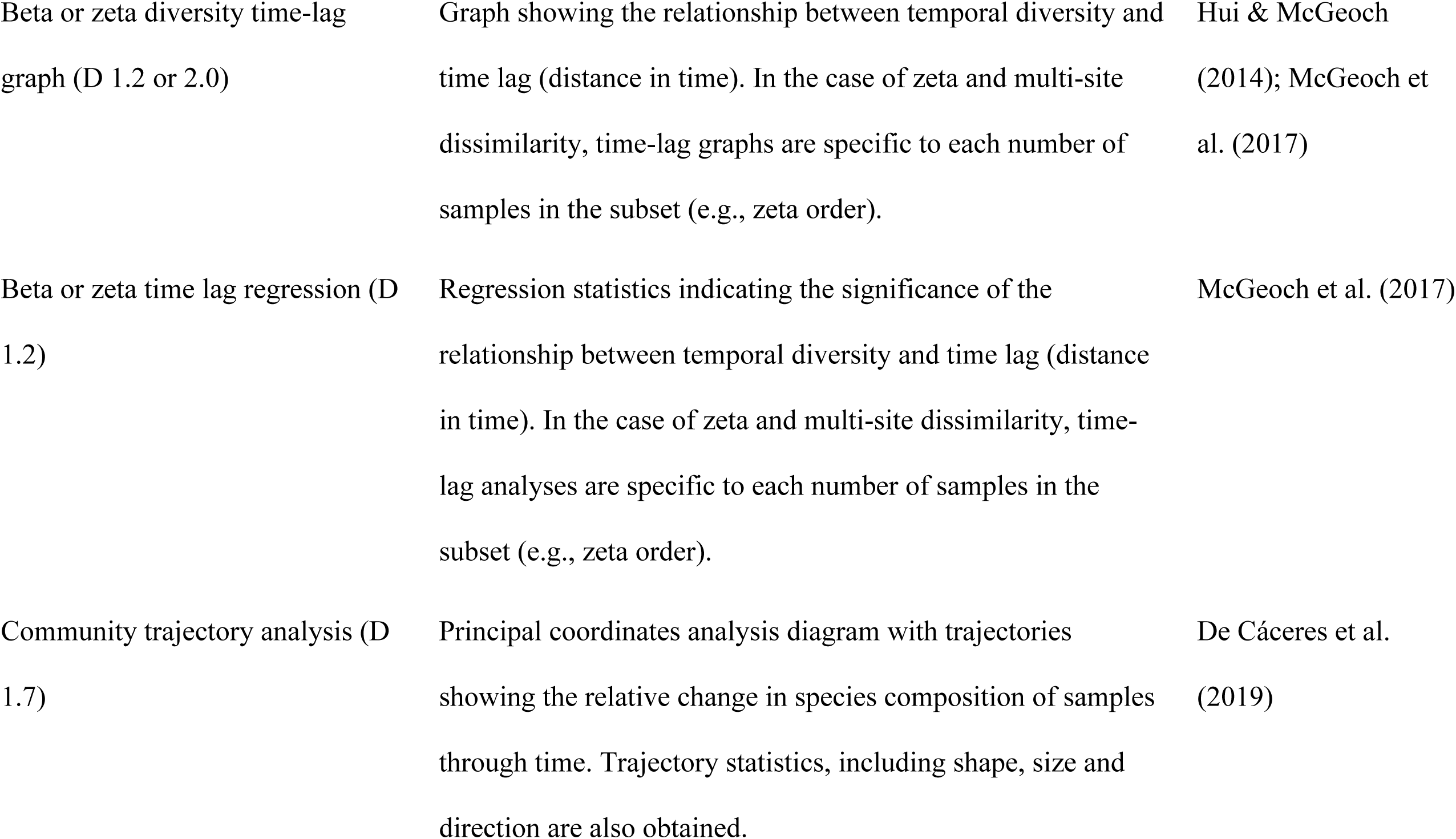
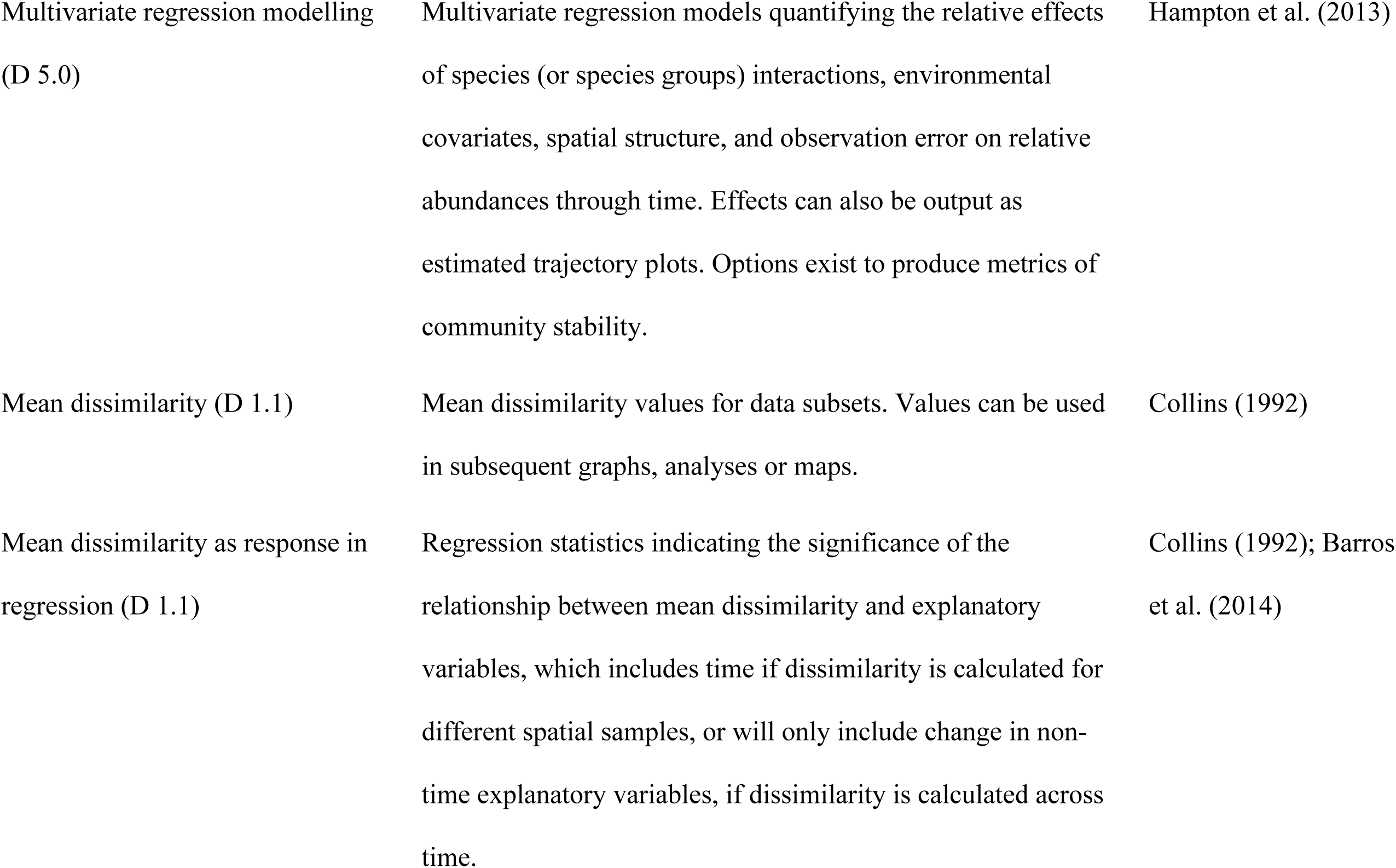
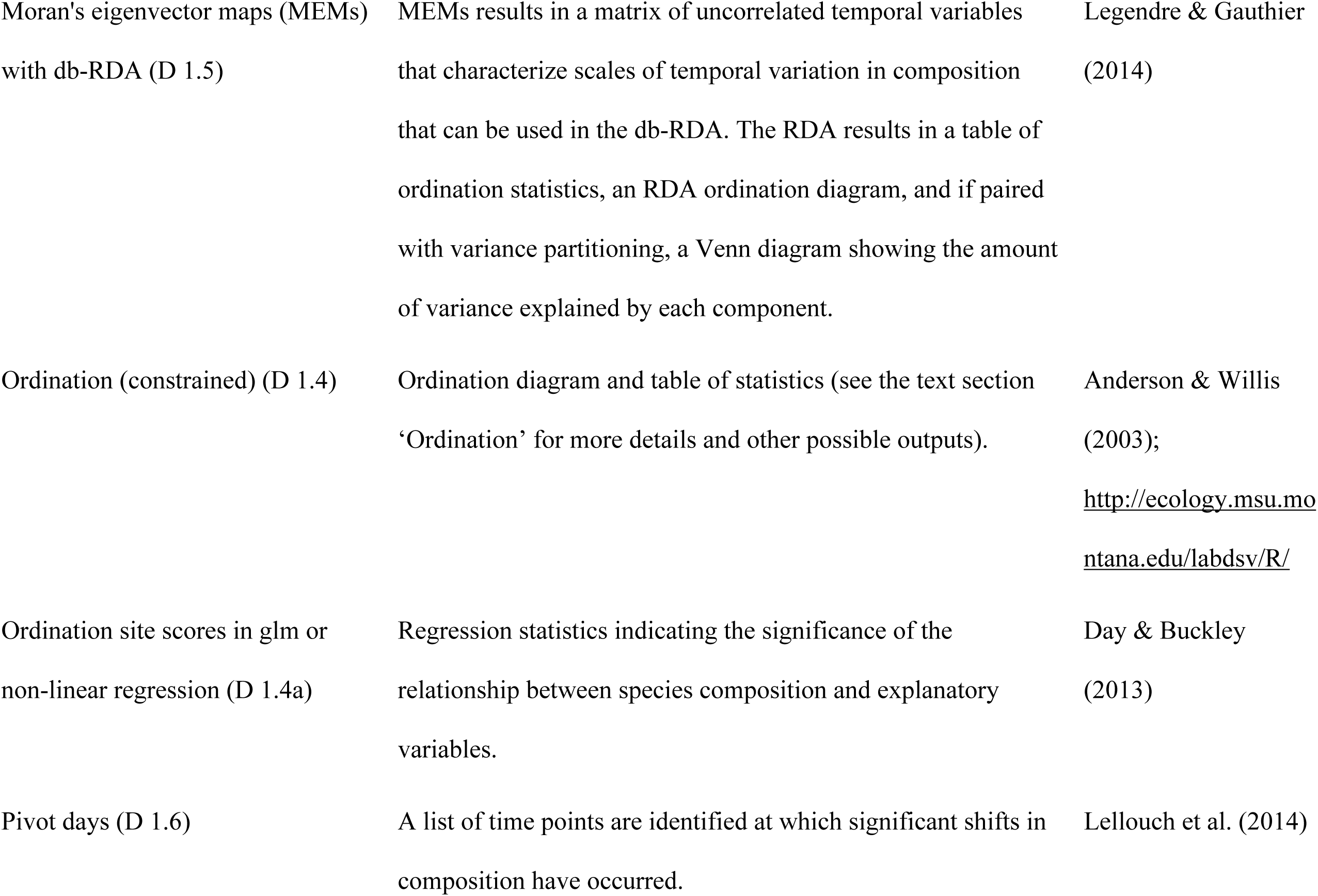
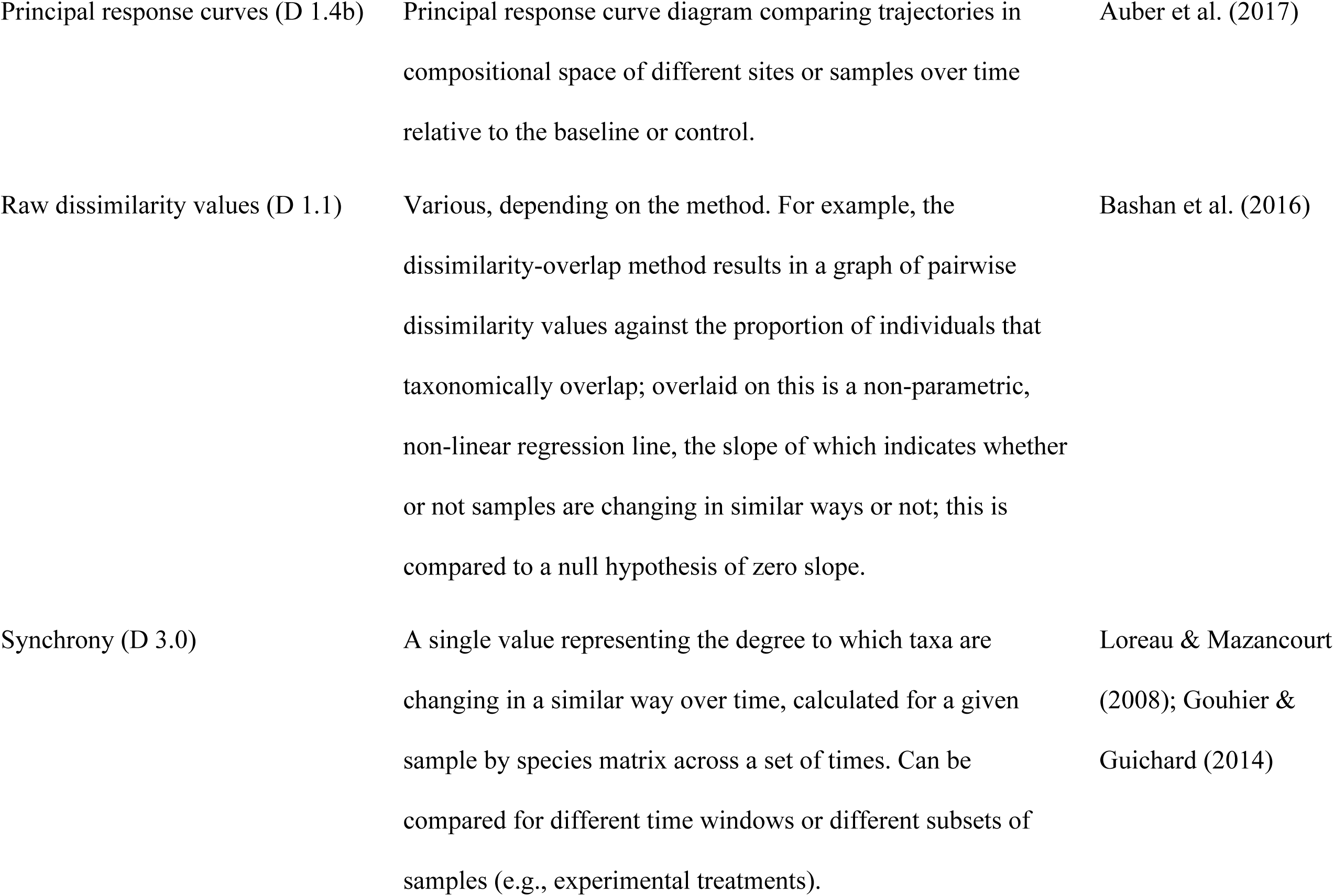
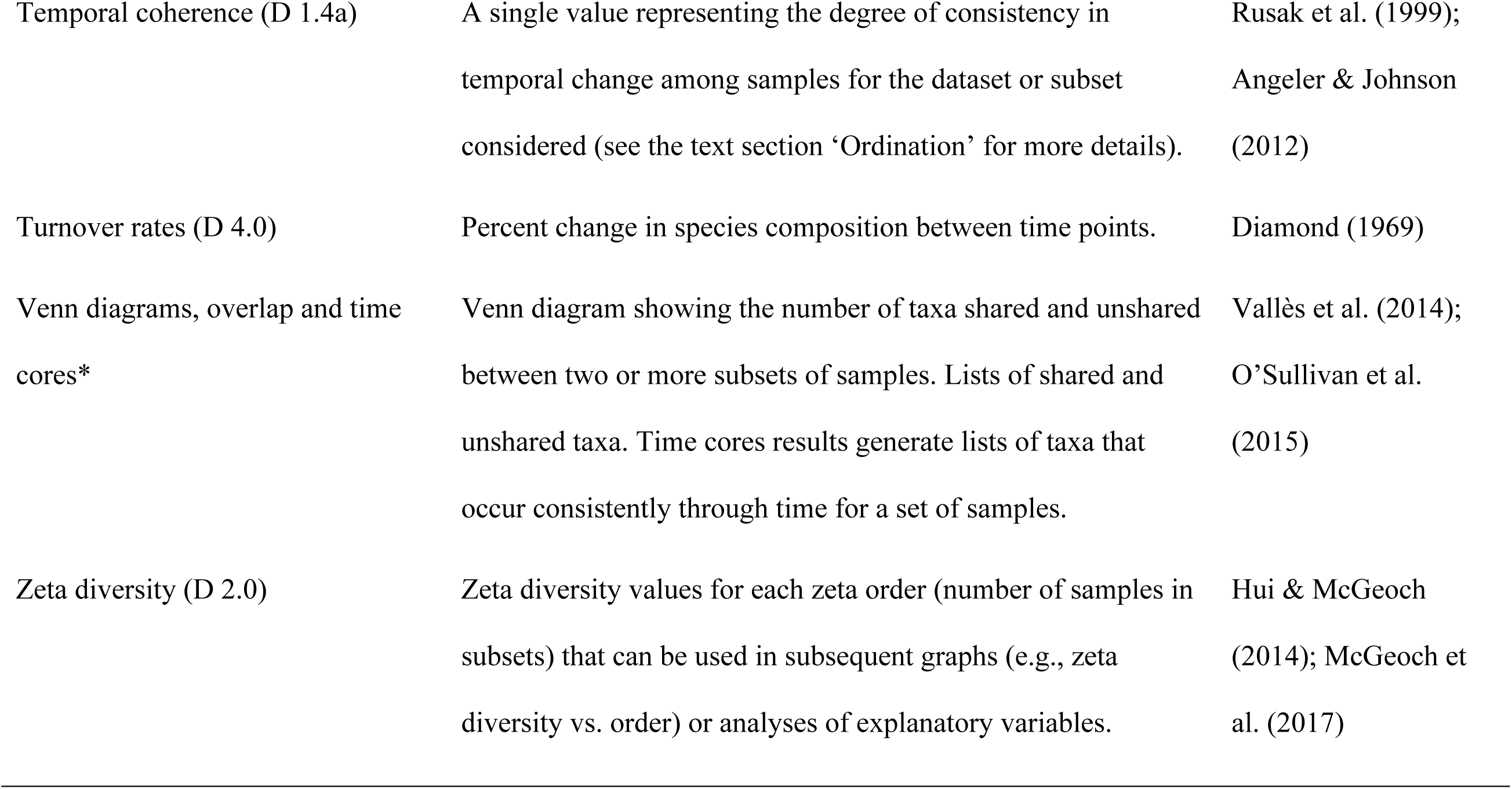
Descriptions of the outputs obtained from analysis methods for temporal change in community composition with example from the literature. LCBD refers to ‘Local Contributions to Beta Diversity’ and SCBD refers to ‘Species Contributions to Beta Diversity’. * Venn diagrams and time cores are specifically useful for detecting taxon replacement over time, and are described in the section “Visualising temporal patterns”.

**Table 3:** Data visualisation methods and considerations for their use in temporal community dynamics studies.

| Methods | Description and example references | Considerations |
| --- | --- | --- |
| Ordination axis<br>samples scores | Symbols (e.g., Noll et al. 2005), envelopes (e.g., Rayner et al. 2009), or trajectories (e.g., Rojas-Sandoval et al. 2012) can be added to ordination diagrams to display the relative compositional change among temporal samples. | Can be used to show convergence, divergence, or in/stability of sets of samples. Useful for many samples and taxa. Ordination sample scores can also be used in other graphs or maps to display spatiotemporal compositional variation. |
| Relative abundance<br>distributions | Plot the relative abundance of all taxa in the community (Lambshead et al. 1983, Stead et al. 2003, Ruhl 2008). | Shows changes in evenness over time where separate relative abundance distributions are plotted for each time. Useful only for small numbers of comparisons, e.g., two or three sampling times. |
| Venn diagrams | Displays the overlap in species composition among samples (e.g., O’Sullivan et al. 2015). | Suitable for visualising taxon replacement while specifically identifying the taxa driving compositional shifts. Useful only for small numbers of temporal replicates. |
| Time cores | Identify ‘core’ groups of taxa that remain present over time (Vallès et al. 2014). | Useful for visualising synchronicity in compositional shifts. |
| Heat maps | A grid representing spatial and/or temporal samples where cells are coloured by a gradient representing relative abundances, ordination sample scores, or compositional change (e.g., Pedersen et al. 2017, Pereira et al. 2017). | Useful for representing change between time points across sample space, even in the case of complex patterns. |
| Geographic maps | Grid cells or other geographic polygons displaying numeric values representing relative abundances, ordination sample scores, or compositional change (e.g., Clavero and Garcia-Berthou 2006). | Useful for representing change between time points where variation in geographic space is also important. Useful for relatively few time points. |
| Abundance graphs | Bar graphs, boxplots, scatterplots, or pie charts showing relative abundance of different taxa at different times (e.g., Xiong et al. 2014, Chapman and McEwan 2016, Franca et al. 2016). | Can show complex temporal dynamics such as nonlinearities in species' responses and synchronicity among different taxa, but useful only for a small number of taxa. |
| Relative abundance tables | Relative abundance or percent dominance values across time (e.g., Shertzer et al. 2009). | Useful only for a small number of taxa and spatiotemporal samples. |
| Photographs of gels or electropherograms | Show the relative frequencies of characteristic taxon identifiers (e.g., Shi et al. 2016). | Useful only for a small number of taxa and spatiotemporal samples. |

Our literature review revealed that in numerous cases, researchers performed and compared separate analyses on their community matrices from each sample time, e.g., separate nestedness analyses at each sample time (Azeria et al. 2006); we do not specifically address these applications here in any depth because these methods do not explicitly evaluate community dynamics through time. We also focus on methods that are truly multivariate rather than methods that are applied separately to abundances of individual taxa; however, if less than ten taxa are of interest, it may be that hypothesis testing via univariate approaches (e.g., separate linear regressions of taxon abundances) will be adequate for drawing synthetic conclusions about community change. In the supplementary material (Appendix S2), we summarise univariate methods that were commonly used in the literature review articles: univariate linear methods, temporal stability (*a.k.a*. coefficient of variation), frequency change, multiplicative change, and Markov chain modelling. A few studies used alternative methods such as cluster analysis and network analysis; however, we excluded those because we think the methods presented here provide equally good, or better, alternatives, but we nonetheless provide an explanation of them (Appendix S2) because they appeared in our literature review. Some of the analytical methods described below can be implemented in several different software packages (e.g., PC-ORD, PRIMER, CANOCO), but, for simplicity and widest accessibility, we illustrate their implementation in R (Data S2-4), except for Moran’s Eigenvector Maps (MEMs), which is given in Legendre & Gauthier (2014), and Multivariate Autoregressive State Space modelling (MARSS), which is presented in the documentation associated with the R package ‘MARSS’ (Holmes et al. 2012).

#### 1. Methods based on pairwise dissimilarity values

Pairwise dissimilarity methods can be used to test nearly the full range of questions and hypotheses in community dynamics (Anderson et al. 2011; Figs. 4-7). Pairwise dissimilarity methods are the most frequent way ecologists quantify the difference in the identities of taxa occurring in two samples. The result is a single number that measures the distance in compositional space between the two samples. Dissimilarity can also be represented as similarity, i.e., 1 − dissimilarity (Whittaker 1960, Anderson et al. 2011, Magurran 2013). When applied to multiple samples, it results in a matrix of all possible comparisons (a triangular dissimilarity matrix). For example, Collins (1992) measured the between-year percent change in plant species composition for grassland plots that had been repeatedly measured over a nine-year period. Dissimilarity methods are more desirable than univariate methods, which analyse the abundance of each taxon separately, because they consider the entire community at once and directly quantify change in community structure.

The development of dissimilarity indices is an active area of research. Magurran (2013) is a useful general discussion of dissimilarity measures, Anderson et al. (2011) provides a practical guide to selecting an appropriate index and issues of importance, such as whether to include joint absences in the calculation. Different authors often use different names for the same dissimilarity measure, so we recommend using a comprehensive and consistent guide, such as Legendre and Legendre (2012; Table 7.4: 324). Legendre and De Cáceres (2013) also provide comprehensive descriptions of most dissimilarity measures and rationales for choosing one. Several recent advances are specifically relevant to temporal data (e.g., Podani and Schmera 2011, Baselga 2012, Shimadzu et al. 2015) and some contributions aid in improving dissimilarity measures for use in certain circumstances, such as imperfect detection of taxa (e.g., Chao et al. 2006) or which indices are most appropriate for calculating dissimilarity (Legendre and De Cáceres 2013). For example, the unique fraction (UniFrac) dissimilarity metric was designed for microbial communities where phylogenetic information is available from sequencing that has been used to determine taxa in samples (Lozupone and Knight 2005, Lozupone et al. 2007). This metric measures phylogenetic distance between samples as the fraction of branch lengths leading to descendants from each sample. UniFrac can be used with abundance data, which weights the branch lengths prior to calculating distances (weighted UniFrac), or with presence-absence data (unweighted UniFrac Lozupone et al. 2007).

In the context of temporal community dynamics, dissimilarity is calculated in two different ways: (1) separately among spatial samples at each of a series of time points or (2) among samples taken at different times at the same location(s). Dissimilarity matrices are the basis for several methods recommended in our guide, including ordination analyses and Moran’s Eigenvector Maps (MEMs). It is important to realise that it is not easy to use such methods to simultaneously assess both temporal and spatial variation in the same analysis; however, temporal and spatial dissimilarity can be computed and assessed in parallel to test hypotheses and make inferences about spatiotemporal community dynamics (e.g., Legendre and Gauthier 2014, Lamy et al. 2015).

##### 1.1. Using the raw dissimilarity values

There are two ways in which raw dissimilarity values for samples are typically used. Either of these approaches could be used for different taxonomic subsets of the community to test hypotheses of taxonomic variation in dynamics. First, compositional dissimilarity can be calculated among samples taken at different locations to obtain a measure of spatial heterogeneity in community composition. To use these values to analyse temporal community dynamics, pairwise dissimilarity (spatial dissimilarity) must be calculated separately for each sampling time, and the means of these spatial dissimilarity values can then be compared across time visually in a table (e.g., Collins 1992), graph (e.g., Barros et al. 2014), or using a correlation or regression analysis where time is the predictor variable and amount of spatial variability at each site is the response (e.g., Barros et al. 2014). The mean dissimilarity can be divided by the number of taxa to generate an independent measure of compositional change (Cleland et al. 2013). Such comparisons over time could be done separately for different groups of samples, such as different experimental treatments, to test for predictors of differences in temporal change in composition. Bagchi et al. (2017) illustrate predictions of how temporal differences in dissimilarity can include non-linear patterns and thus how applying linear and non-linear regression modelling to pairwise dissimilarity values can be used to test more complex hypotheses of temporal dynamics. Regression approaches can also include mixed-effects models that account for nesting and other types of non-independence in the data, or non-linear models (e.g., generalised additive models) that may be useful to predict cyclical or threshold patterns in community dynamics (e.g., Shenhav et al. 2017). However, these modelling methods will apply well only in situations with longer time series (i.e., greater than 25 temporal replicates) and the more complex the prediction of temporal change, the greater replication is likely required. Alternatively, a Mantel test can be used to compare pairs of dissimilarity matrices computed at different times and to test for significant difference over time (see section in the main text ‘Test for statistical significance’). This approach is only useful for a small number of spatiotemporal observations (i.e., ten or fewer) because the comparisons are pairwise.

Second, if a set of samples were all taken at the same site at different times, ‘temporal heterogeneity’ can be calculated using the mean dissimilarity of the entire sequence of temporal samples to obtain a single measure of within-site temporal variability (Collins and Smith 2006). If multiple sites are sampled through time, this method can be applied at different spatial scales (Collins 1992, Collins and Xia 2015) to examine spatial scale-dependent temporal dynamics (Collins and Smith 2006). Regression approaches that relate mean temporal dissimilarity values to other explanatory variables for sites or samples could be used to explore spatial variability in temporal change. Alternatively, the ‘dissimilarity-overlap’ method, first described by Bashan et al. (2016) and further explained by Kalyuzhny and Shnerb (2017), can be applied to raw dissimilarity values to test for similarity among samples in community dynamics. This method uses the relationship between the pairwise dissimilarity values and the proportion of individuals in pairs of samples that overlap taxonomically to detect if changes in taxonomic composition (degree of overlap) alters relative abundances in a consistent way across samples (dissimilarities). It can be used for very large numbers of samples and taxa and could be applied to different groups of taxa to test for synchronicity.

##### 1.2. Beta diversity decomposition

‘Beta diversity’ is term used for several different types of measures in ecology, e.g., (Whittaker 1960, Baselga 2010, Legendre and De Cáceres 2013), including as a synonym for compositional dissimilarity. It always describes some measure of how taxonomic composition changes across a set of samples. A number of recent advances in the calculation of beta diversity have illustrated that, if calculated in particular ways, it can be ‘partitioned’ into different components that relate to different predictions of how taxa or their abundances change in space or time (Baselga 2010, Ellison 2010, Legendre and De Cáceres 2013). Beta diversity decomposition methods can be applied to testing almost any hypothesis of compositional change (Fig. 17). The outputs are focussed on species replacement and whether samples are subsets of each other (Table 2), so detecting patterns such as convergence or divergence is very easy. However, depending on what subsets of the data are used in separate calculations, beta diversity decomposition is equally able to detect rapid shifts in composition, e.g., perturbation, tipping point, or increases or decreases in temporal variability, e.g., ‘instability’ or ‘stability’. Two approaches so far have been usefully applied to quantifying community dynamics measured in the field (Baselga 2010, Baselga and Orme 2012, Legendre and De Cáceres 2013, Lamy et al. 2015).

First, when among-sample dissimilarity is measured as Sørensen’s dissimilarity [(*b* + *c*)/2*a* + *b* + *c*), where *a* is the number of shared presences and *b* and *c* are the number of unshared presences in each of the two samples], the Simpson’s dissimilarity index (also known as ‘species turnover’: *b* / (*b* + *a*)) can be subtracted from this value, resulting in a measure of ‘nestedness’ (Baselga 2010, 2012, Baselga and Orme 2012). This measure of nestedness reflects the degree to which the species composition of smaller samples (lower species richness) is encompassed by the species composition of larger samples (higher richness); samples may differ in space or time. In contrast, ‘species turnover’ measures the degree to which samples do not share species: how the identities of species differ among samples. It is useful to separate these two components (nestedness and turnover) to obtain a more informative quantification of beta diversity that reflects differences in the underlying processes. Such separation can be done either pairwise or for multiple sets of samples (Baselga 2010, 2017), and has been used to decompose temporal beta diversity in tests of hypotheses unique to temporal data (e.g., Baselga and Orme 2012).

Second, and in contrast, Legendre & De Cáceres (2013) defined beta diversity as the sums of squares in the variation of taxon abundance (or presence) corrected for the number of samples. They partitioned this measure of beta diversity into two different components: ‘Species contribution to beta diversity’ (SCBD) and ‘local contribution to beta diversity’ (LCBD), which describe, respectively, the relative contributions of different taxa and sites to overall beta diversity (Table 2; Legendre and Gauthier 2014). This method allows for tests of predictions of patterns in temporal community dynamics related to different taxa in the dataset. For example, Lamy et al. (2015) decomposed beta diversity (based on Bray-Curtis dissimilarity) of fish communities at each of 13 reef sites into contributions from biomass replacement and biomass differences over time. They demonstrated that total biomass in the community changed little, but that the observed significant temporal changes in fish composition resulted from differential changes in the relative biomass of species. Spatiotemporal patterns in the two components of fish community beta diversity were visualised using principal coordinates analysis (PCoA).

##### 1.3. Time-lag analysis

A time-lag analysis involves relating the amount change in composition to the amount of change in time across increasing temporal distances, called ‘lags’ (Table 2). For example, Collins & Xia (Collins and Xia 2015) compared plant species composition of repeatedly-measured grassland transects across all time lags between 1989-2008 (from a 1- to 19-year difference). There are two ways of assessing the time-lag effect on community dissimilarity: graphically (‘time-decay curve’) and statistically (time-lag regression analysis). For meaningful inference, time-lag graphs require at least five samples and statistical time-lag analysis requires at least ten samples for each temporal sequence. For both approaches, the pairwise compositional dissimilarity (temporal beta diversity) among temporal samples is calculated first.

To generate a time-decay curve, pairwise temporal dissimilarity values are plotted against temporal distance between samples (time lag; pairwise Euclidean distances in sample times) to show how community dissimilarity changes as samples become further apart in time (Collins 2000, Collins et al. 2000, Hallett et al. 2016). This graph can be then used to visually assess whether community composition is converging (becoming more similar) or diverging (becoming less similar) over time (e.g., Collins 2000). For example, Bar-Massada & Hadar (2017) plotted temporal beta diversity (as Bray-Curtis dissimilarity) of plants in grazed and ungrazed treatments in Mediterranean herb communities, revealing that composition diverged over time, regardless of grazing treatment. Alternatively, temporal community dissimilarity can be graphed against dissimilarity in environmental variables (rather than time), if the primary research focus is on response of the community to environmental change rather than time *per se*.

Time-lag regression analysis was developed specifically for data with greater than ten time points to assess directional, cyclic or stochastic changes within communities (Collins 2000, Collins et al. 2000) and has been evaluated against other temporal analysis methods (Angeler et al. 2009). For time-lag regression analysis, the pairwise compositional distances are plotted against the square-root of the time lag, which reduces the bias that results from having fewer data points at larger time lags (Collins 2000, Collins et al. 2000). A regression line is then fitted to determine the pattern of compositional change. A significantly positive relationship indicates that compositional changes in communities is directional, because sampling points that are further apart in time are more dissimilar than those that are closer together in time (i.e., Fig. 2 in Collins 2000). A negative slope shows that some convergence towards the taxonomic composition at one of the earlier time points has occurred, because measurements close together in time are more dissimilar in composition compared to those further apart in time. No slope, or a non-significant regression, implies stochastic changes in the community over time. Non-linear regressions also can be used to describe more complex temporal patterns (Magalhães et al. 2007).

A robust measure of significance in time-lag regression analysis can be determined by running Mantel tests on the dissimilarity matrix and the matrix of pairwise distances among samples in time, with Monte Carlo permutations (Bêche and Resh 2007, Day et al. 2017). Time-lag regression analysis is appealing because it is intuitive, produces one statistic (the regression coefficient), and can be easily visualised (e.g., freshwater macroinvertebrates: Bêche and Resh 2007, gut bacteria: Dimitriu et al. 2013, and plants: Collins and Xia 2015). However, time-lag analysis only provides one value for each sampling unit, so further analyses may be required to investigate changes within treatments or groups.

##### 1.4. Ordination

Ordination summarises multivariate community data by optimising relationships between high-dimensional samples and taxa in low-dimensional space to detect important ecological gradients in communities (Table 2). This dimensionality reduction means that they are not especially useful for directly testing, or visualising, complex changes in community composition over time, such as tipping points, cyclical patterns or synchronicity. However, more simple patterns, such as divergence or convergence, can be more easily detected. Ordinations are either unconstrained, meaning that they do not incorporate explanatory variables, or constrained, which assess the effect of explanatory variables, including time (e.g., Kent 2012, Legendre and Legendre 2012, Palmer 2019). Ordination methods, including constrained ordination and using the ordination scores from an unconstrained ordination in a regression model (e.g., general linear model, mixed-effects model) are only appropriate for testing hypotheses of community dynamics in cases where EDA shows that the ordination axes used strongly represent temporal compositional change and therefore explain a substantial proportion of the variation. This is because the amount of temporal compositional variation represented by the ordination axis scores will be low if there is not a strong temporal gradient along the axes. Ordination methods allow only weak inference of the relationship between taxa and community dynamics if the temporal pattern is complex relative to the ordination axes because some ordinations, e.g., DCA, only provide an ordination axis score for taxa, and others do not provide ‘species scores’ at all, e.g., PCoA and NMDS. PCoA can be used to calculate the amount of variance explained by the ordination axes, which is important if a time-series analysis is to be performed using the axis scores. Where species scores are not provided, weighted averaging can be used to plot points for each taxon on the ordination diagrams to relate differences in site composition to differences in taxon presence or relative abundance if you wish to examine the associations between samples and taxa (McCune and Grace 2002). One advantage of PCoA is that the first PCoA axis represents the gradient explaining the greatest amount of variation in composition, followed by the second axis that explains the next greatest amount, and so on; this is not the case for NMDS (McCune and Grace 2002). Therefore, it is recommended that NMDS scores are not used in community dynamics analyses unless the amount of variation explained by each axis is clearly stated in order to validate their use in further analyses. ‘Co-inertia analysis’ can be coupled with ordination in a ‘partial triadic analysis’ to investigate community dynamics in relation to environmental variables. This method may be particularly useful when testing for temporal stability (Kidé et al. 2015).

###### a. Using ordination site scores to assess compositional change

Ordination axis scores for samples taken at different times can be used to quantify and test hypotheses of compositional change. Ordination diagrams are visually appealing, but for illustrating compositional change through time, it is difficult to interpret the magnitude and significance of temporal dynamics without statistical testing, such as permutational analysis of variance (PERMANOVA: Anderson 2001), unless temporal trends in the data are very strong along the ordination gradients in composition. This difficulty is amplified if there are many samples or if the sampling design includes a large number of sites, environmental factors, or temporal replicates. Note that PERMANOVA requires the assumption of homogeneity of dispersion to be met (this can be tested using the “betadisper” function in the R package vegan; Oksanen et al. 2017). Further, ordination diagrams and their statistics computed for different matrices (different sets of taxa or sites) are not comparable; results of individual ordinations cannot be compared among them. The partial exception to this is Procrustes analysis, which can measure the distortion, or ‘lack of fit’ between two ordinations, or dissimilarity matrices (Jackson 1995, Peres-Neto and Jackson 2001). It can be used to identify which taxa are most important in differentiating between the two ordination patterns.

A more quantitative, and often more flexible, approach is to use the ordination axis ‘site’ scores in linear models with time and other explanatory variables of interest entering as predictor variables (e.g., Berthon et al. 2014, Broadway et al. 2015). For example, Cotton et al. (2015) used DCA scores as a response variable in a linear mixed model to investigate the importance of carbon dioxide, ozone, and time on arbuscular mycorrhizal fungal community composition. This is like doing a constrained ordination, such as RDA, dbRDA, or CCA, where time is used as a constraining explanatory variable in the ordination; these methods will test the significance of time and any other explanatory variables. However, it is more difficult within a constrained ordination to test for non-linear, cyclical or other complex temporal patterns, and in this case, using the ordination axis scores in a more complex model may be more useful.

Ordination axis scores for samples can be used to quantify temporal changes in other ways. For example, Quinn et al. (1991) developed a ‘malleability index’ by regressing the pairwise Euclidean distances between the first three dimensions of ordination space for temporal samples, against environmental variables. One potential problem with this method is caused by the tendency of ordinations to distort the relative distances (dissimilarity) between samples, depending on, for example, differences in the number of rare taxa, and so values from distance-based indices may not be comparable, even in different parts of the same ordination.

Temporal coherence measures the degree to which a response variable is similarly variable in time across a set of sites. In the case of measuring temporal coherence in community composition, the multivariate response variable is taxonomic composition in the form of ordination axis site scores (Angeler and Johnson 2012). Temporal coherence compares the amount of temporal variation in the ordination axis site scores across a set of sites by using statistics generated by an ANOVA of the site scores (using year and site as factors) to calculate the ‘intra-class correlation coefficient’, which ranges from −1/(*n−*1) to +1, where *n* is the number of sites sampled, given that *n* > 1 (for calculation details see Zar 1996 pp 398, Rusak et al. 1999). High positive values of this metric indicate increased coherence of the different sites in composition over time. The measure can be recalculated while dropping each time in sequence (an ordered jackknife) to determine the relative effect of each time on the overall value (Rusak et al. 1999). Different sets of sites or treatments can be compared using a Fisher’s *Z* test (Angeler and Johnson 2012) or randomisation test in the case of low sample sizes (Rusak et al. 1999).

###### b. Principal Response Curves (PRC)

The method of principal response curves (PRC) was first developed by van den Brink & ter Braak (1999) for comparing compositional change in treatment sites relative to a control site. It is similar to redundancy analysis (RDA) in that a multivariate regression is performed on the sample × taxon matrix constrained by variables of interest, but PRC is explicitly constrained by time (Lepš and Šmilauer 2003, ter Braak and Šmilauer 2015). It was developed to eliminate the need for cluttered biplots that make it difficult to see temporal trends within and between treatments. To overcome this problem, PRC uses a single axis, plotted against time, to express trajectories of compositional change (Lepš and Šmilauer 2003: 145); further axes can be calculated if they are of interest. Taxon weights also can be calculated and plotted for each PRC to show the change in the relative abundance (e.g., percent cover) of each taxon in the treatments relative to the control. The significance of the PRC is tested using Monte Carlo simulations (Van den Brink and Ter Braak 1999). As with RDA, PRC yields a measure of the percent variance explained for the overall taxonomic composition and how much is attributable to each variable (treatment and time). For example, Sugiura et al. (2013) used PRC to show that plant invasion in forests on a small island in the north-western Pacific had little effect on the temporal dynamics of invertebrate communities. The potential drawbacks of this method include its requirement for a ‘baseline’ or ‘control’ treatment level to be set for comparison, an attribute that many observational datasets will not have. Furthermore, PRC allows for only one ‘treatment’ variable and each analysis focusses on the change at one site, relative to the control site. However, Auber et al. (2017) have recently generalised this method so that it can be used to analyse compositional change at multiple sites, relative to a baseline or control, but this application removes the ability to consider an environmental or treatment effect. Thus, a principal response curve analysis yields a line graph showing the relative change in taxonomic composition for each ‘treatment’ along one compositional gradient relative to a control (Table 2).

##### 1.5. Moran’s eigenvector maps (MEMs)

Moran’s eigenvector maps (MEMs; also known as ‘principal coordinates of neighbour matrices’, PCNM) is a method designed to characterise the spatial scales of variation in univariate or multivariate data (Borcard and Legendre 2002, Dray et al. 2006, Legendre and Gauthier 2014). However, MEMs have recently been applied to detect temporal scales of variation in communities (Angeler et al. 2009, Caruso et al. 2013, Legendre and Gauthier 2014). For instance, if a cyclical pattern in compositional change, such as seasonality, was detected as significant for a dataset, this can be illustrated using line graphs showing the temporal pattern (Table 3; Legendre and Gauthier 2014). This method is particularly useful for detecting very complex temporal patterns, such as seasonality (periodicity), cyclical changes or tipping points, but requires relatively large temporal sample sizes (Fig. 17). For example, Kampichler et al. (2014) used MEMs to detect temporal patterns in multiple datasets that had at least 20 annual measurements of bird communities.

First, a distance matrix is generated. Then, to focus on neighbour relationships, a threshold value is chosen (see Borcard et al. 2011 for advice). For values below this threshold, the distance value is retained; values above this threshold are assigned an arbitrarily large value. Legendre & Gauthier (2014) recommend using one time interval as the threshold for regular time series or the length of the longest lag for irregular series. A PCoA is then applied to this modified distance matrix to generate a large number of eigenvectors that describe a wide range of temporal scales of variation.

The eigenvectors from the PCoA can be used as explanatory variables, most often in a redundancy analysis (RDA) or multiple regression. Combined with RDA, MEMs can be used to determine whether there is cyclic or synchronised temporal change in composition and which taxa, or taxon groups, are the drivers of it (Fig. 17). MEMs-RDA scores also can be plotted as a function of time to assess visually the intervals of fluctuating community dynamics (Angeler et al. 2009, Kampichler et al. 2014). In a comparison of different methods to assess community dynamics, Angeler et al. (2009) observed that MEMs-RDA consistently explained a greater proportion of variation in data with low taxonomic richness and linear changes in time relative to time-lag regression analysis, Mantel tests, or RDA. However, power was higher for taxon-rich communities because there was a greater probability of more taxa fitting the temporal models. As a scale-detection method, MEMs could be applied to determining the relevant time intervals for community monitoring. This method can be combined with variance partitioning to show the relative variance in composition explained by temporal, spatial and other explanatory variables (Legendre and Gauthier 2014).

##### 1.6. Compositional pivot days

Compositional pivot days is a dissimilarity-based method developed by, and unique to, Lellouch et al. (2014). This method is appropriate for situations where we want to test for the occurrence of fast changes in taxonomic composition across one point in time, such as the application of an experimental treatment or the effects of a rapid change in environmental conditions (Table 2). It works by comparing pairwise distance values for pairs of temporal observations taken at the same sample location to identify the time points where rapid shifts in composition occurred. A ‘pivot day’ is a sampled time point that is identified by pairwise similarities being low before and after that time point, but relatively higher across it. Compositional pivot days are identified either visually from the pairwise distance matrix plot, or using clustering (see Lellouch et al. 2014 for details). This method may be a good option for a preliminary identification of thresholds or rapid shifts in community structure, but does not provide a statistical test for the significance of the threshold relative to background. The power of this analysis will increase as the number of sampled time points increases relative to the number of pivot days.

##### 1.7. Community trajectory analysis

Community trajectory analysis quantitatively compares the temporal change in multivariate compositional dissimilarity space among samples using geometric calculations (de Caceres et al. (2019). These comparisons are based on the relative amounts, rates, and directions of sample movement in compositional space, thus, the method allows flexible testing of a wide range of very specific, quantitative hypotheses of change, such as the divergence or convergence of samples over time. This method is especially useful for a small number of spatial replicates because the output can be graphical; plots use arrows on an ordination diagram to show how samples have moved in compositional space over time; however, larger numbers of samples could be graphed separately. We do not illustrate this method here because comprehensive R code and explanations are presented in the R package ‘vegeclust’ (De Cáceres et al. 2010) and demonstrative examples are given in de Caceres et al. (2019).

#### 2. Zeta diversity

Zeta diversity, a measure of similarity, analyses change in community composition among samples in the dataset by calculating the average number of taxa that are shared between samples that are grouped in pairs, triples, quadruples, etc. (Table 3). The method was developed and introduced by Hui & McGeoch (2014), extended to temporal data by McGeoch et al. (2017), and comes with its own R package, ‘zetadiv’ (Latombe et al. 2017b). Zeta diversity estimates compositional turnover from multiple (*n* ≥ 2) samples, rather than the pairwise comparisons used to estimate beta diversity. What is obtained by the method is a series of zeta values (similarity) for different orders of zeta: zeta_1_ to zeta*_n_*, where *n* is the total number of time points. Each of these values represents the average of the number of taxa shared by each set of times in the dataset. For example, zeta_2_ (zeta order 2) represents the average number of taxa shared by all possible pairs of samples in the dataset (equivalent to 1 − beta diversity or mean similarity), zeta_3_ represents the average number of taxa shared by all possible subsets of three samples, and so on. Zeta_1_ (zeta order 1) represents the average alpha diversity (taxon richness) of all the samples in the dataset.

If there is complete turnover in community composition in time, at some point, zeta*_i_* (where *i* ≠ 1) will equal zero when there are no taxa shared by a given sized subset of samples. In contrast, if community composition never changes through time, zeta*_n_* = zeta_1_. Therefore, the rate of decline in zeta with increasing ‘zeta order’ tells us about the rate of change in community composition in time (‘zeta decline’). ‘Zeta decay’ can be considered analogous to time-lag graphs and time-lag regression analyses, which we described earlier for pairwise dissimilarity measures (Table 3). This means it experiences the same issue of few time points at higher lags, and thus higher uncertainty. The temporal scale(s) of community dynamics can be illustrated by plotting the change in sample zeta diversity of a particular zeta order against the temporal lag of samples, meaning that this method is useful for detecting both directional changes and cyclical patterns. For instance, if a multi-year time series in composition is cyclical (e.g., shows seasonality), zeta would seasonally decrease and then increase. These analyses can be done for different sets of samples to make spatial or temporal comparisons and, if necessary, zeta values can be normalised using the gamma diversity (total number of taxa in the dataset) to account for differences in taxon richness across samples (McGeoch et al. 2017). Further, different methods of creating the subsets for calculating zeta can be used, such as nearest neighbours or relative to a time-zero sample, further facilitating hypothesis testing (McGeoch et al. 2017). Latombe et al. (2017a) used zeta diversity to extend generalised dissimilarity modelling (Ferrier et al. 2007). This enables the user to assess taxonomic turnover at different orders of zeta and understand drivers of turnover in rare and common taxa (Latombe et al. 2017a). This method could be extended to assess temporal turnover at different orders of zeta by adding time as an environmental variable.

#### 3. Synchrony

Although other methods can be performed on taxonomic subsets to look for similarities and differences in the temporal dynamics among groups of taxa (“synchronicity”), a useful measure of the ‘temporal synchrony’ in multivariate taxon abundances was developed by Loreau & Mazancourt (2008) and comes with its own R package, ‘synchrony’ (Gouhier and Guichard 2014). This standardised value measures how closely changes in taxon abundances within a community track one another in time. It varies between zero (complete asynchrony of taxon abundances in the community) and one (perfect synchrony; all taxon abundances changing in the same way) and, by exploring different sized ‘time windows’, can be used to detect scales of temporal variation. Overall significance tests of synchronicity in community change can be obtained by performing null model analysis (e.g., Pedersen et al. 2017).

A similar method (Houlahan et al. 2007) uses the sum of the temporal variances and pairwise covariances of all taxa in the community to distinguish if the community shows ‘compensatory dynamics’ over time. This method results in either a positive, negative or zero community-level covariance value, indicating that taxa are increasing or decreasing in synchrony (positive community covariance), some taxa are increasing, but this is compensated for by decreases in other taxa (negative community covariance), or that no change is evidence (zero community covariance) (Houlahan et al. 2007, Hallett et al. 2016).

#### 4. Turnover Rates

The degree of temporal variation in community composition can be assessed by calculating the turnover rate of the community (a.k.a. ‘temporal turnover’ or ‘species turnover’), which is a measure of the rate of change in taxonomic composition for a ‘site’ over time (e.g., Aguirre et al. 2003). Turnover rates can be computed to measure the magnitude of compositional change, but in a way that does not result in an understanding of the direction of change or the taxa involved in compositional shifts. This method can detect when, and how rapidly, temporal shifts occur in a temporal sequence to detect complex temporal patterns such as perturbations, tipping points and pivot points (Fig. 17). Turnover rates are calculated using combinations of colonisation, immigration, extinction, mortality, recruitment, and survival. One of the commonly used methods is that of Diamond (1969) where the percentage turnover rate, *T*, is calculated as *T* = 100 × (*E* + *C*) / (*S*1 + *S*2), where *E* is the number of taxa that went extinct between two time points, *C* is the number of taxa that colonised between the same two time points, *S*1 is the number of taxa present at time 1 and *S*2 is the number of taxa present at time 2. *T* gives the percent change in taxon identities over the time period and varies from 0, representing no turnover, to 1, indicating complete turnover (Diamond 1969). This measure can be calculated only for a single location at two time points; pairwise turnovers must be calculated separately for datasets with multiple time periods or multiple sites. *T* values can be compared through time, at different time intervals, or at different spatial scales (e.g., Boulinier et al. 2001, Tworek 2004). They also can be used in a regression-style analysis to assess the effects of a treatment variable or other explanatory factor on compositional turnover (e.g., Aguirre et al. 2003, Tworek 2004). These methods are only useful for communities that have relatively high rates of temporal turnover in taxa. The degree of compositional turnover also is tied to the temporal extent of the study for the given system and the average lifespan of taxa. For example, complete turnover in prokaryotes can be observed over a period of days (e.g., Kara and Shade 2009), whilst many decades may be required to detect any species turnover in some plant communities (e.g., Kardol et al. 2010). However, because it can be standardised by the number of taxa in the community, *T* is a potentially comparable measure of compositional change among different datasets, if they are on similar timescales and encompass communities with a similar distribution of life-histories.

#### 5. Multivariate regression modelling

Multivariate autoregressive state-space models (MARSS) are multivariate linear regression methods that model the stochastic temporal dynamics in matrices of species’ abundances in relation to abiotic or biotic covariates, while accounting for potential observation (measurement) error (Hampton et al. 2013). The models are autoregressive in that previous measurements in time are assumed to be correlated to future measurement, and these autocorrelations are formalised by modelling the current values of the response matrix as a weighted linear sum of previous values at a specified temporal lag (Penny and Harrison 2006). MARSS models can be used to disentangle temporal variation in species abundances into the relative contributions of inter- and intraspecific interactions, abiotic variations, and of demographic stochasticity (Mutshinda et al. 2009). Effect sizes of these different components are estimated from the multivariate model structure through the use of conditional least-squares, maximum-likelihood, or Bayesian estimation methods (Hampton et al. 2013). Observation errors, and other complexities such as spatially-replicated time series and non-uniform sampling, can be formally modelled within the state-space framework (Holmes et al. 2012). Further, MARSS models can be used to compute several metrics that indicate the degree to which communities are possibly becoming less stable over time (Ives et al. 2003). This level of flexibility means that they are able to test for even the most complex temporal patterns in communities, and can simultaneously reveal similarities and differences among taxa, and account for environmental factors or other covariates. However, they do require relatively long time series (ideally more than 25 temporal replicates), and, like any statistical model, more independent replication is required for more complex models. There are several examples of the use of MARSS models in freshwater and marine ecology, particularly in terms of estimating aquatic food web dynamics (Beisner et al. 2003, Hampton et al. 2008, Thibaut et al. 2012); there have been relatively few applications for terrestrial communities (e.g., Yamamura et al. 2006, Mutshinda et al. 2009). The ‘MARSS’ R package (Holmes et al. 2012) enables the fitting of MARSS models and includes a range of ecological examples, so we have not included example code in this paper.

Multispecies N mixture models and joint species distribution models are other regression methods that can be used to model explicit, quantitative hypotheses of community change, with a focus on interactions among taxa (e.g., Dorazio et al. 2015, Ovaskainen et al. 2017). These methods can allow for useful additions, such as accounting for uncertainty in the detection of taxa; however, due to their complexity and high computational requirements, they have yet to be implemented for datasets with large numbers of species.

### Step 3: Test for statistical significance

Whichever multivariate analysis has been selected, most ecologists interested in community dynamics seek a way to determine if the changes that they see over time in the results of those analyses are greater than, or less than, they would expect by chance alone. Some of the methods presented above have their own built-in significance tests, often based on randomisations of the community matrix. Commonly-used randomisation tests include the Mantel test (Legendre and Legendre 2012) and permutational multivariate analysis of variance (PERMANOVA; Anderson 2001, Legendre and Legendre 2012). Null models, whereby community matrices are randomised under sensible criteria (Gotelli and Graves 1996, Gotelli 2000, 2001, Gotelli and Entsminger 2001, Ulrich and Gotelli 2007, 2010), can be used to test the significance of any observed change in compositional dissimilarity, and can easily be incorporated into analyses described above including zeta diversity. The primary consideration in applying any randomisation test is what assumptions are being made about taxon niches and whether they are valid. If the interest is in testing the effects of interactions among taxa on their co-occurrence, but the randomisation procedure (null model) allows taxa to occur in samples where they are unable to occur for other reasons, such as habitat differences, a difference among samples may be falsely detected (an inflated Type I error rate). Thus, if strong inferences are to be made about compositional differences, we must be sure that the differences among the groups of sites that we are testing are the only difference. Similarly, the sample size needs to be high enough to allow the detection of differences among sets of sites for the given level of compositional variability (Type II error rate).

Schaefer, et al. (2005) introduced a novel method for detecting significant shifts in community composition using randomisations under null models of taxon mean abundances through time. They demonstrate that this approach, which uses both compositional distance (pairwise dissimilarities) and the coefficient of variation (CV), could detect significant dynamics in community composition, even where only small amounts of change had occurred. In this method, temporal changes in communities are assessed by using a compositional distance measure that is selected by the user. This quantifies the multivariate distance between the randomly generated communities (null model randomisations) and observed ‘reference’ samples (e.g., ‘Time 0’ samples, compared to randomisations of ‘Time 1’ samples). If the observed distance is greater than under randomisation, then we can infer statistically significant temporal change. Because this can be done in an ordination context, increases and decreases of particular taxa driving the change can be determined from the ordination plots. Limitations of this method are that it requires a relatively large degree of temporal change in the data compared to spatial variation (i.e., more temporal, than spatial, variation) and it is relatively computationally intensive; however, we provide the R code for a worked example in the supplementary material (Data S2).

### Step 4: Report on the analysis workflow

Our literature review suggested a lack of consistent reporting of critical aspects of the data analysis workflow. Such reporting of ensures that the results are robust, represent the variation deemed important for the research aim and are reproducible. The number of taxa, samples, and independent replicates in space and time all should be clearly reported in the Methods or Results section of the paper (see Hurlbert 1984 for guidance on statistical independence). The dissimilarity measure selected, and the rationale and procedures used to arrive at that decision, should be reported in the Methods section of the paper. We encourage researchers to refer to a specific source, such as Legendre & Legendre (2012), or to give the formula applied to maximise the repeatability and transparency of their research; these indices can be inherent in particular methods or built-in as defaults in software, but nevertheless should still be reported. Different methods of dissimilarity or ordination (or other analysis types) that were compared, e.g., in the EDA phase, should be listed in the Methods section of the paper stating that their results were consistent with each other. The choice of dissimilarity or ordination method is subjective, but rules of thumb can be used for assessing different methods, so that the method selected optimises the reduction of dimensionality in the data and thus maximises the variation accounted for by the analysis (see McCune and Grace 2002 for comprehensive advice).

### Step 5: Present temporal patterns and results

Some of the analysis methods that we describe in this paper do not produce simple visual outputs that clearly show temporal trends in community datasets and visualising temporal dynamics can be difficult for species-rich datasets or complex sampling designs. However, there are a variety of useful methods that can be used to complement a hypothesis-driven analysis (Table 3). Many descriptive methods work best where a small group of taxa is measured only a few times. However, there are examples of descriptive methods being used effectively for larger, more complex examples. Pereira et al. (2017) used a heat map to show compositional changes of 808 bacterial genera, at a single location, sampled 24 times over two years. Certain descriptive methods are useful for only specific types of temporal patterns, e.g., ‘time cores’ is a method that identifies ‘core’ groups of taxa that remain present over time (Vallès et al. 2014). Others, such as Venn diagrams and measures of overlap in composition, are useful only for small numbers of temporal comparisons. We recommend and provide R code (Data S2) for time cores and Venn diagrams because these methods are suitable for assessing hypotheses of taxon replacement while specifically identifying the taxa driving compositional shifts (Tables 2 and 3). Relative abundance distributions (RADs, *a.k.a*. ‘rank abundance curves’), can provide relatively sophisticated visualisations of community dynamics, and rank abundances can be used to quantitatively measure the shift in taxon ranks within samples taken at different time points (Hallett et al. 2016). RADs are plots of the relative abundances (on the *y*-axis) of all species in a community ranked (on the *x*-axis) from highest to lowest. They are closely related to *k*-dominance curves, in which species ranks are shown on the *x*-axis, but cumulative proportional abundance (dominance) is plotted on the *y*-axis (Lambshead et al. 1983). If samples taken at different times are plotted separately, the set of RADs can illustrate changes in community structure over time. RADs also can be used to explore community dynamics if a dissimilarity measure, such as Bray-Curtis, is calculated for the species’ ranks at each time point. The resulting temporal dissimilarity of the RADs can be explored by plotting rank dissimilarity against time lag or by ordinating rank dissimilarities in the same way that spatial compositional dissimilarities are examined. For example, temporal changes in RAD dissimilarity can illustrate changes in species evenness over time as might be expected if compositional homogenisation and a decline in evenness occurs because of invasion by an increasingly dominant species (Ruhl 2008).

## E. DISCUSSION

There is an ever-expanding set of analyses that use a wide variety of multivariate and univariate methods to quantitatively address temporal questions about communities that continue to be developed (e.g., De Cáceres et al. 2019). One of the key reasons for the proliferation of methods appears to be variability in the length of the time series. For example, if the time series is long enough, methods such as temporal MEMs could be used; however, consistent with other reviews (e.g., Vaughn and Young 2010) and databases (Dornelas et al. 2018), most datasets captured by our literature had relatively few temporal measurements (mean = 14.9, median = 7, mode = 4), requiring very different methods. In other cases, the ability to implement some methods in temporal community ecology still requires development, such as the application of time-ordered and time-aggregated network analysis for spatiotemporal community datasets (Blonder et al. 2012).

Some analysis types are well established for spatial data, but can be applied easily to temporal data, potentially yielding useful and comparable results for explicit temporal analyses. Further research is required to generate standard extensions and approaches using such methods for temporal data. For example, nestedness (*sensu* Patterson and Atmar 1986) is usually applied to spatial data and the analysis is repeated for different time points; however, there is no reason that we cannot apply this method across times to look at whether sites are becoming more or less nested over time. For instance, if younger sites were nested within older sites, this may indicate that sites were becoming more homogeneous over time.

Our understanding of temporal dynamics in biological communities, and the methods we use to investigate them, have improved greatly over the past 25 years, probably because of increases in computational power and in dataset size. We can continue to improve our understanding by moving away from merely describing community changes, to linking these changes to understanding and testing the processes driving community change and biodiversity; that is, moving from hypothesis-generating methods towards hypothesis-testing methods enabled by well-defined research questions with specific, testable hypotheses that result in clear predictions for observed temporal patterns (e.g., Figs 4-7). Even for exploratory studies of compositional variation through time, we emphasise the benefits of making *a priori* predictions for how we think composition should change through time.

Interactions between community composition and environmental or other factors across space and time are complicated. For example, Brown (2003) showed that temporal variability in species composition of stream invertebrates was reduced by high spatial heterogeneity, which may be because there are increased niches or refugia in changing environmental conditions. Manipulative experiments are useful, but not always possible, especially for long-lived taxa. Therefore, combining current knowledge of intrinsic and extrinsic factors controlling community change into null models and simulations may be the way forward (Brown and Lawson 2010). Carefully considering sample sizes and appropriate replication of particular factors are also important if we are to properly test hypotheses of species coexistence over time (Philippi et al. 1998). There is a need for us to make useful predictions about how communities may continue to respond to ecosystem disturbances, such as climate change, invasive species, and habitat fragmentation (Hobbs et al. 2007, Kampichler et al. 2012). Many methods commonly used to analyse temporal community dynamics are not quantitative, meaning conclusions can only be narrative descriptions, rather than clear tests of hypotheses. Many of the methods presented here push us to quantitatively assess dynamics, which naturally leads to clearer hypothesis testing.

There is a wide variety of methods available for analysing temporal dynamics of biological communities; however, many studies in our literature review using these methods are descriptive, showing that a large body of literature in community ecology is not focussed on testing quantitative hypotheses of change. Surprisingly few studies in our review presented hypotheses that were more complex than simply predicting that composition would have changed in some way over time, instead of specifying predictions of quantitative and directional patterns of change in community composition. Even additional searching beyond our original search terms that included more specific terminology for compositional change (succession, turnover, synchron*, resilience, homogenization, divergence, convergence) revealed few empirical papers that tested specific, quantitative hypotheses. This search resulted in an additional 1,514 papers of which we read 10% (151 papers). Of these, 60 papers met the inclusion criteria of our original review and only 13 papers tested hypotheses that were more sophisticated than the simple prediction that composition would have changed in some way over time, e.g., directional change or that the abundances of some taxa would change more than others.

This lack of specific, quantitative hypothesis testing is not surprising because to predict change in composition in a quantitative way, we require an in-depth understanding of a system and how different components of the diversity differ in their ecology or other factors that could cause directional, cyclical, or other differential change in time. In many cases, particularly in the observational, rather than experimental, community ecology literature, this is not the situation. This means that, although such studies may state hypotheses, these hypotheses are not very specific or quantitative. In most cases, they predict some change in response to time or another factor that is correlated with time (such as disturbance or pollution), but the types of temporal change (divergence, convergence, synchrony, etc.) are not specified. We strongly recommend that researchers generate *a priori* hypotheses of compositional change (e.g., Figs. 4-7; Bagchi et al. 2017). These predictions should be used to select appropriate quantitative data analysis methods (Fig. 17). The comprehensive description and hypothesis testing of temporal community patterns requires the use of multiple data analysis methods, but by using standard exploratory data analysis methods followed by testing *a priori* hypotheses, the methods selected can be explicitly focussed on the most appropriate tests for the situation. We hope that this synthesis will allow researchers to make informed decisions about their choice of data analysis for their research question and pave the way for more uniformity in reporting of these analyses and/or innovation in comparability. Our understanding of patterns and drivers of changes in species communities would benefit by having more longer-term and fine-scale studies.

## ACKNOWLEDGEMENTS

We thank Audrey Lustig for initial database and R code development. This paper is a contribution from the Harvard Forest LTER program. We thank members of the Forest Ecology Research Group at Wilfrid Laurier University, Glenda Mendieta-Leiva, Brian Inouye, Miguel De Cáceres, and an anonymous reviewer for comments on the manuscript. We thank the authors of Porensky et al. (2016a, 2016b) and Gilbert et al. (2012) for making their data publicly available. The BCI forest dynamics research project was founded by S.P. Hubbell and R.B. Foster and is now managed by R. Condit, S. Lao, and R. Perez under the Center for Tropical Forest Science and the Smithsonian Tropical Research in Panama. Numerous organizations have provided funding, principally the U.S. National Science Foundation, and hundreds of field workers have contributed.

## SUPPORTING INFORMATION

Data S1. Description of attributes, and associated data, extracted from papers in the literature review.

Data S2. Example R code for plant community dataset Data S3. Example R code for BCI dataset

Data S4. Example R code for marine bacterial community dataset (from high throughput sequencing)

Appendix S1. Supplementary figures from the literature review.

Appendix S2. Descriptions of methods encountered in the literature review that are not included in the main text.

## Appendix S1

**Figure S1.**
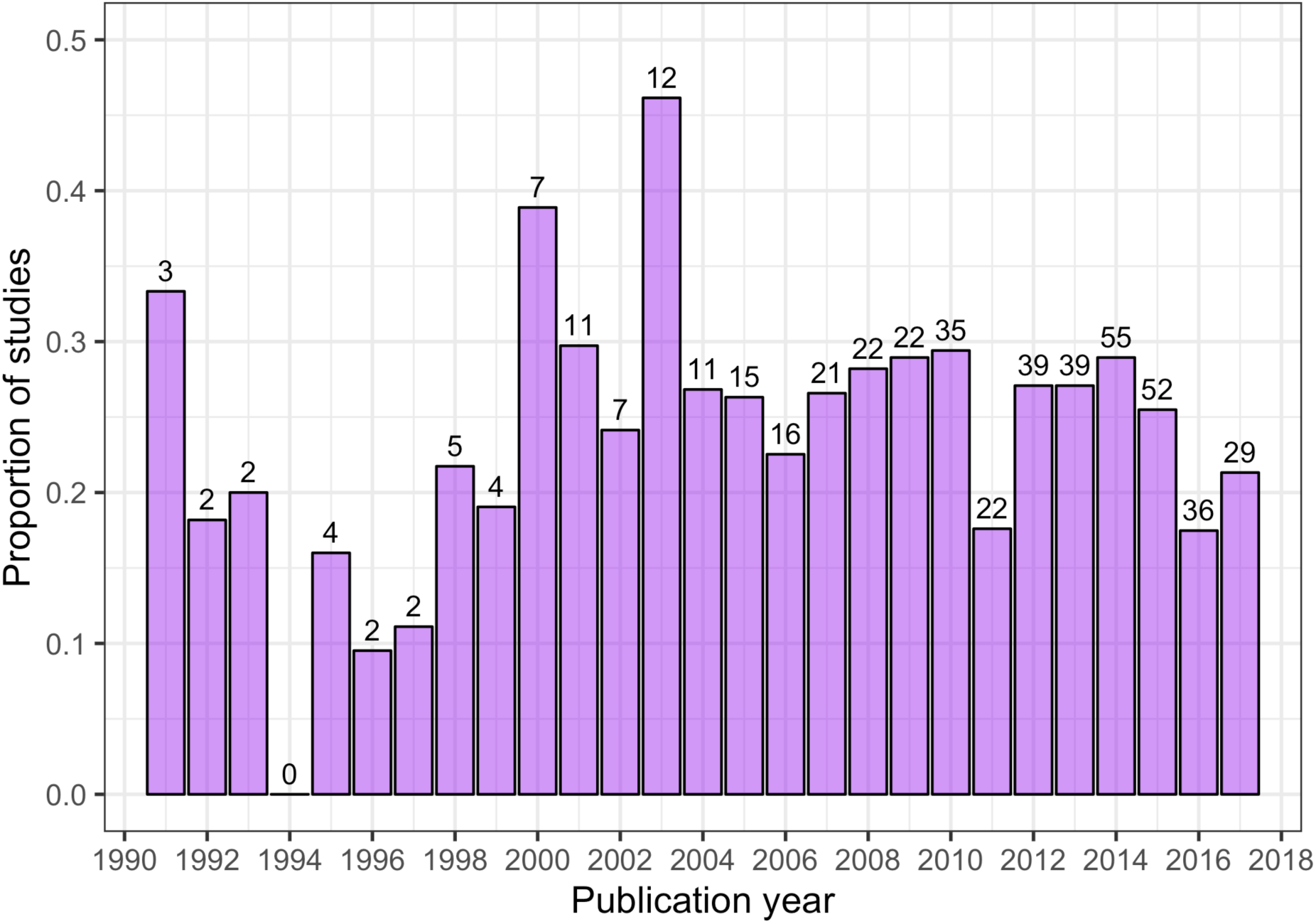
Proportion of studies identified using our search terms that were published each year until September 2017. No studies prior to 1991 were captured by our search terms. The total number of studies in each year is the value above the bars.

**Figure S2.**
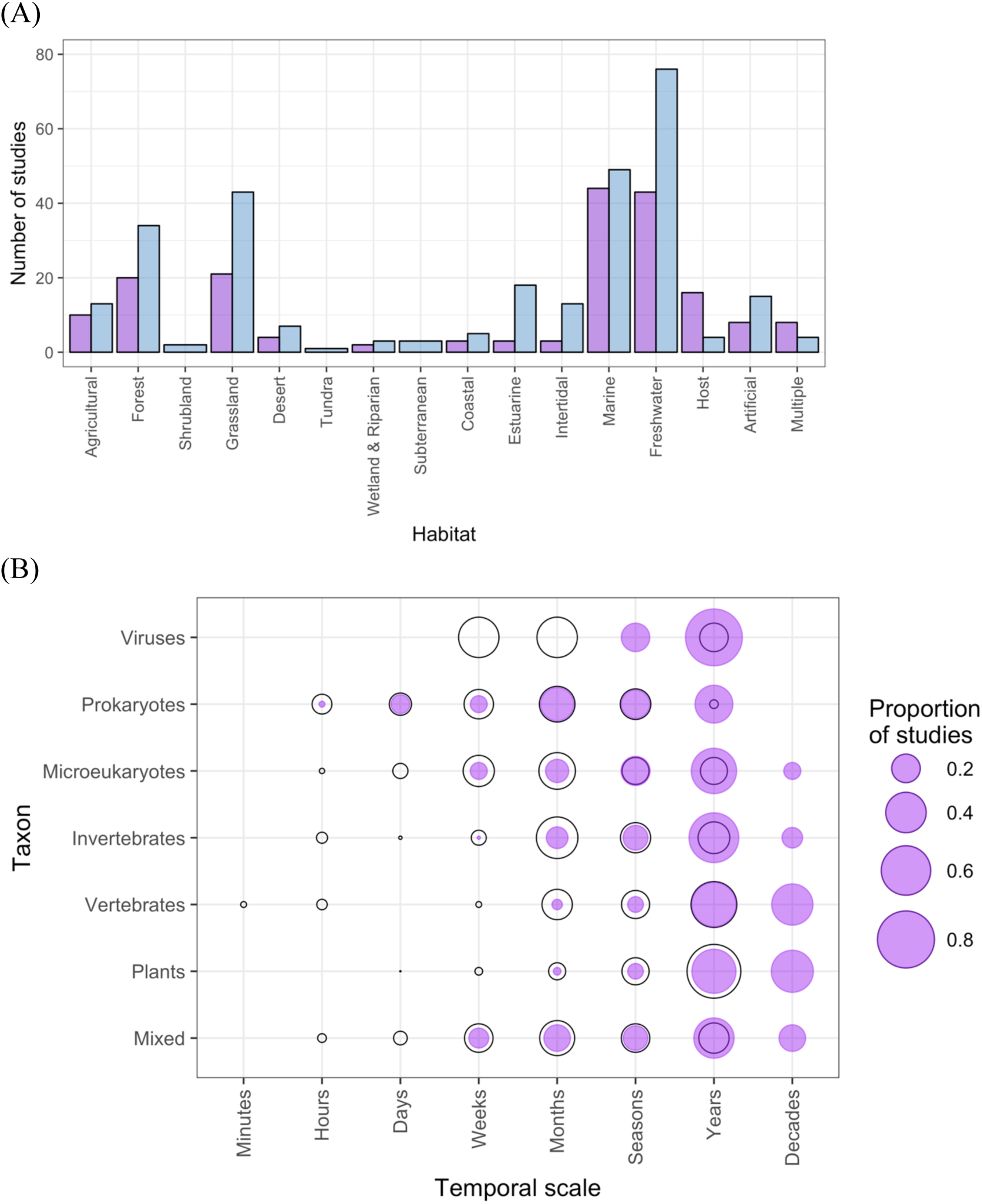
(A) The numbers of small and large spatial scale studies by habitat category. Small spatial scale (blue bars) studies were considered to be those carried out at micro, point sample, and local scales, while large spatial scale studies (purple bars) were those at regional, continental, or global scales. (B) The proportion of studies of different taxa within categories defined by the temporal grain (black circles) and extent (purple dots). Temporal grain is the minimum time between sampling events within the study and the temporal extent is the maximum time between sampling events within the study.

**Figure S3.**
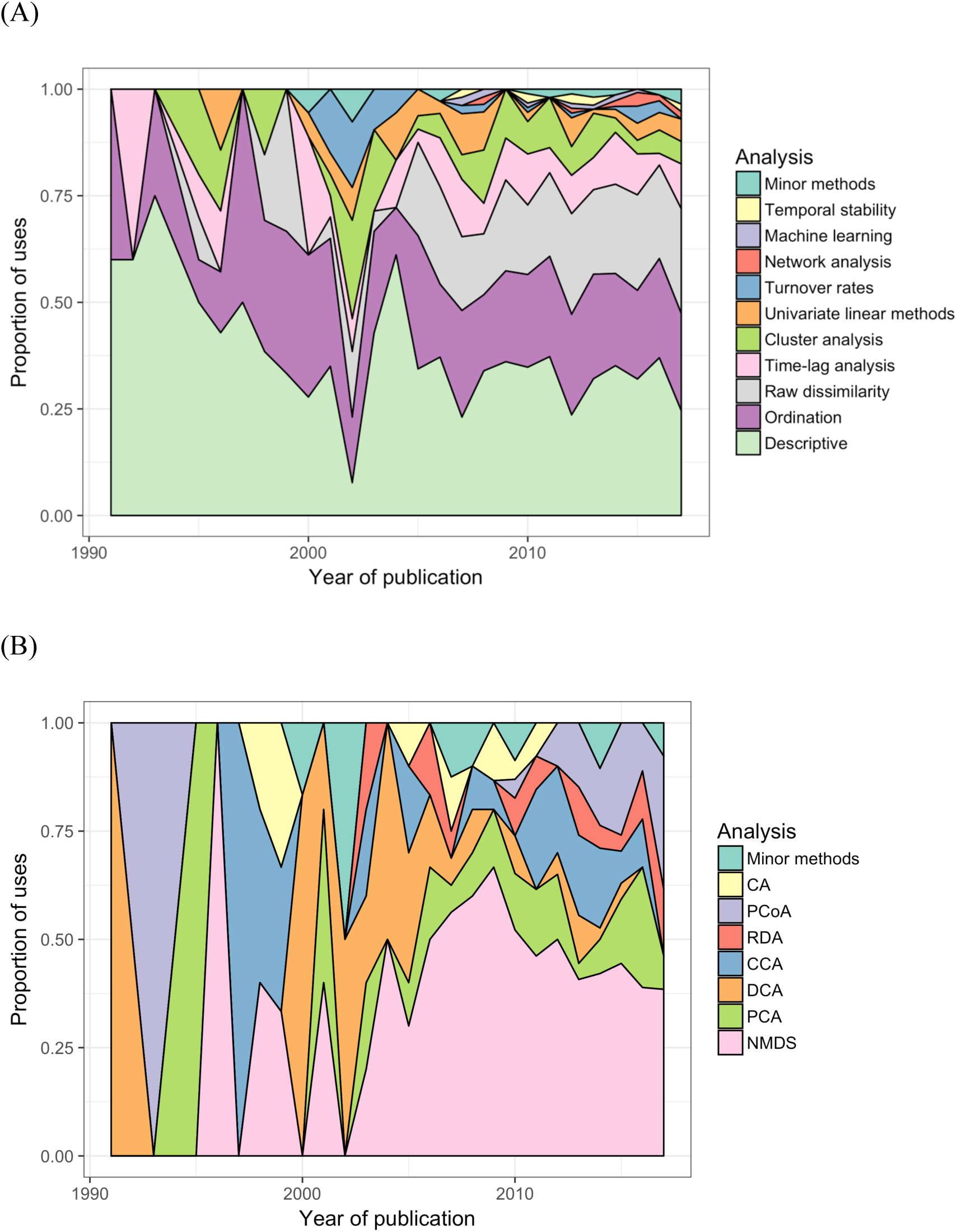
(A) The proportion of uses of each type of analysis across all sampled publications between 1990 and 2017. ‘Minor methods’ include compositional pivot days, frequency change, Markov chain modelling, multiplicative change, Moran’s eigenvector maps (MEMs), Nestedness analysis, and synchrony. (B) The popularity of the different ordination methods used across the 27 years. ‘Minor methods’ are: dbRDA, DCCA, pCCA, pRDA, Procrustes, RA and multiple co-inertia analysis. Descriptions of analysis types are detailed the S1 Text supplementary material.

**Figure S4.**
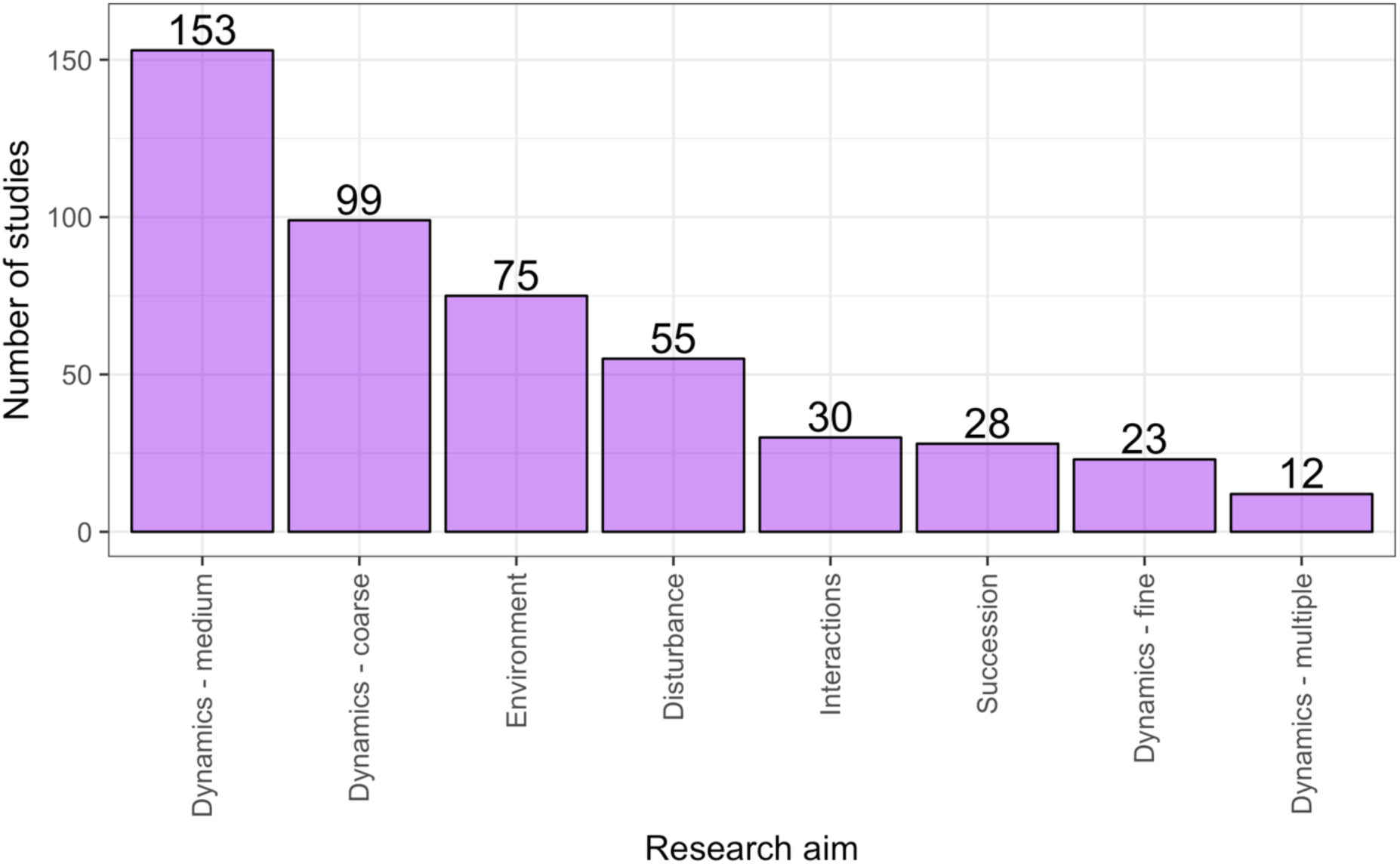
The distribution of studies across the different research aims. Research aims vary from understanding compositional dynamics at fine, medium and coarse temporal scales, to investigating the effect on species composition over time of species interactions, environmental conditions, and ecological disturbance, to understanding successional change in communities. Scales of temporal dynamics were classified as fine (days), medium (weeks, months, or seasons), and coarse (annual, years, or decades).

## Appendix S2. Other methods encountered in the literature review, not explained in the text

### Univariate linear methods

Univariate linear methods, including analysis of variance (ANOVA), analysis of covariance (ANCOVA), repeated measures ANOVA (rmANOVA), simple correlation (e.g., Pearson’s and Spearman’s correlation coefficients), and regression (linear regression, generalised linear models, generalised linear mixed effects models, logistic regression, *etc*.), are used to examine temporal changes in a univariate response (dependent) variable. The independent variable is either time, or an environmental variable that changes over time, and the response variable usually is the (relative) abundance of a focal taxon or functional group.

Univariate linear methods are applied most frequently to community dynamics by constructing separate models for each species. These methods of assessing compositional dynamics are limited because multiple analyses, conducted in parallel on selected taxa, are required to make inferences regarding composition. For even modest numbers of taxa, these multiple analyses can become unwieldy and require *P*-value corrections for multiple tests (Benjamini and Hochberg 1995, Zar 1996). For example, Bugalho et al. (2013) used repeated-measures mixed-effects modelling to assess the effects of deer exclusion over time on the abundance of seedlings and saplings of forest tree species. In this case, the univariate modelling approach was usefully applied because fewer than ten species were analysed. Alternatively, abundance can be analysed for all species in the same model as long as there are enough degrees of freedom in the model to include species as a fixed or random effect term (e.g., Buckley and Freckleton 2010); see Harrell (2001) for a rule of thumb stating that we should have at least ten independent samples for each predictor variable in our model. Generalised linear models are more flexible in that they can handle designs with unbalanced data and nested sampling (e.g., plots within sites) (Bolker et al. 2009, Zuur et al. 2009). R code is provided in Appendix S4.

### Cluster analysis (a dissimilarity based method)

Cluster analysis is a catch-all term applied to dissimilarity-based methods that either group samples together (agglomerative methods) or split all samples into sub-groups (divisive methods); the clustering is based on the distances between clusters (Legendre and Legendre 2012). There are two approaches for using cluster analysis to assess community dynamics: identifications of transitions between clusters over time, and (2) examination of differences of clusters of repeated observations. In the first approach, a separate cluster analysis is computed for each temporal sample of community data (e.g., Day and Buckley 2013). In the second approach, all samples taken at different times are included in a single cluster analysis (e.g., Araujo et al. 2008). If compositional shifts are small relative to the total compositional variation in the dataset, the first approach will not be able to detect temporal differences. However, if compositional shifts are relatively large, it yields a clear way of showing shifts in ‘community types’: groups of samples of similar composition that can be characterised using a method such as indicator species analysis (e.g., Day and Buckley 2013). The first method can potentially be applied to a larger number of time points if all that is desired is to report the percent of sites that changed cluster groups each time. For example, cluster analysis of 34 phytoplankton species in Laoshan Bay, China revealed distinct groups of species occurred at different times over a six-month period (Xu et al. 2010).

### Temporal Stability (a.k.a. coefficient of variation)

The coefficient of variation (CV) is used to measure temporal stability of the abundance or biomass of an individual taxon, or group of taxa, across all times (not space); this is not a multivariate method (Grossman et al. 1990, Brown and Lawson 2010, Hector et al. 2010). The CV is the ratio of the standard deviation to the mean, and often is multiplied by 100 to obtain a percentage. The mean of the CV values for all taxa (e.g., abundance or biomass) is used as an aggregate measure of temporal stability for a whole community; smaller values imply greater stability. Because the CV is a ratio, changes in community composition can be compared even if there are differences in total abundances among taxa or in taxon richness among taxon groups. Such values are often presented in tables or the text as a measure of variation or are sometimes used as a response variable against other (spatial) variables of interest (e.g., Doxa et al. 2012). For example, CV can be calculated for disturbed and undisturbed communities to assess if the disturbance has resulted in a more temporally variable community. Grossman et al. (1990) categorised particular values of CV to assess stability of freshwater fish communities: a community was ‘highly stable’ if it had a CV < 25%, ‘moderately stable’ if 25% < CV < 50%, ‘moderately fluctuating’ if 50% < CV < 75% or ‘highly fluctuating’ if CV > 75%. The statistical significance of CV values could be obtained by comparing the observed values to those from communities generated using randomisations under sensible null models. Note that temporal CV is necessarily be measured on a per site basis to exclude spatial variation.

### Frequency change

Changes in the frequency of individual taxa within a community over time can be assessed by subtracting their frequency at each sampling occasion from a prior sampling occasion; this is not a multivariate method. Permutation tests can be used to test the significance of these changes in frequencies (Kapfer et al. 2011). Community samples (with their given taxon frequencies) are randomised among sampling times and the change in taxon frequency recalculated with each permutation. A significant change in the observed taxon frequency is inferred if the observed change is greater than the 95^th^ percentile change from the permutations. Permutations should be spatially restricted so that only samples from homogeneous habitat types are compared; otherwise the chance of spuriously detecting change is greatly increased. The outcome of these tests is a list of taxa that had significant increases or decreases in frequencies, and the amount, between sampling times. For example, Kapfer et al. (2011) used permutation tests to assess changes in frequencies of 85 plant species from two time points 50 years apart. This method could be used on data from multiple sampling occasions, but would quickly become cumbersome with many temporal observations. It can only be used to compare individual taxa over time, does not apply to rare taxa (taxa need to occur in enough sites to make the calculations tractable), and results in a (potentially) large table of taxa with significant (or not) changes, therefore, it has limited applicability to taxon-rich datasets where changes in rare taxa are of interest.

### Multiplicative change

The change in percent cover of a taxon, or group of taxa, at a single site can be used to obtain a ‘growth rate’ using a linear regression equation; this is not a multivariate method (Debinski et al. 2010, Collins and Xia 2015). This rate of change can then be used in a regression or ANOVA to assess predictors of change for each species (‘multiplicative change’). For example, different sites or sets of sites, can be compared in their relative change in the percent cover of different functional groups (Debinski et al. 2010), as long as communities consist of taxa with similar ecology and life-histories, e.g., grassland plant communities in different moisture regimes. This method appears to be most useful for studies where (1) there is little temporal replication, or at least where differences among temporal replicates are not of interest, because the growth rate is calculated using only two temporal samples, and (2) few species, species groups, are of interest because a different analysis must be performed for each. However, it would be possible to calculate the change in percent cover for different time periods within the overall study, if one was interested in investigating finer-scale trends.

### Markov chain modelling

Markov chain modelling uses a series of transition probabilities (a ‘Markov chain’), which describe how the occupancy of a taxon or functional group in a site changes from time *t* to time *t* + 1 (Hill et al. 2004). The user varies these probabilities in space and/or time to create a set of different hypothesis-driven models that predict the observed occupancy. The model set can then be evaluated against the observed data using log-linear modelling and AIC model selection to determine the influential and significant processes on taxon site occupancy through time (Hill et al. 2004). Community dynamics usually are examined by comparing predicted changes in individual taxa, making the method easier to apply to communities of few taxa (or groups of taxa). Because models often are parameterised using the modelled data, dependence of samples needs to be dealt with through sampling design or post-hoc statistical analyses (Hill et al. 2004). This method is most appropriate for sessile communities where patch occupancy is easy to replicate and define (e.g., Hill et al. 2004, Pueyo and Beguería 2007). It best describes directional dynamics, such as secondary succession (e.g., Baasch et al. 2010, Jiménez-Franco et al. 2011) and it is relatively computationally difficult. However, it makes detailed, taxon-level predictions that can be useful for the analysis of experimental studies where researchers want to explore specific predictions from multiple scenarios, such as taxon extinctions (e.g., Wootton 2004) or restoration manipulations (Baasch et al. 2010).

### Machine learning and data mining methods

Machine learning is branch of computer science that deals with the development of learning algorithms that are used to explore large, multivariate datasets, and has the primary aim of generating accurate, predictive models (Carbonell et al. 1983). Common machine-learning methods include artificial neural networks and decision-tree methods which can be used with either univariate or multivariate response variables, and Bayesian versions can incorporate prior knowledge (Cutler et al. 2007, Olden et al. 2008). Whilst the field of machine learning focusses on making predictions from data, many of its methods also are used in a data-mining capacity to discern spatiotemporal patterns from complex and noisy datasets and can be used for classification, regression, clustering, and data reduction (Olden et al. 2008).

Machine-learning methods are attractive because they do not have many of the restrictive assumptions, such as linearity or data normality, that underpin many traditional statistical methods (Olden et al. 2008). Thus, machine-learning methods are becoming increasingly popular for exploratory and predictive analyses of community dynamics (e.g., Kampichler et al. 2010). For example, Vallès et al. (2014) used self-organising maps, an implementation of artificial neural networks (Kohonen 1982), to show that complex microbial communities sampled from infants’ guts formed distinct clusters at each of five sample times (based on relative abundance data), suggesting that successional processes were occurring within the first year of these infants’ lives. Sheaves et al. (2010) used a multivariate classification and regression tree analysis (De’ath 2002) to investigate spatial and temporal differences in fish assemblages; results showed that despite considerable spatial variation in community structure among different estuaries, there nonetheless were clear patterns of temporal change in species’ abundances and assemblage composition. These methods are very flexible and can handle both univariate and multivariate input datasets with large numbers of predictor variables, including space and time, allowing simultaneous investigation of spatial and temporal community dynamics. Standardised approaches for characterising different types of temporal patterns and causal processes in community datasets will hopefully emerge from further research. R code is not provided, but R packages are available (the MultivariateRandomForest package: Rahman 2017, and rpart package: Therneau and Atkinson 2018).

### Network analysis

Ecological network analysis enables the investigation of the complex associations among species, and among species and their environment, through space and time (Newman 2010, Blonder et al. 2012). This method recognises that an ecological community is a complex biological system comprised of interconnected units whose associations can be modelled mathematically using constructs such as vertices (representing taxa) and edges, which are the connections between the vertices (representing ecological interactions) (Blonder et al. 2012). In the context of investigating community dynamics, patterns of species’ associations and community memberships are most typically visualised as a topological network diagram created from taxon abundance data collected at multiple time points (*sensu* “time-aggregated networks” (Blonder et al. 2012). From the network topology, a variety of statistics can be calculated that measure variability in community composition and interactions at particular times, which can then be compared. This method is most applicable for multi-trophic communities or where interaction networks can be clearly defined for the community; it is less useful where a single trophic level is of interest and interactions are not quantified, e.g., a plant community, but see Cram et al. (2015) for an example of spatiotemporal microbial networks. In R, the package ‘timeordered’ is the easiest way to implement a network analysis of community dynamics data. R code is not provided.

